# Rescuing the Function of Missense-Mutated Tumor Suppressor *VHL* using Stabilizing Small Molecules

**DOI:** 10.64898/2026.01.02.697373

**Authors:** Mariam Ahmed Fouad, Christopher S. Parry, Sven A. Miller, Shipra Malhotra, Glenn A. Doyle, Ezra Shimabenga, Mashhura Nurilloeva, Elena Bondarenko, Grigorii V. Andrianov, Wayne Childers, Benjamin E. Blass, Petr B. Makhov, Johnathan R. Whetstine, Margie L. Clapper, Erica A. Golemis, John Karanicolas

## Abstract

Somatic mutations in the *VHL* gene, coupled with *VHL* loss of heterozygosity, drive sporadic clear cell renal cell carcinoma (ccRCC). From structural considerations, we hypothesized that certain mutations in the *VHL* gene thermodynamically destabilize the folded protein product; these mutations would increase the ratio of unfolded:folded pVHL, and thus cause loss of function. To test this, we used computational structure-based screening followed by biophysical characterization and cellular assays to identify small molecules that bind to the folded (native) conformation of pVHL and stabilize it. These studies led to creation of an agent, CP4.29, that stabilizes the native folded structure of mutant pVHL: this in turn restores wild-type pVHL activities to cells that harbor mutant *VHL*. These compounds may serve as starting points for further development into an unprecedented new class of kidney cancer drugs. The approach described herein may also serve as a blueprint for developing agents to correct destabilized mutations underlying other human diseases.

**Highlights:**

- Designed small molecules that re-fold and re-activate destabilized mutant *VHL*
- Lead agent re-folds and re-activates multiple common *VHL* mutants
- This approach may apply to other destabilized proteins that underlie human disease

## Introduction

Biallelic loss of tumor suppressor genes (and their associated protein products) represents a key factor in cancer development and progression [1–3]. Germline or somatic mutations in one such tumor suppressor gene, *VHL*, lead to development of von Hippel-Lindau Disease (VHLD) and clear cell renal cell carcinoma (ccRCC), respectively [4–7]. Decades of research have revealed that *VHL*’s protein product, pVHL, serves as an adaptor protein in diverse protein complexes that regulate numerous cellular functions [7–9].

The pVHL / Elongin C / Elongin B (“VCB”) complex is the most well studied of these protein complexes. The VCB complex is part of an E3 ubiquitin ligase complex that induces polyubiquitination and proteasomal degradation of substrates bearing a hydroxyproline mark, most notably members of the Hypoxia Inducible Factor Alpha (HIF-α) family [10–13]. Inactivating mutations in *VHL* thus lead to aberrant accumulation of HIF-1α and HIF-2α, which in turn induce transcription of hypoxic response genes including *VEGF-A*, *CCND1* and many more, that collectively drive angiogenesis, cellular metabolism, and cell survival [7,14]. In addition to regulating HIF, pVHL also has many important HIF-independent activities: these include ubiquitinating substrates such as ZHX2 [15], nuclear clusterin [16], IKKβ [17], AURKA [18], RPB1 [19,20], and atypical protein kinase C [21]. pVHL has also been implicated in extracellular matrix assembly, through direct interactions with collagen IV [22,23] and fibronectin-1 [24].

Mechanistic understanding of *VHL* signaling has contributed to targeted therapy for VHLD and ccRCC patients. Dramatic clinical successes came first through inhibition of HIF’s downstream effector VEGFR [25–27], and then by inhibiting HIF-2α itself [28–31]. Nonetheless, the fact that *VHL*’s rich tumor suppressive activities clearly extend beyond HIF [15,32–36] suggests that *direct* pharmacological restoration of *VHL* activity may represent a more robust and promising approach than simply targeting individual downstream effectors such as HIF. Attempts to restore the cellular activity of missense *VHL* mutants have been reported [32,37,38], but none of these yielded suitably drug-like hits to serve as a compelling starting point for further translational advancement.

In some kidney cancers *VHL* is inactivated indirectly, through epigenetic silencing or through mutations to accessory proteins such as Elongin C (*TCEB1*) [39]. Among somatic *VHL* missense mutations in kidney cancer, the Catalog of Somatic Mutations in Cancer (COSMIC) database [40] shows that mutation sites are distributed broadly throughout the protein sequence (**Figure 1a**). Despite the extremely high overall prevalence of *VHL* mutations in in kidney cancer, there are no “hotspot” positions with very high mutation frequency: rather, many different sites are mutated with low frequencies for each. Mapping these positions onto a high resolution crystal structure of the VCB complex (pVHL / Elongin C / Elongin B) bound to a hydroxyprolinated peptide from HIF [41,42] (**Figure 1a**), about 59% of kidney cancers have *VHL* mutations at sites located in regions that interact with either the Elongins or with the HIF peptide. Importantly though, the other 41% of *VHL* missense mutations in human samples do not localize to surface residues involved in protein interactions, but rather correspond to buried residues comprising the hydrophobic core of the folded protein.

**Figure 1:**
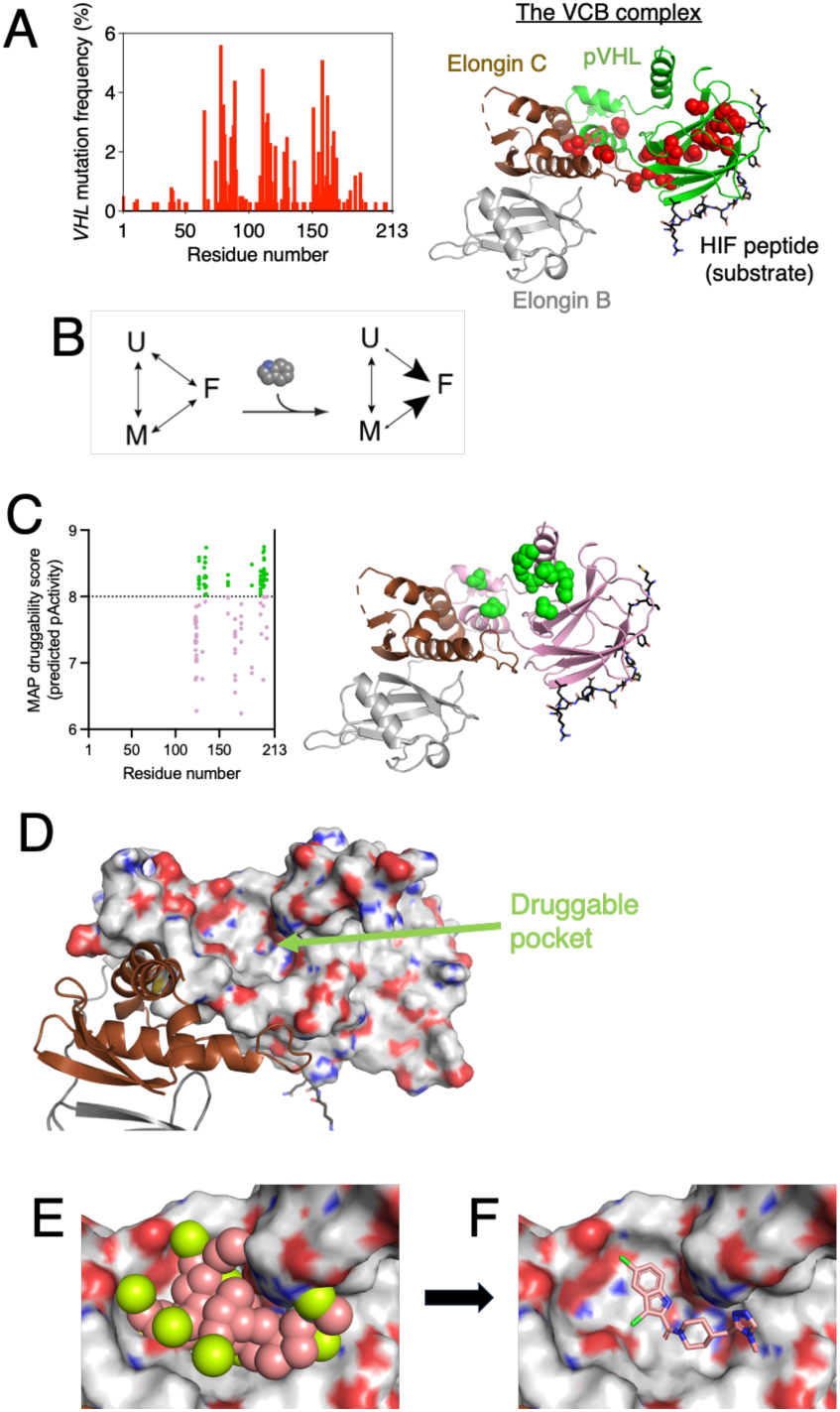
Designing pharmacological pVHL refolders by targeting a distinct surface pocket. (A) Missense mutations observed in kidney cancer are broadly distributed across the VHL sequence, and mapping them to the protein structure (PDB 1LM8) shows that many of the 20 most frequently occurring mutations (*red*) are located in the hydrophobic core of the folded protein structure. (B) Overarching design strategy for re-folders / re-activators. In principle, ligands that preferentially bind to the folded/active conformation of a protein (F) will shift the conformation equilibrium towards this state, at the expense of unfolded (U) or misfolded (M) states. (C) The MAP druggability score is intended to provide a measure of how potently some (yet unknown) ligand might engage a given protein surface. With respect to pVHL, multiple simulations implicated the same surface pockets as having the highest MAP score. Spheres in this figure (*green*) indicate locations of residues with the highest MAP score. (D) The location of this druggable pocket is distinct from the binding surfaces of the Elongins, and of the HIF recognition site. (E) Building an exemplar indicates how an ideal ligand would engage this surface pocket. Spheres indicate volume that would be filled by an ideal ligand, marked by lipophilic (*pink*) or polar atoms (*yellow*). (F) Using this exemplar as a template, virtual screening provides models of specific ligands that complement this surface pocket.

Consistent with the hypothesis that its marginal stability underlies the loss of function associated with single missense mutations in the hydrophobic core, prior studies also found that isolated (WT) pVHL is not fully folded but rather occupies a molten globule state that is particularly susceptible to aggregation [43]. Stability of pVHL in cells is dependent on the presence of Elongins B and C [44], and indeed crystallographic studies of pVHL inevitably also use the VCB complex [41,42]. Recognizing that certain disease-associated *VHL* mutations affect protein stability [45–47], a recent machine learning model employed protein stability predictions to assess the risk of ccRCC associated with a given *VHL* mutation [48]. Collectively, these observations suggest that small molecules binding to the wild-type conformation of mutant pVHL in cells ought to enhance the protein’s thermodynamic stability, thus reversing the destabilizing effect of the mutation (**Figure 1b**).

Here, we report a computational structure-based screening approach and initial optimization leading to a chemical probe (CP4.29) that directly engages the VCB complex and functionally restores *VHL* activity to different mutants in assorted ccRCC cell lines. This represents the first drug-like agent that directly restores stability and activity to mutant pVHL. We expect that this compound will serve as an entry point for developing pVHL refolders as an entirely new class of therapeutics for ccRCC and VHLD.

## Results

### Computational design of pVHL binders

In principle, binding of a small molecule to the transiently folded conformation (F) of mutant pVHL would stabilize this conformation, thus shifting the conformational equilibrium away from misfolded (M) or unfolded (U) states (**Figure 1b**). The potential binding site for such a ligand is not known *a priori*; however, targeting some non-functional region of the protein surface would be preferred, to avoid competitive inhibition with pVHL’s endogenous interaction partners. Rather than simply search for druggable pockets on the surface of the pVHL crystal structure, we expanded the potential binding sites we considered by additionally evaluating cryptic pockets transiently sampled on the protein surface [49,50]. To this end, we used a biased conformational sampling strategy [50,51] implemented in the Rosetta software suite [52] to generate low-energy pocket-containing conformations of pVHL. To ensure that formation of a cryptic pocket would not allosterically inhibit pVHL’s interaction with either the Elongins or its substrate(s), we carried out these simulations in the context of the (wild-type) pVHL / Elongin C / Elongin B / HIF-1α complex.

We carried out a focused set of individual simulations on each of pVHL’s surface-exposed residues (see Supplemental Methods). For each pocket observed in these simulations, we applied a gradient boosting machine learning model (GBM) to predict the “Maximum Achievable Potency” (MAP) value for each surface pocket [53–55]. Briefly, this model was developed by extracting pockets from a diverse collection of experimentally characterized protein-ligand complexes, then discarding the ligand and training the model to predict the ligand’s potency using only features from the protein pocket (primarily hydrophobicity and concavity). By applying this analysis to pockets from simulations of the pVHL surface, we identified a collection of pockets predicted to enable development of a ligand with potency in the 1-10 nM range (MAP values between 8 and 9) (**Figure 1c**). Mapping the location of these pockets to the protein structure revealed that each of these simulations had identified small structural variations of a large pocket already evident from the crystal structure of the VCB complex (**Figure 1d**). In principle, conformational variations of this pocket could allow for discovery of different and complementary collections of ligands. Accordingly, we advanced each low-energy conformation with a highly druggable pocket (MAP>8) for virtual screening.

For each protein conformation, we built an “exemplar” [56]: essentially a receptor-based pharmacophore that defines the shape and relative positioning of hydrogen bond donors/acceptors for a ligand that would optimally complement the protein surface (**Figure 1e**). We then used these exemplars to carry out a two-step virtual screen of chemical vendor Enamine’s collection of make-on-demand compounds. In the first phase, we screened a set of 12.5 million diverse compounds to identify 100 chemical scaffolds complementary to this region of the protein surface. In the second phase, we identified 100 analogs of each initial scaffold and combined these into a target-focused library of 10,000 compounds. We aligned each of these onto the parent scaffold, used Rosetta to refine the protein/ligand structure, and then used the vScreenML machine learning classifier [57] to distinguish likely binders from likely computational artifacts. The *Supporting Information* contains complete details of this computational protocol, including a summary of the workflow (**Figure S1**). complementarity to this surface of pVHL, both in terms of shape and hydrogen bonding groups (**Figure 1f**). The top 18 compounds based on vScreenML score were selected (**Table S1**), and these were requisitioned via Enamine’s on-demand synthesis.

### CP4 directly engages the VCB complex consistent with the intended binding mode

Having acquired 18 initial compounds from this virtual screen, we first sought to assess each for binding to purified (WT) pVHL. As noted earlier, pVHL is not stable in the absence of Elongins B and C [44], and accordingly studies of purified pVHL use the VCB complex [41,42]. For these reasons, we elected to test our initial designed hits using label-free binding assays that could be applied in the context of the VCB complex, rather than pVHL alone. Further, since our compounds are intended to bind to the folded conformation of pVHL, we elected to use WT pVHL for primary testing rather than a destabilized mutant.

We therefore purified the recombinant (WT) pVHL / Elongin C / Elongin B (“VCB”) protein complex, and we evaluated the 18 hit compounds via Saturation Transfer Difference (STD) NMR. Among the compounds showing evidence of binding was CP4 (**Figure 2a**). To fully confirm direct binding of CP4 to the VCB complex, we evaluated binding using three separate and orthogonal ligand-observed NMR assays: STD-NMR, WaterLOGSY, and CPMG [58,59]. Parenthetically, all three NMR techniques have already been used in the context of fragment screening against the VCB complex [60,61], albeit with the intention of designing new warheads for targeted protein degradation (e.g. PROTACs) to replace the widely used HIF-α peptidomimetic VH298 [62,63].

**Figure 2:**
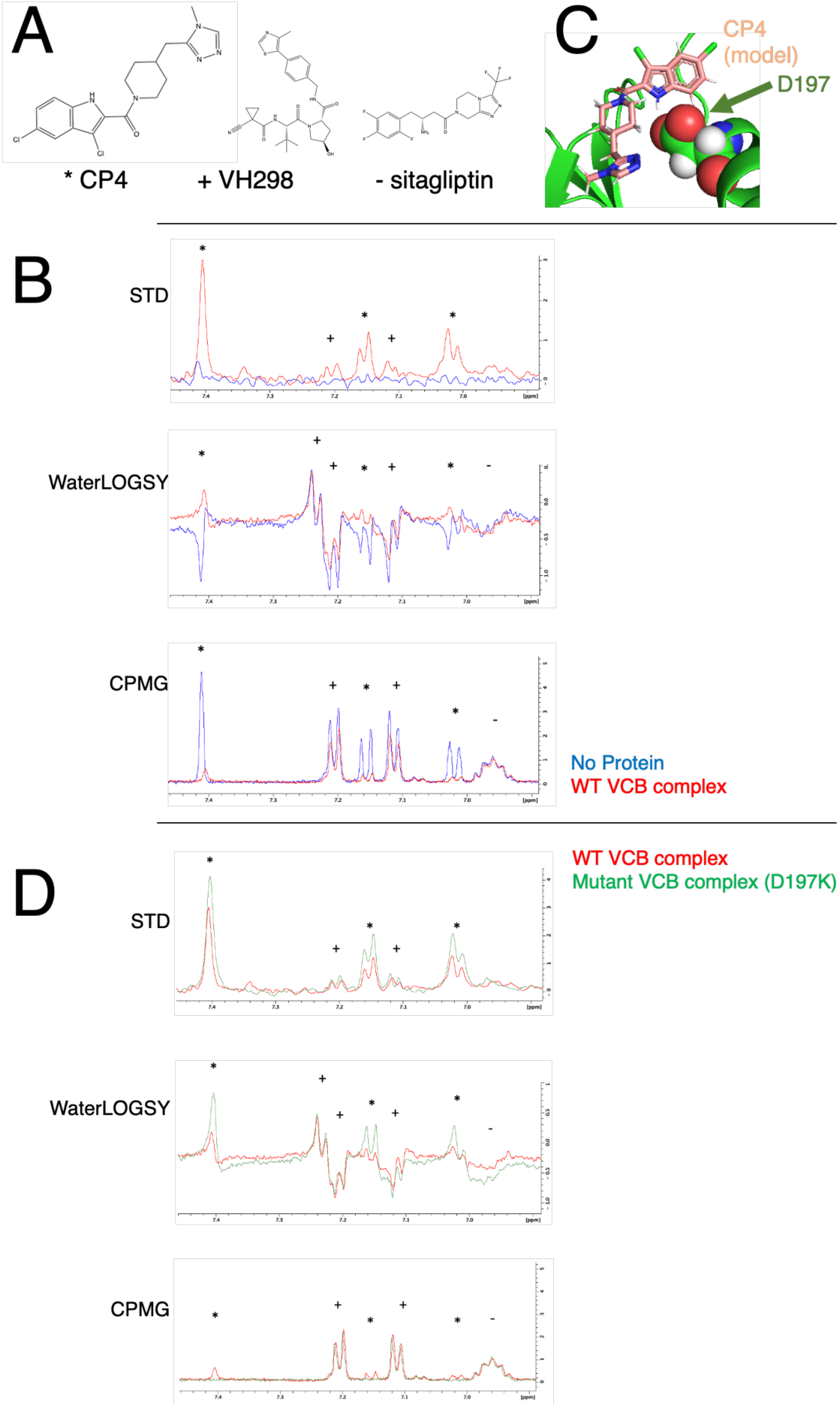
Direct evidence of CP4 binding to the VCB complex. **(A)** Chemical structure of CP4, as well as those of VH298 (positive control) and sitagliptin (negative control). **(B)** Orthogonal ligand-observed NMR spectra for three separate experiments: STD-NMR, WaterLOGSY, and CPMG. Each sample contains a mixture of CP4, sitagliptin, and VH298 in the presence (*red*) or absence (*blue*) of the VCB complex. Peaks are labeled based on whether they belong to CP4 (*), VH298 (+), or sitagliptin (-). **(C)** Based on our model of CP4 engaging the VCB complex, we elected to mutate *VHL* Asp197, a residue in the intended binding pocket. **(D)** The same three ligand-observed NMR assays were carried out using VCB complex harboring designed *VHL* mutant D197K. This mutant complex does not show any change in binding for VH298 or sitagliptin (relative to the WT VCB complex), but it *does* show a difference in binding for CP4. Due to the kinetic nature of these experiments, they cannot be used to differentiate between enhanced versus diminished CP4 binding. Full spectra from these experiments are included as **Figures S5-S10**.

An advantage of ligand-observed NMR assays is that multiple compounds can be tested in a single tube, provided that their ^1^H spectra are non-overlapping. As a positive control in these assays, we therefore included the above-referenced HIF-α peptidomimetic VH298 (**Figure 2a**). As a negative control, we sought to include a readily available compound with physicochemical properties matched to those of CP4: this led us to select the anti-diabetes drug sitagliptin (**Figure 2a**). By comparing with individual spectra for each compound, the peaks in this mixture could be unambiguously assigned as belonging either to CP4, to sitagliptin, or to VH298 (**Figure S2-S4**).

In the STD-NMR experiment, addition of VCB complex to the mixture yielded clear saturation transfer differences for peaks in the aromatic region corresponding to CP4 and VH298 (indicating binding), but not for sitagliptin (**Figure 2b**, *top*). Importantly, none of the aromatic protons show appreciable STD signal in the absence of protein: this rules out direct irradiation of these protons as an artifactual source of the STD signal [64]. Additionally, the observation of STD signal for protons from positive control compound VH298 also confirm that (WT) VHL is correctly folded in the sample, making it unlikely that CP4 is interacting non-specifically with some unfolded/misfolded form of the protein.

In the WaterLOGSY experiment, all non-exchangeable protons on the small molecules phase down in the absence of protein, as expected; the only peaks phased upward correspond to a doublet at 7.24 ppm from an exchangeable proton on VH298. Upon addition of the VCB complex, CP4 peaks flip from phased down to phased up, consistent with an interaction between the two (**Figure 2b**, *middle*). While peaks from VH298 do not change phase, the magnitude of their downward phase is diminished by the interaction with VCB; by contrast, peaks from negative control sitagliptin are unaffected.

In contrast to the first two experiments, which probe effects of magnetization transfer (either from the protein itself or bulk water), the CPMG experiment probes differences in T2 relaxation time associated with binding to a macromolecule [65,66]. Here too, peaks corresponding to CP4 exhibit dramatic loss of intensity upon addition of the VCB complex, consistent with binding (**Figure 2b**, *bottom*). The peaks from positive control VH298 also lose intensity, whereas peaks from negative control sitagliptin do not. Collectively then, all three ligand-observed NMR experiments are consistent in their demonstration of a direct interaction between CP4 and the VCB complex.

To determine whether CP4 indeed engages the intended pocket on the surface of VHL, we designed a mutation at the base of the binding site, D197K (**Figure 2c**). As noted earlier, HIF-α and its peptidomimetic VH298 both bind at a site that is distant from our intended binding surface target pocket, leading us to expect that this mutation would not affect binding of VH298 to the protein complex. Upon expressing and purifying the V_D197K_CB protein complex, we applied the same suite of NMR experiments described above.

In all three experiments, VH298 binding was indeed unaffected by the mutation: this observation also confirms that the D197K mutation did not affect foldedness of the protein complex. Conversely, all three experiments show differences in CP4 binding (**Figure 2d**), consistent with CP4 binding at the intended site from our computational designs.

We note that the magnitude of each probed effect is greater for the mutant complex V_D197K_CB than for WT VCB. A key limitation of ligand-observed NMR methods is that evidence of binding diminishes when the binding affinity is *either* too weak or strong to be observed via this class of methods [67,68]. The observed signal these techniques scales with the number of binding/unbinding events, because these events are responsible for observed signal that underlies all three methods. For this reason, changes to the protein structure that lead to faster off-rate (i.e. weaker affinity) would result in more binding/unbinding events, and thus greater signal as detected by these methods. These methods are best suited for determination of binding affinities in the 10 μM to 1 mM range, with observed signal diminishing as binding affinity is outside this range in either direction.

Ultimately, we hypothesize that the D197K mutation indeed weakens the VHL’s interaction with CP4, leading to faster off-rate (and thus more binding/unbinding events in the NMR timescale). We can rule out the fact that the mutation affects the amount of folded VHL, because the mutation does not affect the signal from VH298. Regardless of the specific effect on binding affinity, however, these data strongly corroborate the hypothesis that CP4 binds to the VCB complex near pVHL’s Asp197, as designed in the computational model. Alternative biophysical methods to directly probe binding / protein stability are confounded by the fact that pVHL is not readily purified without the Elongins present, and such characterization will therefore be deferred for a future study.

### CP4 stabilizes mutant pVHL in ccRCC cell lines

Having observed CP4’s direct engagement of purified WT pVHL, we next sought to evaluate the effect of CP4 on mutant pVHL in a more relevant cellular context. Accordingly, we obtained three different cell lines from human ccRCC that each harbor a different *VHL* missense mutation: RCC-MF (*VHL* P86S), RCC4 (*VHL* S65W), and 769-P (*VHL* I180N). Each mutation is at a buried/interfacial position in the VCB complex, but they map to different parts of the pVHL structure: P86S is in the central core, S65W is near the HIF-α binding site, and I180N is near the Elongin C binding surface (**Figure 3a**). The latter mutation is of particular interest, because mutations in the interfacial region between pVHL and the Elongins have been shown to disrupt both pVHL and Elongin stability, suggesting that each component of the VCB complex is dependent on the others for cellular stability [44]. In fact, the 769-P cell line is sometimes (misleadingly) described as “VHL null” because of the extremely low abundance of mutant pVHL [69,70]. Importantly though, none of these mutation sites are close to the CP4 binding site, allowing us to test the hypothesis that ligand binding can rescue the effect of a mutation at a distant site.

**Figure 3:**
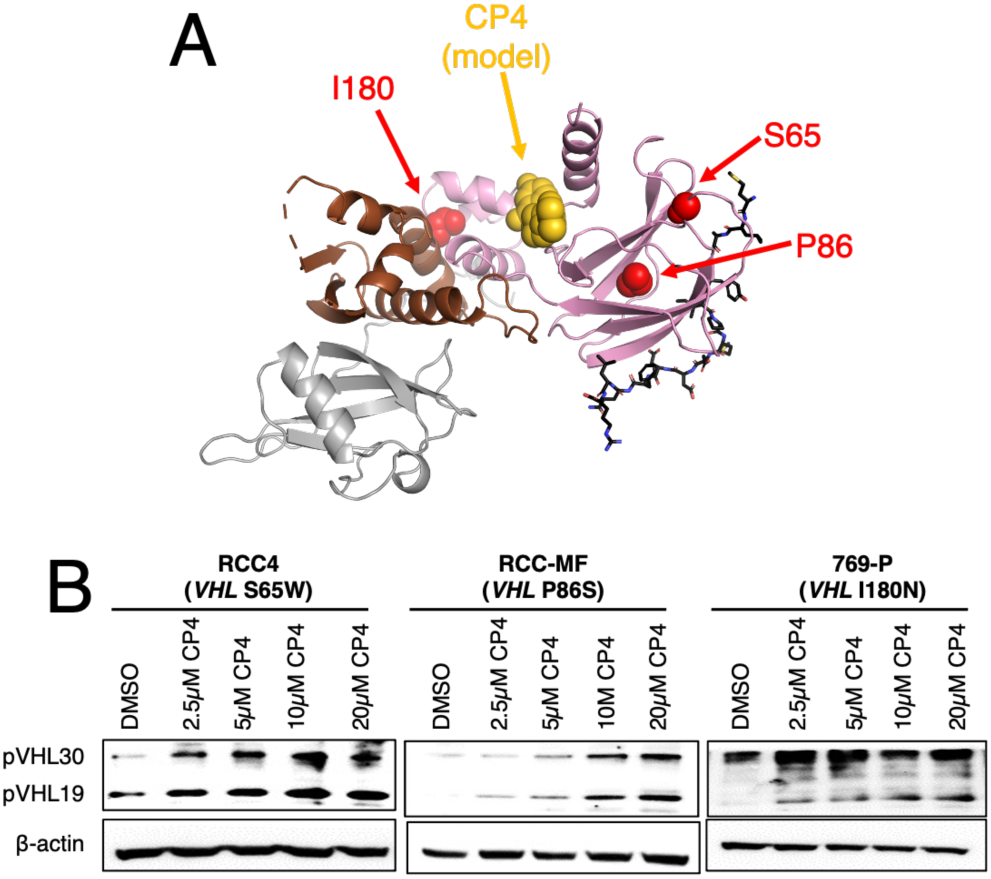
CP4 stabilizes pVHL in ccRCC cell lines harboring *VHL* missense mutations. (A) Our cellular studies use three cell lines that each harbor a different *VHL* missense mutations. The sites of these three mutations are structurally remote: P86 is in the central core, S65 is near the HIF-α binding site, and I180 is near the Elongin C binding surface. None of these positions are in direct contact with the CP4 designed binding site. (B) CP4 treatment for 1 hour leads to accumulation of both isoforms of mutant pVHL.

Many *VHL* mutants have been found at lower cellular abundance than the WT protein, due to shortened cellular half-life [38]. Accordingly, we hypothesized that treatment with CP4 would increase cellular abundance of mutant pVHL by extending its half-life. To test this, we treated each cell line with an increasing concentration of CP4 for 1 hour, and we probed for pVHL abundance via Western blot. Cells produce pVHL in two isoforms, pVHL19 and pVHL30. pVHL19 arises due to an alternative initiation site at Met54, thus resulting in a protein product that lacks 53 residues in the disordered N-terminus of pVHL30. Despite slightly different subcellular localization, the shared folded structure of these isoforms allow both to bind Elongins and recognize substrates including HIF-α [71]. We find that treatment with CP4 leads to accumulation of both pVHL isoforms, in all three cell lines (**Figure 3b**).

### CP4.29 induces VHL-dependent depletion of HIF-2α

The most relevant HIF isoform in the context of ccRCC is HIF-2α [72,73]. Given that CP4 induces refolding of mutant pVHL in cells, we expected that CP4 treatment would also restore pVHL activity in cells – most notably including the ubiquitin-mediated destruction of HIF-2α. Initial experiments evaluating CP4 in this context showed little to no reduction in HIF-2α, however, despite CP4 stabilizing mutant pVHL isoforms in the same cell lines.

In past studies using an artificial system, we screened small molecules to evaluate their complementation of a designed cavity introduced by mutation into an enzyme [74]. We found that certain ligands bound to this cavity but led to slight distortions in the protein structure, such that enzyme activity was not rescued. In fact, restoring enzyme activity required that the ligand bind to the designed cavity in a manner that precisely recovered the original enzyme conformation [75]. Thus, close analogs could have dramatic differences in their rescue of enzyme activity, that were not simply attributable to differences in binding affinity [74].

We therefore surmised that CP4 may bind to mutant pVHL such that the resulting conformation is very close to that of the WT protein, but not precisely so. While CP4 itself did not recover HIF-2α degradation by mutant pVHL, we hypothesized that a close chemical analog of CP4 might do so. Accordingly, we relied on the straightforward synthetic accessibility of CP4 (**Figure 4a**) to prepare a library of close analogs, and then supplemented this collection using additional analogs available from Enamine for a total of 46 compounds (**Table S2**).

**Figure 4:**
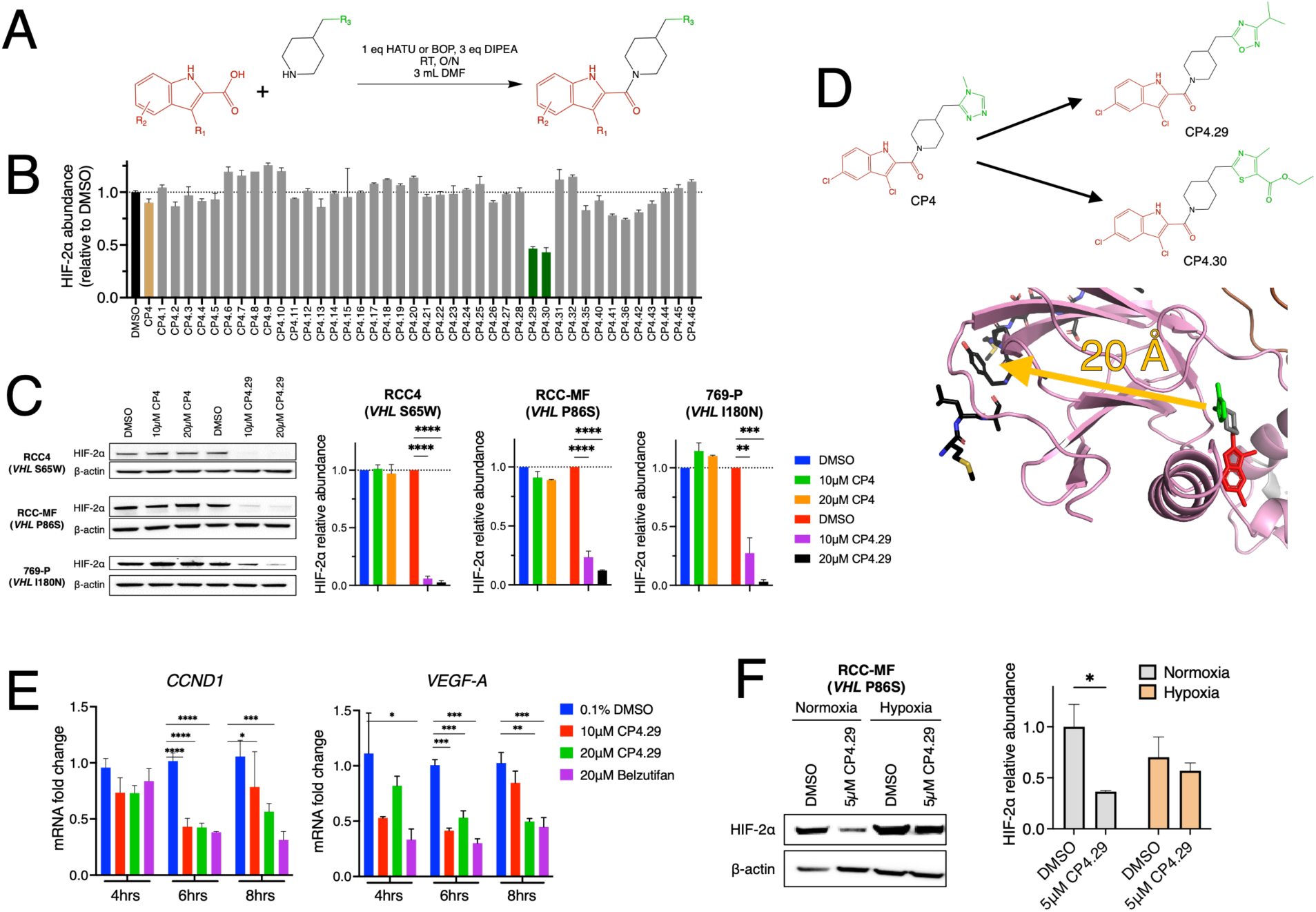
CP4.29 degrades HIF-2α by re-activating mutant pVHL in ccRCC cell lines. **(A)** Reaction scheme for preparing CP4 derivatives. **(B)** CP4 derivatives were screened at 20 μM in RCC-MF cells, by using ELISA to probe HIF-2α abundance after 2 hour treatment with each compound. **(C)** Treatment with CP4.29 for 2 hours led to depletion of HIF-2α in RCC4, RCC-MF, and 769-P cell lines. **(D)** CP4 and CP4.29 differ only slightly in chemical structure, at a site approximately 20 Å from substrate HIF-α in our model of target engagement. **(E)** Application of CP4.29 to 769-P cells reduced mRNA levels of HIF-2α target genes *CCND1* and *VEGF-A*. **(F)** CP4.29 treatment for 4 hours induces HIF-2α degradation in RCC-MF cells under normoxic conditions (20% O2) but not hypoxic conditions (2% O2), consistent with pVHL-mediated recognition of HIF-2α.

We evaluated this panel in RCC-MF cells (*VHL* P86S), by applying a 2 hour treatment of each compound at 20 μM, then probing for HIF-2α by ELISA. Whereas most CP4 derivatives (and CP4 itself) did not lead to any appreciable difference in HIF-2α, related compounds CP4.29 and CP4.30 induced a dramatic reduction of HIF-2α (**Figure 4b**). To test whether a single compound could recover activity of multiple *VHL* mutants, we applied CP4.29 to all three cell lines, and we observed depletion of both HIF-2α (**Figure 4c**) and HIF-1α (**Figure S11**) in all three cell lines.

The distinction between CP4 and CP4.29 is remarkably modest. Relative to CP4, derivative CP4.29 replaces a 4-methyltriazole with a 3-isopropylozadiazole. While we cannot rule out subtle differences in the compounds’ physical properties that may have led to enhanced cell permeability, the fact that other substitutions did not confer this activity implies that this is not the case (**Table S2**). In our model of CP4 bound to the VCB complex, this substituent that differentiates CP4.29 and CP4.30 from CP4 is in direct contact with a long loop that connects the two halves of pVHL’s β-sandwich, about 20 Å from the bound HIF-α peptide (**Figure 4d**). The surprising impact of these specific substitutions suggests an unexpected allosteric coupling between the two sides of the β-sandwich, such that subtle structural differences on one side can impact binding at the other side. Through evaluation of further CP4.29 analogs (**Table S3**), we identified alternate substituents with similar activity to CP4.29. Changes to the dichloroindole group inevitably led to loss of activity, suggesting that this side of the ligand is already well-optimized for binding to pVHL.

A key consequence of HIF-α destruction is the expected downregulation of HIF target genes. Due to the role of HIF-α in transcriptional activation, the principal activity of HIF-2α antagonists (including approved drug belzutifan) is to inhibit expression of HIF-2α dependent genes such as *CCND1* (cyclin D) and *VEGF-A* (VEGF-A) [76]. Using the 769-P cell line that contained very little pVHL in the absence of our stabilizers, we applied RT-PCR to probe the activity of CP4.29 on mRNA levels of both genes, using belzutifan as a positive control. Remarkably, CP4.29 showed similar activity and kinetics as belzutifan (**Figure 4e**), despite their very different mechanisms of action.

Having established CP4.29 as our most promising degrader of HIF-2α, we next sought to evaluate its mechanism of action. As noted earlier, pVHL recognition of HIF-2α depends on the latter bearing a hydroxyproline post-translational modification in response to a normoxic environment (high O_2_). Under hypoxic conditions (low O_2_), HIF-2α does not receive this modification and is not targeted by pVHL for destruction [7,10,11,14]. To test whether CP4.29 induces HIF-2α degradation in an oxygen-dependent manner, we applied 5 μM CP4.29 to RCC-MF cells that had been acclimated for 24 hours at either a normoxic environment (20% O_2_) or a hypoxic environment (2% O_2_). Consistent with a pVHL-dependent mechanism of CP4.29, we observe HIF-2α degradation only under normoxic conditions (**Figure 4f**).

### CP4.29 restores pVHL’s ubiquitin ligase activity against multiple substrates

*VHL* tumor suppressive activity derives not just from degrading HIF-α: it also induces degradation of other oncogenic substrates including Aurora kinase A (AURKA) [18] and ZHX2 [15]. Using the same 769-P cell line (*VHL* I180N), we therefore probed for depletion of these substrates and found that CP4.29 rescues these activities of mutant pVHL as well (**Figure 5a**). This observation is important because CP4.29 was selected based on its activity in rescuing HIF-2α depletion: the fact that CP4.29 also depletes other pVHL targets provides more stringent evidence that CP4.29 may be fully recapitulating the WT pVHL structure, rather than simply re-constructing a few key interactions sufficient for recruiting HIF-α.

**Figure 5:**
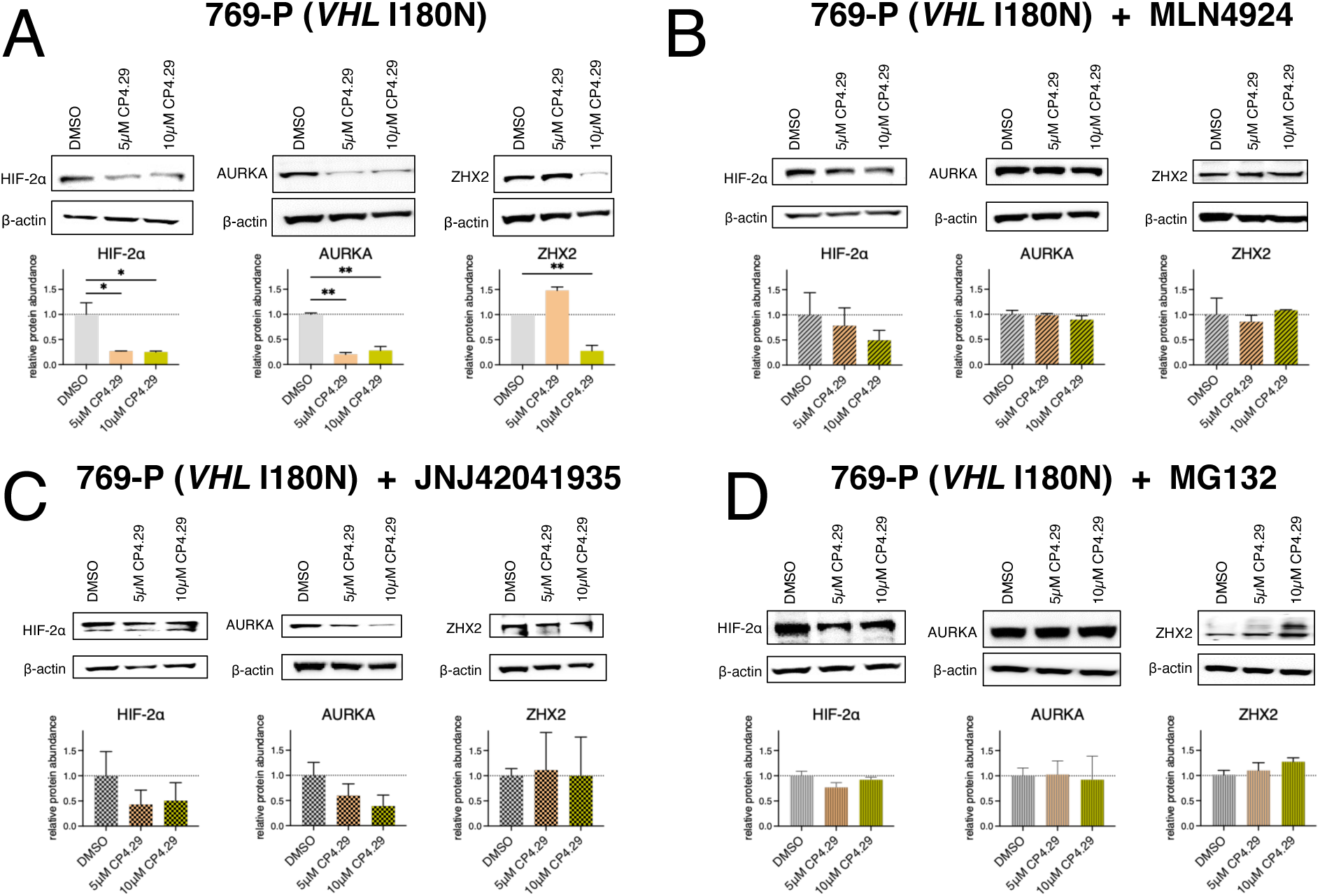
Activity of CP4.29 using mutant pVHL recapitulates the expected dependencies of recognition by WT pVHL. (A) In 769-P cells (*VHL* I180N) treatment with CP4.29 induces depletion of HIF-2α, and also depletion of additional pVHL substrates AURKA and ZHX2. Analysis of HIF-2α was obtained after 2 hr treatment, whereas analysis of AURKA and ZHX2 were obtained after 24 hr treatment. (B) CP4.29’s depletion of all three substrates is mitigated by co-treatment with 1 μM neddylation inhibitor MLN4924, demonstrating that CP4.29 activity is dependent on a cullin-RING E3 ligase. (C) CP4.29’s depletion of HIF-2α and ZHX2 (but not AURKA) is mitigated by co-treatment with 50 μM PHD inhibitor JNJ42041935, demonstrating that CP4.29 activity in this mutant *VHL* cell line recapitulates the molecular recognition patterns of WT pVHL. (D) CP4.29’s depletion of all three substrates is mitigated by co-treatment with 20 μM proteasome inhibitor MG132, consistent with a ubiquitin-mediated (proteasomal) path to their destruction=

To establish that the observed depletion of these pVHL substrates is indeed dependent on the ubiquitin ligase pathway, we applied several complementary pharmacological controls. First, we co-treated cells with CP4.29 and MLN4924, an inhibitor of NEDD8-activating enzyme (NAE) [77]. Cullin-RING E3 ligases (CRLs) require NEDD8ylation of the cullin to assemble into active complexes; thus, inhibiting the NEDD8ylation cascade using MLN4924 disrupts ubiquitination by CRLs (including pVHL). We found that treatment with MLN4924 abrogated the effect of CP4.29 (**Figure 5b**), confirming that our observed depletion of HIF-2α, AURKA, and ZHX2 is dependent on a CRL.

Second, we co-treated cells with CP4.29 and JNJ42041935, an inhibitor of prolyl hydroxylase (PHD) enzymes [78]. Hypoxia-dependent recognition of HIF-α and ZHX2 by pVHL depend on their bearing a hydroxyproline post-translational modification; this mark is added by the PHD family of enzymes. By contrast, AURKA is degraded by pVHL in a hypoxia-independent manner [18]. We found that treatment with JNJ42041935 protected hypoxia-dependent substrates HIF-2α and ZHX2 from depletion by CP4.29, but not AURKA (**Figure 5c**). Thus, this observation implies that CP4.29-mediated depletion of these three targets is consistent with the same modes of recognition used by WT pVHL.

Next, we co-treated cells with CP4.29 and MG132, a direct inhibitor of the 26S proteasome [79,80]. If depletion of these three targets is indeed due to ubiquitination by re-activated mutant pVHL, their ultimate destruction would occur at the proteasome. In the presence of MG132 to block the proteasome’s ability to degrade ubiquitinated proteins, we observe markedly less depletion of HIF-2α, AURKA, and ZHX2 (**Figure 5d**), confirming that their destruction by CP4.29 is indeed primarily through the proteasome.

To confirm that the observations from this mechanistic analysis was robust towards the *VHL* mutation / cell line selected, we applied similar pharmacological controls using the RCC4 cell line (*VHL* S65W) and obtained analogous findings (**Figures S12 and S13**). Finally, we sought to directly confirm that CP4.29-induced re-activation of mutant pVHL leads to enhanced (poly-)ubiquitination of these three targets. We used the cell lysates from the 769-P cells co-treated with MG132 (**Figure 5d**), to trap ubiquitinated proteins without allowing their degradation. We then used immunoprecipitation against each of the three pVHL substrates, followed by Western blotting against ubiquitin. We observed high molecular weight species for all three substrates upon treatment with CP4.29, but only in parental 769-P cells and not in an isogenic *VHL^-/-^* knockout line (**Figure S14 and S15**).

Taken together, these results demonstrate that CP4.29 re-activates ubiquitination patterns associated with WT *VHL*, in cells harboring mutant *VHL*. Ahead of characterizing CP4.29 through *in vivo* studies, further development of this agent will be needed. Whereas our initial optimization focused solely on achieving on-target cellular activity, the ADME properties of CP4.29 must next be improved.

Specifically, this agent is too lipophilic for immediate advancement, as evidenced by its poor solubility in the absence of DMSO and its high plasma protein binding (**Figure S16**), and affirmed by its high calculated logP value (cLogP=4.95 per ALOGPS 2.1 [81]). In addition to improving potency of pVHL re-activation into the nanomolar range, careful experimentation will also focus on probing for potential off-target activities at the physiologically relevant concentrations of next-generation pVHL re-folders.

## Discussion

Clear cell renal cell carcinoma (ccRCC) and von Hippel-Lindau disease (VHLD) represent two distinct cancer phenotypes linked by their shared molecular underpinnings: loss of *VHL* activity [4,7]. Treatment paradigms for both ccRCC and VHLD initially focused on inhibiting angiogenesis using VEGFR inhibitors, thus disabling a key downstream effector of HIF-2α [25–27]. More recently, the breakthrough drug belzutifan showed impressive clinical responses by inhibiting HIF-2α directly [28–31], thus blunting all of HIF-2α’s downstream effectors rather than a subset of them. Nonetheless, the basis for *VHL* tumor suppression extends beyond its ability to degrade HIF-2α [15,32–36]. Because of this, there remains an unmet opportunity for improved therapeutic intervention in *VHL*-mutated cancers, and perhaps even for cancer prevention in VHLD individuals who harbor germline mutations in *VHL*.

Despite pVHL having been identified as the unambiguous driver of these diseases for many years, campaigns to pharmacologically restore pVHL activity have met with limited success [32,37,38]. Challenges developing suitable assays have certainly hampered these efforts. The fact that mutant pVHL is not amenable to purification – as even the WT protein requires partners Elongin B and C for stability – diminishes opportunities to use biochemical or biophysical screening techniques. Separately, phenotypic screens can be difficult to implement due to pVHL’s many activities; further, they tend to uncover compounds that act on the *VHL* axis indirectly, such as inhibitors of histone deacetylases that modulate cellular chaperones including Hsp90 [38,82]. While manipulating cellular proteostasis may yet provide favorable clinical outcomes [83], these agents’ broad pharmacology and indirect mechanism of action leaves open concerns surrounding what other cellular changes may arise from their use.

For identifying refolder agents, then, structure-based computational design presents a natural alternative to traditional screening methods. As demonstrated in this study, computational methods can be applied to identify a suitably druggable region on the folded protein (**Figure 1**), and then to select compounds that engage this surface in a targeted manner (**Figure 2**). Many approved drugs act via allosteric mechanisms, including GPCR agonists that bind to the active receptor conformation at the (extracellular) orthosteric site and thus activate (intracellular) signaling. The refolding strategy we describe here is a select example of this general mechanism, differentiated by the fact that the unliganded (inactive) protein target resides in an unfolded conformation.

Importantly, however, an agent that binds to the target protein and induces refolding does *not* necessarily re-activate the protein’s activity. Whereas computational design provided an initial ligand CP4 to bind the intended pocket on pVHL and confer cellular stability (**Figure 3**), further development into derivative CP4.29 was necessary before pVHL activities were restored. This finding points to the inadequacy of our initial model for pVHL re-activation (**Figure 1b**): re-folding the protein target is necessary but not sufficient for re-activating it. In retrospect, this insight speaks in direct analogy to biased agonism of GPCR ligands [84,85]. In such examples, synthetic ligands can engage a receptor and impact its structure or dynamics in a manner that nearly – but not quite – recapitulates that of the natural ligand. Due to the resulting differences in the bound receptor, downstream signaling can be affected in subtle and complex ways.

In this context, a further challenge identifying re-activators of destabilized pVHL comes into focus: the “steepness” of the structure-activity relationships (SAR) surrounding CP4.29. In the case of a phenotypic screen searching for agents that induce depletion of pVHL targets, compounds such as CP4 or many of its analogs would not be recognized (**Figure 4b**). Searching at the outset for pVHL re-activators would therefore have necessitated extremely dense coverage of chemical space in an initial hit-finding campaign, due to the apparent sensitivity of re-activation to relatively subtle chemical substitutions (**Figure 4d**). Despite this apparent sensitivity, however, CP4.29 re-activates pVHL variants in which the mutation occurs at diverse locations in the protein structure (**Figure 3a**).

Analysis of sequencing data from human tumors implicated a set of 125 genes as harboring cancer driving mutations: 71 of these genes are tumor suppressors, and 54 are oncogenes [86]. Despite the abundance of well-validated tumor suppressor targets, however, nearly all targeted therapies in oncology address oncogenes rather than tumor suppressors [87]. This is a consequence of the fact that it is inherently easier to develop molecular agents that inhibit the activity of a molecular target, rather than to develop agents that re-awaken the lost activity of a mutated target; and accordingly, it remains standard practice to address tumor suppressors by targeting their downstream effectors.

As demonstrated above, both CP4.29 (pVHL re-folder) and belzutifan (HIF-2α antagonist) inhibit HIF-2α dependent gene expression (**Figure 4e**). Because of their fundamentally different mechanisms of action, however, CP4.29 also rescues pVHL’s activity on additional substrates (**Figure 5**), and thus may restore additional (HIF-2α independent) tumor suppressive activities that belzutifan does not. As with many other multi-functional tumor suppressors that can become inactivated by destabilizing mutations [88,89], this example demonstrates that direct pharmacological re-activation of a mutant tumor suppressor holds the potential for restoring *all* dysregulated activities, rather than just the subset associated with a single downstream effector.

Availability of cancer genomic data makes it tempting to speculate what fraction of ccRCC and VHLD cases harbor mutations that could be addressed by compounds such as CP4.29. Such an analysis is premature at this stage, however, because cellular rescue is confounded by a variety of factors including the extent of destabilization conferred by the mutation, the concentration of Elongin proteins (which indirectly impact pVHL stability), and more. For these reasons, future studies must also include direct evaluation of the impact of CP4.29 on a broader collection of cancer cell lines. This will allow the domain of applicability for CP4.29 to be unambiguously determined, and such studies may also guide development of analogs that expand the set of mutants which can be addressed. Ultimately, the *in vivo* applicability of future CP4.29 analogs will be critical to determine across multiple heterogeneous patient-derived samples, and ongoing developments in new preclinical models will certainly enable such efforts [90].

## Conclusions

Given the broad assortment of human diseases causally linked to protein folding defects, there has long been enthusiasm for developing “pharmacological chaperones” to reverse the effect of destabilizing pathogenic mutations [91]. Despite the simple conceptual framework by which such agents might act (**Figure 1b**), there are only four examples that have reached FDA/EMA approval; moreover, the mechanisms of action underlying these four examples are somewhat complicated by the nature of their respective targets [92,93]. Here, we describe CP4.29 as a new pharmacological agent that re-folds and re-activates mutant pVHL, which may serve as a starting point for focused drug discovery efforts.

Equally important, however, these studies define the challenges associated with identifying pharmacological re-activators for other tumor suppressors, and they demonstrate how structure-based computational design is well-positioned to address such targets.

## Methods

Detailed descriptions of computational and experimental methods are provided in the *Supporting Methods* section.

## Acknowledgements

This work made use of Fox Chase Cancer Center’s CCSG shared resources, specifically the Cell Culture Facility (CCF), Cell Sorting Facility (CSF), Genomics Facility (GF), and Spectroscopy Facility. This work was supported by grants from DOD (W81XWH-20-1-0844) and the NIH National Institute of General Medical Sciences (R01GM112736, R01GM141513, R35GM144131). This research was also funded in part through the NIH/NCI Cancer Center Support Grant P30CA006927, TUFCCC/HC Regional Comprehensive Cancer Health Disparities Partnership U54CA221705, and through Fox Chase / Moulder Center Pilot Funds. This work used the Extreme Science and Engineering Discovery Environment (XSEDE) allocation MCB130049, which is supported by National Science Foundation grant number 1548562. This work also used computational resources through allocation MCB130049 from the Advanced Cyberinfrastructure Coordination Ecosystem: Services & Support (ACCESS) program, which is supported by National Science Foundation grants 2138259, 2138286, 2138307, 2137603, and 2138296.

## Notes

### Competing Interest Statement

The authors have declared no competing interest.

### Summary of Updates

Author affiliations updated. Funding sources updated. No other changes.

