## Supplemental Materials for "Rescuing the Function of Missense-Mutated Tumor Suppressor *VHL* using Stabilizing Small Molecules"

[†]Equal author contributions.

### Supporting Methods

<u>*Details of Computational Methods*</u>

*Computationally generating pocket-containing conformations of pVHL*

The presence of a surface pocket is necessary, but not sufficient, for the druggability of a protein. Druggable low-energy pockets may or may not be evident from a crystal structure of the unbound protein [S1,2]: accordingly, it can be beneficial to explore alternate low-energy protein conformations [S3,4]. Simulations were started from the crystal structure of the VCB complex bound to a hydroxyproline containing HIF-1α peptide (the "VCBH" complex), PDB ID: 1LM8 [S5]. All usage of the Rosetta3 software suite used developer version: 7dc49c1abe6a9b8aa9c0c5dc3cc2e81968ad1198.

Prior to (biased) pocket opening simulations [S3], a set of 100 unbiased trajectories were first generated using Rosetta's "relax" application then the "minimize" application [S6,7] as follows:

```
relax.default.linuxgccrelease -s 1lm8.pdb -relax:fast -nstruct 100 -ex1 -ex2 -ex1aro
       -ignore_unrecognized_res -database <path_to_Rosetta>/database
minimize.default.linuxgccrelease -s 1lm8_00##.pdb -out:file:scorefile miniscore.sc
       -database <path_to_Rosetta>/database
```

Individual pocket opening simulations seek to find cryptic pockets in direct contact with some pre-specified "target" residue [S3]. To select all reasonable target residues for pVHL, we calculated the solvent accessible surface area (SASA) for each pVHL residue in the context of the VCBH complex as follows:

```
sasa_list_v2.default.linuxgccrelease -s 1lm8.pdb
```

A target residue was defined as any pVHL residue that 1) had no difference in SASA between the VCBH complex and pVHL alone, and 2) had SASA value > 10 $Å^2$. The first criterion in this definition rules out any residues at the interface between two protein partners (thus preventing the inadvertent design of direct inhibitors of VHL activity). The second criterion avoids any target residues that would dramatically change the protein conformation, since the residue would otherwise be completely buried.

Using the lowest-energy conformation from the unbiased simulation, a set of 100 biased simulations [S3] were initiated for each target residue as follows:

```
relax.default.linuxgccrelease -s 1lm8.pdb -relax:fast
             -nstruct 10 -ex1 -ex2 -ex1aro -ignore_unrecognized_res
             -pocket_zero_derivatives -pocket_max_spacing 12
             -pocket_psp false -pocket_sps -pocket_num_angles 2
             -score:patch pocket.wts.patch -constraints:cst_file constraint
             -pocket_grid_size 10 -database <path_to_Rosetta>/database
```

Here, “pocket.wts.patch” is a text file containing a single line reading “pocket_constraint = 1.0” and “constraint” is a text file containing a single line consisting of the strength of the biasing term (we used -0.25) followed by a space and then the target residue).

Following generation of these “pocket opened” structures, each was subjected to (unbiased) minimization as shown earlier, to enabling fair energetic comparison between unbiased and biased structures. As in earlier studies [S3], output structures from the biased simulations were discarded if they were more than 2.5 REU worse than the maximum energy from the unbiased collection.

For the remaining output structures from the biased simulations, the pocket volume at the target residue was determined as follows (where ## refers to the relaxed/minimized index number in the file name, and “#:V” target residue number (“#”) on chain V (“:V”) in the PDB file):

```
pocket_measure.default.linuxgccrelease -s 1lm8_00##_0001.pdb
       -num_angles 100 -pocket_max_spacing 12 -pocket_psp false -pocket_sps
       -central_relax_pdb_num #:V -database <path_to_Rosetta>/database
```

Any pockets that had volume of less than 150 Å$^3$ were excluded from further analysis.

*Computationally generating exemplars from pocket-containing conformations of pVHL*

For each of the pocket-containing conformations generated above, exemplars were built at the target residue pocket as described previously [S4,8], using the following command:

```
make_exemplar.default.linuxgccrelease -s 1lm8_00#_0001.pdb
        -pocket_num_angles 100 -pocket_psp -pocket_grid_size 10 -pocket_max_spacing 12
        -central_relax_pdb_num ##:V -pocket_filter_by_exemplar -pocket_static_grid
        -pocket_limit_exemplar_color -pocket_dump_exemplars -min_atoms 1 -max_atoms 65
        -database <path_to_Rosetta>/database
```

Concatenating the PDB files for the protein and an exemplar together produced a VCBH:exemplar complex, as shown in **Figure 1d**.

*Computational predictions of pocket druggability*

The "Maximum Achievable Potency" (MAP) that a ligand might obtain for a given surface pocket was predicted using a recently-developed machine learning model [S9-11]. The PDB file containing the VCBH:exemplar complex was first used to define all protein atoms within 4.0 Å of any exemplar atom, using a script provided in GitHub [S10]. Input features for the ML model were calculated using these atoms, and these served as the basis for predicting the MAP value (in units of -log(potency)).

Individual exemplars were carried forward for virtual screening only if the yielded a MAP value greater than 8 (implying that thorough optimization could yield a small molecule with Kd better than 10 nM).

*Virtual screening to identify pVHL binders*

Virtual screening was carried out in two phases: an initial screen to select chemical scaffolds complementary to the pVHL surface, and then a second screen to explore more deeply available compounds based on these scaffolds.

To prepare a suitable database for the first phase of screening, we began with Enamine's REAL Space Diverse Set, corresponding to ~15M distinct small molecules at the time. Compounds with molecular weight below 300 Da or above 550 Da were removed using OpenEye's "filter" application in Omega2 [S12-14]. Removing these compounds left a set of ~12.5M compounds available for screening.

The initial library (as SMILES strings) was split into smaller files each containing ~50,000 SMILES strings. Omega2 was then used to build conformers for the compounds in each file, as follows:

```
omega2 -in <input_file>.smi -out <output_file>.oeb.gz
       -prefix <insert_prefix_name> -warts -strictstereo false
```

Each pVHL exemplar was then used as the basis for screening this library, as described earlier [S8]. Briefly, OpenEye's ROCS software [S15,16] was used to align each conformer from the database onto the exemplar and provide a quantitative measurement of similarity for both geometric and chemical features. As in our previous studies [S8], an initial overlay was generated using FastROCS [S17] and then refined using the standard ROCS application. Prior to the second ROCS step, compounds' protonation states were adjusted using OpenEye's "fixpka" application in QUACPAC [S18] with default parameters. To allow for fair comparison between hits coming from different exemplars, aligned scores were expressed as a Z-score relative to all compounds screened against that exemplar. In contrast to previous work [S8], the TanimotoCombo score was used rather than the TverskyCombo score.

The top-scoring 20,000 matches were converted from ".oeb.gz" files to ".sdf" file format using the convert.py script in the OpenEye OEChem toolkit [S17]. From the ".sdf" file format, each was parameterized for use in Rosetta using the following command:

```
Rosetta/main/source/scripts/python/public/molfile_to_params.py
       <ligand_conformer>.sdf -p <ligand_conformer>
```

The resulting .pdb files for each of the top 20,000 hits were concatenated into the corresponding pocket-opened structure, to yield an initial docked complex. Full-atom Rosetta minimization was carried out on each protein:ligand complexes as follows:

```
minimize_ppi.default.linuxgccrelease -extra_res_fa <conformer_params_file>.params
       -s <cat_protein_ligand_structure>.pdb -database Path_to_Rosetta/main/database
```

Minimized complexes were then used as input into the vScreenML machine learning classifier [S19]. Based on vScreenML scores, the top 100 compounds were selected for further exploration of these scaffolds.

Using the Enamine REAL Space Navigator application [S20], 100 analogs for each of these initial scaffolds were identified from amongst a total of ~21 billion compounds: collectively then, these served as a new focused library of 10,000 compounds. Conformers for each of these 10,000 compounds were generated using Omega2 as described above. Following conformer generation, each conformer for a given compound was aligned onto the parent scaffold in the minimized model, using the following command in ROCS:

```
rocs -query <parent_compound_conformer_post_minimization>.pdb
       -dbase <analog_id>.oeb.gz -prefix <analog_id> -status none
       -rankby TanimotoCombo -besthits 1
```

Following the identification of the best overlapping analog conformer, the resulting conformer was converted from ".oeb.gz file" format to .sdf file format using the following OEChem [S17] command:

```
python openeye/examples/oechem/convert.py
       <top_analog_conformer>.oeb.gz <top_analog_conformer>.sdf
```

Since standard output files from ROCS contain both the initial conformer used as query (in this case the parent scaffold) and the top scoring database conformer (the top conformer of the aligned analog), the resulting .sdf file was split into two pieces using OpenEye's "chunker" application [S15,16] as follows:

```
chunker -in <analog_ID>.sdf -base <analog_ID__chunk_> -chunksize 1
```

The standard output from ROCS lists the query molecule first, and so only the second chunk file (containing the top conformer of the analog) was selected. To ensure the top conformer was properly protonated, the resulting .sdf file was run through the "fixpka" application in OpenEye's QUACPAC [S18] with the following command:

```
fixpka -in <analog_ID__chunk_0000002.sdf> -out <analog_ID__chunk_0000002_FIXPKA>.sdf
```

Finally, the analog's top scoring conformer following this step was parameterized for Rosetta compatibility and minimized as described above. The resulting minimized models were again ranked using vScreenML.

The top-scoring 18 compounds from vScreenML were visually inspected, then purchased directly from Enamine (**Table S1**).

*Identification of additional Enamine analogs for CP4.29*

Once our wetlab studies led us to prioritize CP4.29, we searched for additional analogs that could enrich our understanding of structure-activity relationships (SAR). By this time the REAL Space Navigator application had been discontinued, and so we used the newer infiniSee application [S21] to search the Enamine REAL Space database for additional analogs (version 2021-10, 21 billion compounds). Compounds that afforded the opportunity to evaluate the effect of individual substitutions (relative to CP4 or CP4.29) were selected for purchase.

*Plasmids, Cloning, and Site Directed Mutagenesis*

A polycistronic expression vector containing coding regions for an N-terminal 6xHis-Trx tagged pVHL19 protein construct, full length Elongin B, and Elongin C (residues 17-112) was generously provided by Dr. Song Tan [S22]. The plasmid was transformed into BL21(DE3) competent *E. Coli* according to manufacturer's protocols (New England Biolabs, catalog number C2527H), plated onto prewarmed 100 μg/mL ampicillin containing LBBA agar plates, and grown overnight at 37ºC. A single colony was selected and transferred to LB media with 100 μg/mL ampicillin sulfate. The starter culture was subsequently placed in a 37ºC shaker at 250 rpm overnight. The resulting plasmid DNA was extracted from the starter culture using a QIAprep Spin Miniprep Kit (Qiagen USA, catalog number 27140 or 27106) according to manufacturer's protocols. Final aliquots of the plasmid at a concentration of 50 ng/μL were frozen at -80ºC.

To incorporate the D197K mutation into VHL, two primers were purchased as HPLC purified oligomers with the following sequences: 5'-CAC CCA AAT GTG CAG AAA AAG CTG GAG CGG CTG ACA CAG-3' (forward) and 5'-CTG TGT CAG CCG CTC CAG CTT TTT CTG CAC ATT TGG GTG -3' (reverse). These were diluted to 100 μM in low EDTA, TE buffer. Using the QuikChange II Site-Directed Mutagenesis Kit (Agilent Technologies, catalog number 200523), the wild-type plasmid and primers were used to generate the $V_{D197K}$CB plasmid in accordance with manufacturer's protocols. The resulting $V_{D197K}$CB plasmid was transformed into Stellar chemically competent *E. Coli* (Takara Bio USA, catalog number 636763). Starter cultures were prepared, and plasmid DNA was isolated using the same method as described above. The sequence of the wild type and $V_{D197K}$CB constructs were validated using Sanger sequencing.

*Expression and Purification of the VCB and $V_{D197K}$CB Complexes*

To begin expression/purification of either WT VCB or mutant ($V_{D197K}$CB) complex, 2 μL of 50 ng/μL of the corresponding plasmid was transformed into BL21(DE3) *E. coli* according to manufacturer's protocols (New England Biolabs, Ipswich, MA; C2527H). After overnight growth of colonies at 37ºC on 100 μg/mL ampicillin containing LBBA agar plates, single colonies of each respective construct containing bacteria were transferred to one of two 300 mL Erlenmeyer flasks containing 100 mL of LB media with 100 μg/mL ampicillin sulfate and left to shake at 37ºC for 6 hours at 250 rpm. Starter cultures were subsequently diluted 1:100 into 2L of LB media containing the same concentration of ampicillin in appropriately labelled (i.e. "wild type" or "D197K") 4L Erlenmeyer flasks and left to shake at 190 rpm and 37ºC until the $OD_{600}$ = 0.6. Flasks were then placed at 4ºC for approximately one hour at which time IPTG was added to a final concentration of 200 µM to induce expression. After addition of IPTG, the flasks were placed back into a 16ºC shaker and left to shake overnight at 190 rpm. The following morning, bacteria expressing either construct were pelleted by centrifugation at 3,057xg at 4ºC for 20 minutes. Pellets from 4L worth of bacteria expressing wildtype VCB or pellets from 4L of bacteria expressing $V_{D197K}$CB were combined appropriately and frozen at -20ºC.

Both VCB and $V_{D197K}$CB were purified from their respective pellets using the same method described below. Pellets were first resuspended in 100-150 mL of 20 mM Tris, 500 mM NaCl, pH=7.5 buffer. After the addition of benzamidine HCl, lysozyme, and DNAse I from bovine pancreas, the suspension was left to stir vigorously at room temperature for at least 30 minutes. Bacteria were lysed by 3 rounds of French press (pressure between 500 – 1000 psi) at 4ºC. The lysates were cleared by centrifugation at 17,418xg for 20 min at 4ºC and supernatants were filtered through a syringe driven 0.2 µm PES filter. All remaining steps were conducted at 4ºC unless otherwise noted.

A HisTrap HP 5 mL affinity chromatography column (Cytiva Life Sciences; Marlborough, MA) was equilibrated with 10 CVs of 20 mM Tris, 500 mM NaCl, pH=7.5 buffer attached to an AKTA Pure protein purification system. Following equilibration, the column was detached and hooked up to a peristaltic pump (pre-equilibrated in the same buffer). The filtered supernatant was subsequently loaded onto the column using the peristaltic pump at a speed of approximately 3 mL/min. After loading, the column was re-attached to the AKTA system and washed at a flow rate of 3 mL/min for a total of 70 mL with 20 mM Tris, 500 mM NaCl, 20 mM imidazole, pH=7.5 buffer while collecting 2 mL fractions. Protein was eluted from the column over a 40 mL stretch using a linear gradient of the same buffer in which the imidazole concentration increased from 20 to 500 mM while taking 2 mL fractions. Fractions were analyzed for the target protein using SDS-PAGE and those confirmed to contain tagged VCB (or tagged $V_{D197K}$CB) were pooled.

The 6xHis-Trx tag was removed by placing the pooled fractions into a pre-soaked 15 mL dialysis cassette (10K MWCO) along with 3 mL of A280=4.0 TEV protease (stored at -20ºC in 20% glycerol containing buffer). The entire cassette was placed into 1 L of 20 mM Tris, 500 mM NaCl, pH=7.5 buffer overnight at 4$^{o}$C with slow stirring to ensure efficient dialysis of the imidazole. The following morning, the protein was removed from dialysis and manually loaded via syringe onto a HisTrap HP 5 mL column (pre-equilibrated with 10 CVs of 20 mM Tris, 500 mM NaCl, pH=7.5 buffer) at 4$^{o}$C and a rate of approximately 3 mL/min. The column was loaded back onto the AKTA and subjected to the same wash/elution steps already described. Consistent with efficient tag cleavage, the untagged VCB (or $V_{D197K}$CB) was identified in the wash fractions. The three most concentrated fractions were subsequently pooled and stored at 4$^{o}$C.

The pooled fractions were further purified using size exclusion chromatography with either a HiLoad 16/600 Superdex 200 pg or HiLoad 16/60 Superdex 75 pg column (Cytiva Life Sciences; Marlborough, MA) equilibrated in 20 mM Tris, 150 mM NaCl, 2 mM DTT, pH=7.0 buffer. Samples of 2 mL of either respective protein construct were injected onto the column at a rate of 1 mL/min. The protein was eluted using an isocratic elution in the same buffer at a rate of 1 mL/min and collected in 2 mL fractions. Fractions eluting at a rate consistent with the proper molecular weight were pooled and final protein purity was assessed by SDS-PAGE.

*Ligand Observed NMR Experiments*

Three separate ligand-observed NMR approaches were utilized to assess CP4 binding to the VCB complex: Saturation Transfer Difference (STD-NMR), WaterLOGSY, and Relaxation Filtered (or "CPMG") experiments. Each of these experiments is independently capable of detecting binding and can be considered orthogonal (with the caveat that STD and WaterLOGSY do leverage a similar underlying principle of magnetization transfer) [S23].

As a starting point, aliquots of 80 mM CP4, 49.6 mM sitagliptin phosphate, and 40 mM VH298 in DMSO-d6 were thawed from -80$^{o}$C to room temperature. CP4 was diluted to a working concentration of 40 mM with additional DMSO-d6. 1 µL of 40 mM CP4, 1 µL of 49.6 mM sitagliptin phosphate, 1 µL of 40 mM VH298, and 1 µL of DMSO-d6 were added to 3x1.5 mL microcentrifuge tubes and mixed gently by pipette. Tubes were capped and frozen at -80ºC until ready for use. Meanwhile, 2x15 mL Amicon 10K MWCO concentrators were used to concentrate the pooled fractions of VCB or $V_{D197K}$CB (following size exclusion chromatography) to a final volume of approximately 1 mL by centrifugation at 3000xg and 4ºC. Protein was subsequently placed on ice.

Meanwhile, two Nap-10 Sephadex G-25 DNA grade columns were drained of storage solution and washed with at least 15 mL of MilliQ water. Columns were equilibrated in 3x5 mL washes with ice cold 50 mM sodium phosphate buffer containing 10% $D_2O$/90% MilliQ water (v/v), pH=6.92 (here after referred to as "NMR buffer"). VCB or $V_{D197K}$CB samples (approximately 1 mL) were removed from ice and loaded onto Nap columns and allowed to drip through. Protein was eluted using 1.5 mL of NMR buffer, diluted to a final volume of 2 mL in 2 mL tubes, and placed on ice. Concentration of each protein was determined by absorbance at 280 nm ($\varepsilon$ = 25900 $M^{-1}$ $cm^{-1}$, MW = 42497 g/mole for VCB and MW=42510 g/mole for $V_{D197K}$CB) using a ThermoFisher NanoDrop 1000 spectrophotometer [S24]. Each solution of protein was measured five times to ensure robust determination of protein concentration. Since the $V_{D197K}$CB protein was the more dilute of the two (measured at 7.7 µM), an aliquot of the more concentrated wild type VCB was diluted in NMR buffer to the same concentration and a final volume of 2 mL. To ensure this dilution was done correctly, the concentration of this diluted wild type VCB was measured five times and found to also equal 7.7 µM. Final solutions of wild type VCB and $V_{D197K}$CB were immediately placed on ice.

Samples for NMR analysis were prepared by taking one of the tubes containing 4 µL of combined CP4/sitagliptin phosphate/VH298/DMSO-d6 and adding 396 µL of either 1) NMR buffer (kept on ice), 2) 7.7 µM VCB (in NMR buffer), or 3) 7.7 µM $V_{D197K}$CB (in NMR buffer) and mixing gently by pipette. The inclusion of a control sample containing no VCB (sample 1) for each experiment greatly diminishes the chances of a false positive result, while the matched concentrations of VCB and $V_{D197K}$CB rule out different protein concentrations as a potentially confounding variable.

For each sample, final compound concentrations were 100 µM for each of CP4, sitagliptin (after accounting for the phosphate), and VH298. DMSO-d6 was kept at 1%. Samples were transferred to NMR tubes and immediately placed into a 600 MHz Bruker NMR with TCI CryoProbe and Bruker Avance Neo console (running TopSpin 4.2.0) equilibrated at 278K. Four experiments consisting of a $^{1}H$ NMR with excitation sculpting water suppression, STD NMR, WaterLOGSY, and CPMG NMR were carried out in sequential order prior to preparing the next sample and repeating the process using protocols adapted from others' work [S25-27]. A detailed description of the protocol is provided below.

The sequence of experiments run on a given sample is as follows: 1) $^{1}H$ NMR with excitation sculpting water suppression (ZGESGP), 2) Saturation Transfer Difference NMR (SCREEN_STD), 3) WaterLOGSY (SCREEN_WLOGSY), and 4) CPMG (CPMGPR1D). To start, an existing experiment saved on the instrument was opened and the command "new" was typed into the command line. In the resulting dialogue box, a new folder name was entered and the EXPNO was changed to 1. Additionally, the "read parameterset" bubble was selected and the option for "ZGESGP" was selected. The checkbox for "Set solvent" was also checked and the "H2O+D2O" solvent option was chosen from the list. Under the "Additional action" section, the bubble for "Execute getprosol" was selected. In the title box, the experiment description was written and then the "OK" button was clicked. This resulted in a new experiment being opened in TopSpin. To lock onto the solvent, the command "lock" was typed into the command line and the option for "H2O+D2O" was selected. Once locked, the instrument was tuned by typing "atma" into the command line. Upon completion of tuning, the instrument was shimmed by first typing "topshim gui" into the command line. In the resulting dialog box, the "Dimension" option was set to "1D," the "Optimisation" option was set to "solvent suppression," the "Optimise for" option was set to "1H," and the "Use Z6" option was left unchecked. Under the "Tune" section of the dialog box, the "Before" and "After" were both set to "Z-X-Y-XZ-YZ-Z" and "Start" was clicked. Once shimming was complete, the sample icon was double clicked and the "Shim" tab was selected followed by the "Power" option. The power was adjusted until the lock signal was approximately 75% of the way to the top of the screen and "STDBY" was clicked. To determine the pulse length, the command "pulsecal -auto" was used and the number under "5.702W" was recorded. The pulse length was subsequently set for the experiment by typing "getprosol 1H #.## 5.7021W" where "#.##" was the pulse length recorded in the previous step. The receiver gain was set automatically using the command “rga” and the number of scans was set to 64 through the “ns” command. Finally, expected time for the experiment was determined via the “expt” command and the experiment was started via the “zg” command.

To begin the Saturation Transfer Difference (STD) experiment, the flask icon on the left side of the screen was clicked, followed by double clicking the “SCREEN_STD” option in the “Drug_discovery” library. In the resulting dialog box, the same options as described above for the “new” command were selected (with the EXPNO being updated to “2”). The newly created experiment (EXPNO 2) was double clicked in the “Data” list and the pulse length set with the following command “getprosol 1H #.## 5.7021W” as before using the same value for “#.##”. The number of scans was set to 128 using the “ns” command and the expected time of experiment was determined by the “expt” command. Finally, the experiment was started using the “zg” commad.

While the STD experiment was running, the WaterLOGSY experiment was queued up using the following protocol. To find the proper experiment, the flask icon was clicked and the “SCREEN_WLOGSY” option under the “Drug_discovery” library was selected. The same options as described above were selected in the dialog box (except for the appropriate change in the “EXPNO” value). The newly created experiment was selected in the “Data” list and the pulse length was set using same command (“getprosol 1H #.## 5.7021W”). The number of scans was set to 512 using the “ns” command. The number of dummy scans was adjusted to 4 by first clicking the “ACQUPARS” tab and changing the value in the “DS” box. The “SPECTRUM” button was clicked to close the “ACQUPARS” tab. Time of experiment was determined by the “expt” command. In order to queue up the next job in the scheduler, the Spooler icon was clicked and the “Job” button in the toolbar was clicked, followed by the “new” button. In the resulting box, the command “rga” was typed and OK was clicked. This same process was repeated with the “zg” command instead of the “rga” command to queue up the experiment.

Finally, a CPMG experiment was queued up on the sample using a similar protocol. First, the 1H excitation sculpting experiment (EXPNO 1) was opened and the “new” command was entered. The dialog box was filled in according to the same method described earlier for the first experiment with the EXPNO set to 4 (ZGESGP was still set as the "Read parameterset" option). After the new experiment was created, it was selected from the "Data" list. Under ACQUPARS, the pulse program was changed from "zgesgp" to "cpmgpr1d" by clicking the "…" icon across from the "PULPROG" option and selecting "cpmgpr1d" and clicking "Set PULPROG to Dataset." The pulse length was set to the same value as all other experiments for the sample using the same "getprosol 1H #.## 5.7021W" command. The loop counter was set to a value of 400 (corresponding to a KD value in the uM range)[S25] by typing the "L4" command and entering "400" into the box. The relaxation delay was set to 15 seconds using the "D1" command and entering "15" in the dialog box. The mixing time was set to 0.001 seconds by typing "D20" and setting the value to 0.001 seconds. The number of scans was set to 64 using the "ns" command. The length of the experiment was predicted using the "expt" command and the job was queued up using the same Spooler protocol described above for WaterLOGSY.

Upon completion of the experiments for a given sample, initial data processing was done for each experiment using TopSpin 4.2.0. For the 1H excitation sculpting water suppression experiment (zgesgp), the raw data was opened by double clicking the experiment from the "Data" list. The spectrum was viewed using the "efp" command and phased using the "apk" command. The WaterLOGSY experiments were processed simply by clicking the experiment and typing the "efp" command. The CPMG experiments were analyzed by opening the experiment, typing the "efp" command, and phasing the spectrum using the "apk" command.

Analysis of the STD experiment was carried out by adapting others' protocol [S26]. The initial 2D experiment was opened by double clicking the experiment from the "Data" list. The line broadening was set to a value of 3.0 Hz using the "lb 3" command. The "off resonance" spectrum was extracted using the "efp 1 2" command. The spectrum was phased using the "apk" command. To ensure the phasing was saved properly, the ".ph" command was entered followed by the ".s2d" and ".sret" commands. The original 2D experiment was reopened by typing the "rep 1" command. The "on" resonance spectrum was extracted using the "efp 2 3" command. The original 2D experiment was reopened using the same command as above and the "off" resonance spectrum was opened using the "rep 2" command. Multiple

Display mode was opened and the “on” resonance spectrum was pulled up by typing the “rep 2; .md” and “rep 3” commands, respectively. To calculate the difference between the “off” and “on” resonance spectra (i.e. the saturation transfer difference), the delta icon followed by the floppy disk icon was clicked. In the resulting dialog box, the number “5” was entered.

<u>*Cell Culture and Reagents*</u>

Human renal cell carcinoma (ccRCC) cell lines, including Caki-1 (CVCL_0234), 769-P (CVCL_1050), and RCC4 (CVCL_0498) were obtained from Fox Chase Cancer Center’s Cell Culture Facility (Philadelphia, PA). The RCC-MF (KTCTL-1M) cell line was generously provided by Dr. Philip Abbosh (Fox Chase Cancer Center, Philadelphia, PA).

Initial stocks were cryopreserved in FBS with 10% DMSO, and at every 6-month interval, a fresh aliquot of frozen cells was thawed and passaged for subsequent experiments. Cells were grown as monolayer in T75 flasks using RPMI- 1640 media supplemented with 10% fetal bovine serum, sodium pyruvate (1 mM), and non-essential amino acids (0.1 mM) and 1% penicillin/streptomycin at 37°C in 5% $CO_2$ atmosphere.

To compare the effect of CP4.29 under normoxic versus hypoxic conditions, a hypoxic culture glove box and incubator (Terra Universal) was used for continuous (uninterrupted) treatment and incubation of cells with 5µM CP4.29 at low oxygen conditions. Cells were first cultured on two 6-well plates at normoxic conditions (about 20% $O_2$) in the 5% $CO_2$ incubator until attachment. One plate was then transferred into hypoxic conditions (2% $O_2$ / 5% $CO_2$) in the glove box. Cells were left for 24 hours at each condition, then were treated with 5uM CP4.29 for 2 hours, then were lysed for Western blotting.

*VHL* knockout cells were generated by using Fugene lipofectamine from Promega (# E5911) to transfect cells with pLenti-Cas9-GFP (Addgene, # 83165) and sgRNA for *VHL* (Sg.VHL_1: 5`-CGCGGAGGGAATGCCCCGGA-3`, Sg.VHL_2: 5`-CCTCGGCGCCCAGTTCCTCC-3`, Sg.VHL_3: 5`-CCTCCCCGCCGTCTTCTTCA-3`). Knocked out clones were segregated into individual cells via

FACS cell sorting guided by the green florescence signal from the GFP-tag fused to Cas9. Effectiveness of the knockouts was verified in the expanded clones via Western blotting (**Figure S13**).

All reagents were of high purity grade and for use in cell culture experiments. MG132 and JNJ42041935 were obtained from EMD Millipore Corp. (# 474791 and # 400093, respectively). VH298 and MLN4924 were from Sigma Aldrich (# 1896, and # 951950, respectively). Belzutifan was from MedChemExpress (# HY-125840). Each was prepared in a stock of 40 mM and diluted to the desired culture treatment concentration.

*Western Blotting and Immunoprecipitation*

Adherent cells were rinsed once with ice-cold phosphate-buffered saline (PBS) (pH 7.4) and scraped in 100 μL of 2% SDS lysis buffer supplemented with complete protease inhibitors. Protein concentration of the resulting whole-cell lysates was determined by Pierce BCA Protein assay Kit (Thermo Scientific # 23228). For Western blotting, 20 μg of total protein for each sample was diluted in Laemmli sample buffer, heated at 95 °C for 5 min, loaded into 4- 20% Sure PAGE™, Bis-Tris gels (Gene Script # M00657), separated by sodium dodecyl sulfate polyacrylamide gel electrophoresis, and transferred to iBlot2 PVDF Regular Stacks System (Invitrogen # IB24001) according to the manufacturer's specifications. The membrane was then incubated for 40 min. at room temperature in 5% nonfat milk in TBS-T to block nonspecific antibody binding with constant agitation. Milk was washed off the membrane by two 5-min washes in TBS-T. The membrane was incubated overnight with constant agitation at 4 °C with primary antibodies (listed below) diluted in TBS-T supplemented with 1% protease-free bovine serum albumin and 0.05% sodium azide. The following day, the primary antibody solution was removed, and the membrane was washed three times with TBS-T for 5 min. Washed membranes were incubated with horseradish peroxidase (HRP)–conjugated secondary antibodies (goat anti-mouse immunoglobulin G [IgG] (Invitrogen # G21040) or goat anti-rabbit IgG (Invitrogen # G21234) diluted 1:5,000 in TBT-T containing 1% nonfat milk for 1 h at room temperature with constant agitation. The secondary-antibody solution was removed, and the membrane was washed three times with TBS-T for 5 min. Bound antibodies were detected with enhanced chemiluminescent HRP substrates SuperSignal West Pico PLUS Chemiluminescent Substrate (Thermo Scientific # 34578), or West Femto Maximum Sensitivity Substrate (Thermo Scientific # 34095), or West Atto Ultimate Sensitivity Substrate (Thermo Fisher # A38555), each used based on the intracellular protein level. Chemiluminescent signal was detected with (Fluorchem # E3031).

For immunoprecipitation (IP), cells were grown for treatment with either 0.1% DMSO or 5µM CP4.29 for 4 hours till harvesting with cold non-denaturing cell lysis buffer containing 1X complete protease inhibitor. The lysates were immunoprecipitated overnight using an anti-HIF2a or anti-AURKA or anti-ZHX2 primary antibodies. The immune complexes were incubated with pre-washed Protein A/G Sepharose Beads (Abcam, ab206996) for 4 hours, and washed three times with 1X cell lysis buffer before resuspending the pellet in loading dye and boiling for 5 min at 95 °C. Eluted proteins were immunoblotted with anti-ubiquitin.

Primary antibodies were used at the following dilutions:

- Rabbit anti-HIF2α, Cell Signaling Technologies (# 7096S), used at 1:1,000
- Mouse HIF1α, BD Biosciences (# 610958), used at 1:1,000
- Rabbit anti-VHL, Cell Signaling Technologies (# 68547S), used at 1:1,000
- Rabbit anti-AURKA, Cell Signaling Technologies (# 3092S), used at 1:1,000
- Rabbit anti-Fibronectin-1, Cell Signaling Technologies (# 26836S), used at 1:1,000
- Rabbit anti-ZHX2, Cell Signaling Technologies (# 20937S), used at 1:1,000
- Mouse anti-ubiquitin, Invitrogen (# 13-1600), used at 1:10,000

*Screening small molecules with Enzyme Linked Immunosorbent Assay (ELISA)*

Capture assay “sandwich” ELISAs were used to determine the levels of HIF2a (DuoSet IC ELISA, R&D Systems # DYC2997-2) after treatment with small molecules. RCC-MF cells were grown in 6-well plates overnight, treated with 20µM of small molecules for 2 hours, then lysed with a cocktail of 1mM EDTA, 0.5% TritonX-100, 5mM NaF, 6M urea, 1mM activated sodium orthovanadate, 2.5mM sodium pyrophosphate, 10 μg/ml leupeptin, 10 μg/ml pepstatin, 100 μM PMSF, 3 μg/ml aprotinin in PBS- pH 7.2-7.4. The protein concentration of each treatment was determined by BCA (ThermoFisher # 23228). Cell lysates were added to the pre-coated 96-well plates with HIF2a and the biotinylated detection antibody specific for HIF2a/ EPAS1 is used to detect the protein utilizing a standard Streptavidin-HRP format, with Spectromax Plate Reader.

*Reverse Transcription and qPCR*

769-P cells were treated with DMSO (vehicle control), 10 and 20 μM concentrations of CP4.29 along 20 μM belzutifan (positive control) for 4, 6, and 8 hours. RNA was extracted using phenol-chloroform based method with Trizol (Life Technologies # 15596018). RNA concentration was measured using NanoDrop Lite (ThermoFisher Scientific). First strand cDNA synthesis was performed with High-Capacity cDNA Reverse Transcription kit (Applied Biosystems # 4368814) according to manufacturer's instructions. The generated cDNA was diluted tenfold and used as a template for qPCR, which was performed with Applied Biosystems QuantStudio 5 system using PowerTrack™ SYBR Green Master Mix (Applied Biosystems # 4309155). Relative quantification of genes expression was performed using $2^{-\Delta\Delta Ct}$ method, and the primer sequences were:

CCND1 forward: CCGTCCATGCGGAAGATC

CCND1 reverse: ATGGCCAGCGGGAAGAC

VEGF forward: CGAAACCATGAACTTTCTGC

VEGF reverse: CATCCATGAACTTCACCACTTC

Actin forward: ACCAACTGGGACGACATGGAGAAA

Actin reverse: TAGCACAGCCTGGATAGCAACGTA.

*Cell proliferation assay*

Cells were plated in 8 replicates in 96-well plates (2000 cells/well) in 200 μL growth medium. Cells were treated with increasing concentrations of CP4.29 (0.05, 0.5, 5, and 10 μM) for 24, 48 and 72 hours. At the indicated time points, cells were changed with 190 µL fresh growth medium supplemented with 10 µL 12mM MTT reagent (Invitrogen # M6494) at 37 °C for 1-4 hours. The optical density value was detected at 540 nm using a 96-well plate reader.

*Colony formation assay*

A colony formation assay was used to analyze the effect of CP4.29 on cell growth. 769-P cells were used along their CRISPR knocked out generated clones of cells. Cells were seeded into six-well plates (20,000 cells/well). After four days, the colonies were stained with 0.5% crystal violet (Thermo Scientific # 40583-0250). The density of colonies formed in each well was measured using imageJ and then normalized to control wells (0.1% DMSO treated). Experiments were performed in triplicate, and pairwise test of comparison in the one-way ANOVA was used to analyze the difference between control and treatment groups.

*Wound healing assay*

Parental 769-P cells and *VHL* CRISPR-KO cells were seeded into six-well plates with a density of approximately 95% confluence after 24 h. The monolayer was lightly scratched with a 10-µl pipette tip across the well. After scratching, the well was washed with PBS and then replenished with serum-free medium supplemented with either 0.1% DMSO or 5 µM CP4.29 or 10 µM CP4.29 for 24 h. Wells were stained with 0.5% crystal violet (Thermo Scientific # 40583-0250) and the scratch area was measured with imageJ. Experiments were performed in triplicate, and pairwise test of comparison in the one-way ANOVA was used to analyze the difference between control and treatment groups.

*Statistical Analysis*

GraphPad Prism was used to analyze the statistical significances of data. All graphs depict mean ± SEM. *, ** and *** represent P value of < 0.05, 0.01, and 0.001, respectively. GraphPad Prism was used to generate all the graphs.

*Compound storage / preparation of purchased compounds*

Compounds purchased from Enamine arrived as dried powders/oils in amber vials. The initial 18 CP series compounds were dissolved in DMSO to a final concentration of 40 mM and several initial 40 μL aliquots were prepared in small amber vials. Aliquots (along with remaining volume in original vials) were frozen at -20°C. Prior to initial screening experiment using STD-NMR, aliquots of each compound were pooled into a pre-weighed vial and lyophilized overnight to dryness. Compounds were subsequently re-dissolved in DMSO-d6 to a final concentration of 160 mM (CP1, CP3, CP4, CP9, CP10, CP11, CP12, CP14, CP17, CP18), 80 mM (CP2, CP5), or 40 mM (CP6, CP7, CP8, CP13, CP15) depending on solubility and frozen at -20°C. The compound designated CP16 was not included in the initial screen.

Subsequent batches of CP4 were later delivered from Enamine and diluted to 80 mM in DMSO-d6. 10 μL aliquots were prepared in amber 1.5 mL microcentrifuge tubes and frozen at -80°C.

Compounds synthesized in-house (including CP4.29, CP4.35, CP4.36, CP4.40, CP4.41, CP4.42, CP4.43, CP4.44, CP4.45, and CP4.46) were dissolved into DMSO-d6 at a concentration of 160 mM and aliquoted. A single aliquot of each compound was further diluted to 40 mM, aliquoted, and frozen at -20°C in anticipation of cellular testing. Remaining aliquots of the 160 mM stocks were frozen at -80°C.

Additional analogs of CP4.29, termed CP4.1-CP4.34, were delivered from Enamine as either dry powders or oils in amber vials. The compounds designated CP4.33 and CP4.34 were not included in experimental characterization. Compounds were dissolved to a final concentration of either 80 mM or 160 mM in DMSO-d6 (depending on solubility) and frozen at -20°C.

Additional batches of CP4.29 used for experiments other than the initial screen were purchased directly from Enamine (catalog number Z5717230270), dissolved to 160 mM in DMSO-d6, and frozen at -80°C. The equivalence of our synthesized CP4.29 and the batches purchased from Enamine was confirmed by comparing each purchased batch to our reference $^{1}H$ and $^{13}C$ NMR spectra (**Figures S17-S18**).

Synthesized derivatives of CP4.29 were dissolved in DMSO-d6 to a final concentration of 40 mM and aliquoted into 1.5 mL microcentrifuge tubes. Two aliquots of each compound were frozen at -20°C for immediate use in cellular testing. Additional compound was frozen at -80°C for long term storage.

Sitagliptin phosphate was purchased as a powder and stored at -20°C. 8 mg of sitagliptin phosphate was dissolved in DMSO-d6 to a final concentration of 49.6 mM and aliquoted as 15 µL aliquots in 1.5 mL amber microcentrifuge tubes and frozen at -80°C. VH298 was purchased from Sigma Aldrich (catalog number SML1896-5MG) and dissolved to 40 mM in DMSO-d6. Aliquots of 15 µL were prepared in amber 1.5 mL microcentrifuge tubes and frozen at -80°C.

_Chemical Synthesis_

*Synthesis of (3,5-dichloro-1H-indol-2-yl)(4-((3-isopropyl-1,2,4-oxadiazol-5-yl)methyl)piperidin-1-yl)methanone (CP4.29)*

In a 20 mL vial, 53 mg (0.230 mmol) of 3,5-dichloro-1H-indole-2-carboxylic acid (Enamine Ltd., Kyiv, Ukraine; EN300-36960), 96.2 mg (0.218 mmol) BOP, 75.7 µL DIPEA (d=0.742 g/mL, 0.434 mmol) were combined and dissolved in 2 mL DMF. Solution was stirred at room temperature for 15 minutes at which time 53.4 mg (0.217 mmol) of 4-[[3-(propan-2-yl)-1,2,4-oxadiazol-5-yl]methyl]piperidine hydrochloride (Enamine Ltd; Kyiv, Ukraine; EN300-260977) were added. Reaction was allowed to run overnight at room temperature. The following morning, the DMF was removed using a rotary evaporator. Crude product was dissolved in ethyl acetate and filtered. Organic layer was washed with 1 N HCl followed by saturated sodium carbonate. Any precipitation was removed by filtration. The organic layer was dried with magnesium sulfate and filtered. Solvent was removed with a rotary evaporator before being dissolved back into DCM. Product was loaded onto a 40 g silica gel column equilibrated in DCM and eluted using a 0-10% MeOH gradient. Fractions containing product as determined by LC-MS were pooled and brought to dryness using a rotary evaporator. Product was brought up in ethyl acetate and washed with ammonium chloride followed by 1N HCl. The organic layer was then washed with saturated sodium carbonate followed by 1N NaOH. Product was assessed by LC-

MS and brought to dryness in a clean vial using a rotary evaporator. Notably, this entire procedure was repeated a second time and final product was pooled. Final compound weight was 14.3 mg corresponding to a 7.9% yield.

*Synthesis of (4-ethyl-1H-indol-2-yl)(4-((3-isopropyl-1,2,4-oxadiazol-5-yl)methyl)piperidin-1-yl)methanone (CP4.35)*

In a 20 mL vial, 52.0 mg (0.275 mmol) of 4-ethyl-1H-indole-2-carboxylic acid (Enamine Ltd; Kyiv, Ukraine; EN300-321400), 205 mg (0.539 mmol) HATU, and 133 µL of DIPEA (d=0.742 g/mL, 0.764 mmol) were combined and dissolved in 2 mL DMF. Solution was stirred at room temperature for 15 minutes at which time 82.9 mg (0.337 mmol) of 4-[[3-(propan-2-yl)-1,2,4-oxadiazol-5-yl]methyl]piperidine hydrochloride (Enamine Ltd; Kyiv, Ukraine; EN300-260977) were added. Reaction was left to run overnight at room temperature. The following morning, the DMF was removed using a rotary evaporator. Crude product was dissolved in ethyl acetate and filtered. The organic layer was washed three times with saturated ammonium chloride and brine three times. The organic layer was transferred to a clean vial and solvent was removed using a rotary evaporator. The remaining product was brought up in a 95% DCM/5% MeOH mixture and a precipitate was noted and filtered into a separate vial. The vial containing the product remaining in solution was dried down on a rotary evaporator, brought up in a minimal amount of DCM, and purified through a 40 g silica gel column using a DCM/MeOH gradient (0-10% MeOH). Fractions containing product (as assessed by LC-MS) were combined and brought to dryness on a rotary evaporator. NMR analysis of product showed trace amine impurities, so the product was re-dissolved in DCM, washed twice with 1 N HCl, and the organic layer was dried with $MgSO_4$. The dried organic layer was transferred to a clean pre-weighed vial and solvent removed with a rotary evaporator. Final product was stored a -20°C for 1 week before being dried in a desiccator. LC-MS analysis was done to confirm purity of final product. Final product weighed 14.4 mg, corresponding to a 13.8% yield.

*Synthesis of (4-ethyl-1H-indol-2-yl)(4-((4-methyl-4H-1,2,4-triazol-3-yl)methyl)piperidin-1-yl)methanone (CP4.36)*

In a 20 mL vial, 50 mg (0.264 mmol) of 4-ethyl-1H-indole-2-carboxylic acid (Enamine Ltd; Kyiv, Ukraine; EN300-321400), 1 eq BOP, and 2 eq DIPEA were combined and dissolved in 3 mL DHF. Solution was stirred at room temperature for 15 minutes at which point 67 mg (0.265 mmol) of 4-[(4-methyl-4H-1,2,4-triazol-3-yl)methyl]piperidine dihydrochloride (Enamine Ltd.; Kyiv, Ukraine; EN300-697451) were added. Reaction was left to run overnight at room temperature. The following morning, the DMF was removed using a rotary evaporator. Crude product was dissolved in ethyl acetate and filtered. The organic layer was washed three times with saturated ammonium chloride and brine. The organic layer was subsequently transferred to a clean vial and solvent was removed using a rotary evaporator. Product was brought up in a minimal amount of DCM, and purified through a 40 g silica gel column using a DCM/MeOH gradient (0-10% MeOH). Fractions containing product (as assessed by LC-MS) were combined and brought to dryness on a rotary evaporator. Product was dried on a vacuum desiccator overnight and confirmed by LC-MS. Final product weighed 14.7 mg corresponding to a 15.8% yield.

*Synthesis of (1-methyl-1H-indol-2-yl)(4-((4-methyl-4H-1,2,4-triazol-3-yl)methyl)piperidin-1-yl)methanone (CP4.40)*

In a 20 mL vial, 52.5 mg (0.300 mmol) of 1-methylindole-2-carboxylic acid, 141.7 mg (0.320 mmol) of BOP, and 142 µL of DIPEA (d=0.742 g/mL, 0.815 mmol) were combined and dissolved in 3 mL DHF. Solution was stirred at room temperature for 15 minutes at which point 87.5 mg (0.346 mmol) of 4-[(4-methyl-4H-1,2,4-triazol-3-yl)methyl]piperidine dihydrochloride (Enamine Ltd.; Kyiv, Ukraine; EN300-697451) were added. Reaction was left to run overnight at room temperature. The following morning, 5 drops of 1N HCl were added and the DMF was removed using a rotary evaporator. Attempted to dissolve crude product in EtOAc, but product remained an insoluble oil. EtOAc was decanted off and the crude product was dissolved in MeOH. LC-MS confirmed product was present in MeOH solution. MeOH was removed by rotary evaporator and crude product (an oil) was washed with ethyl acetate and DCM. Washes were discarded. Solvent was removed by rotary evaporator and product was placed in a vacuum desiccator overnight. The following morning, product was dissolved into 1.5 mL of MeOH and purified by reverse phase HPLC on a C18 column using a water/acetonitrile gradient (0-100% acetonitrile). Fractions confirmed to contain product (as determined by LC-MS) were pooled and solvent was removed by rotary evaporator. Final product was transferred to a clean vial and placed in a vacuum desiccator. Final product weighed 1.4 mg corresponding to a 1.38% yield.

*Synthesis of (4-((4-methyl-4H-1,2,4-triazol-3-yl)methyl)piperidin-1-yl)(6-phenylpyridin-2-yl)methanone (CP4.41)*

In a 20 mL vial, 53.3 mg of 6-phenyl-2-pyridinecarboxylic acid (0.268 mmol), 114.3 mg (0.258 mmol) BOP, 126.5 μL of DIPEA (d=0.742 g/mL, 0.726 mmol), and 3 mL of DMF were combined and allowed to stir for 15 minutes at room temperature. To the mixture, 85.4 mg (0.337 mmol) of 4-[(4-methyl-4H-1,2,4-triazol-3-yl)methyl]piperidine dihydrochloride (Enamine Ltd.; Kyiv, Ukraine; EN300-697451) were added. The reaction was allowed to run overnight at room temperature. The following morning, 5 drops of 1N HCl were added and solvent was removed using a rotary evaporator. Attempted to dissolve crude product in EtOAc, but product remained an insoluble oil. EtOAc was decanted off and the crude product was dissolved in MeOH. LC-MS confirmed product was present in MeOH solution. MeOH was removed by rotary evaporator and crude product (an oil) was washed with ethyl acetate and DCM. Washes were discarded. Solvent was removed by rotary evaporator and product was placed in a vacuum desiccator overnight. The following day, the product was dissolved in EtOH/EtOAc (1:1) mixture and heated. Product was found to be soluble in the EtOH/EtOAc solution by LC-MS. Transferred product to clean vial and removed solvent with rotary evaporator. The product was subsequently dissolved in MeOH and purified using reverse phase HPLC on a C18 column using a water/acetonitrile gradient (0-100% acetonitrile). Fractions confirmed to contain product (as determined by LC-MS) were pooled and solvent was removed by rotary evaporator. Final product was transferred to a clean vial and placed in a vacuum desiccator. Final product weighed 3.7 mg corresponding to a 4.0% yield.

*Synthesis of (S)-2-(6-methoxynaphthalen-2-yl)-1-(4-((4-methyl-4H-1,2,4-triazol-3-yl)methyl)piperidin-1-yl)propan-1-one (CP4.42)*

In a 20 mL vial, 47.6 mg (0.207 mmol) of (S)-2-(6-methoxynaphthalen-2-yl)propanoic acid, 94.0 mg (0.213 mmol) BOP, 104.8 µL of DIPEA (d=0.742 g/mL, 0.602 mmol), and 3 mL of DMF were combined and allowed to stir for 15 minutes at room temperature. To the mixture, 62.7 mg (0.248 mmol) of 4-[(4-methyl-4H-1,2,4-triazol-3-yl)methyl]piperidine dihydrochloride (Enamine Ltd.; Kyiv, Ukraine; EN300-697451) were added. The reaction was allowed to run overnight at room temperature. The following morning, 5 drops of 1N HCl were added and solvent was removed using a rotary evaporator. Attempted to dissolve crude product in EtOAc, but product remained an insoluble oil. Washed oil a second time with EtOAc and collected washings. Residual solvent was removed from the oil with a rotary evaporator. The oil was dissolved in MeOH and a sample was used to run LC-MS. Solvent was removed from remaining product using a rotary evaporator and the vial was placed in a desiccator. Final product weighed 94.1 mg corresponding to a 116% yield.

*Synthesis of benzofuran-2-yl(4-((3-isopropyl-1,2,4-oxadiazol-5-yl)methyl)piperidin-1-yl)methanone (CP4.43)*

In a 20 mL vial, 50 mg (0.277 mmol) of benzofuran-2-carbonyl chloride, 136 mg (0.554 mmol) of 4-[[3-(propan-2-yl)-1,2,4-oxadiazol-5-yl]methyl]piperidine hydrochloride (Enamine Ltd; Kyiv, Ukraine; EN300-260977), and 3 mL pyridine were combined. Reaction was allowed to run overnight at room temperature. The following morning, the crude product was filtered and solvent was removed by a rotary evaporator. The crude product was dissolved in MeOH and a sample was analyzed by LC-MS. Product was purified using reverse phase HPLC on a C18 column using a water/acetonitrile gradient (0-100% acetonitrile). Fractions confirmed to contain product (as determined by LC-MS) were pooled and solvent was removed by rotary evaporator. Final product was transferred to a clean vial and placed in a vacuum desiccator. Final product weighed 31.8 mg corresponding to a 32.5% yield.

*Synthesis of (5-bromofuran-2-yl)(4-((3-isopropyl-1,2,4-oxadiazol-5-yl)methyl)piperidin-1-yl)methanone (CP4.44)*

In a 20 mL vial, 50 mg (0.262 mmol) of 5-bromofuran-2-carboxylic acid, 99.5 mg (0.262 mmol) HATU, 137 µL of DIPEA (d=0.742 g/mL, 0.787 mmol), and 3 mL of DMF were combined and allowed to stir for 15 minutes at room temperature. To the mixture, 64.3 mg (0.262 mmol) of 4-[[3-(propan-2-yl)-1,2,4-oxadiazol-5-yl]methyl]piperidine hydrochloride (Enamine Ltd; Kyiv, Ukraine; EN300-260977) were added. The reaction was allowed to run overnight at room temperature. The following morning, the crude product was filtered, and solvent was removed by a rotary evaporator. The crude product was dissolved in MeOH, a sample of which was analyzed by LC-MS. Product was purified using reverse phase HPLC on a C18 column using a water/acetonitrile gradient (0-100% acetonitrile). Fractions confirmed to contain product (as determined by LC-MS) were pooled and solvent was removed by rotary evaporator. Final product was transferred to a clean vial and placed in a vacuum desiccator. Final product weighed 21.3 mg corresponding to a 21.3% yield.

*Synthesis of (4-ethyl-1H-indol-2-yl)(piperidin-1-yl)methanone (CP4.45)*

In a 20 mL vial, 50 mg (0.264 mmol) of 4-ethyl-1H-indole-2-carboxylic acid (Enamine Ltd; Kyiv, Ukraine; EN300-321400), 116.8 mg (0.264 mmol) BOP, 92 µL DIPEA (d=0.742 g/mL, 0.528 mmol), and 3 mL of DMF were combined and allowed to stir for 15 minutes at room temperature. To the mixture, 26.7 µL of piperidine (d=0.862 g/mL, 23 mg, 0.270 mmol) were added. The reaction was allowed to run overnight at room temperature. The following morning, the DMF was removed using a rotary evaporator. Crude product was dissolved in ethyl acetate and filtered. Organic layer was washed with 1 N HCl followed by saturated sodium carbonate. Any precipitation was removed by filtration. The organic layer was dried with sodium sulfate and filtered. Solvent was removed with a rotary evaporator before being dissolved back into DCM. Product was loaded onto a 40 g silica gel column equilibrated in DCM and eluted using a 0-10% MeOH gradient. Fractions containing product as determined by LC-MS were pooled and brought to dryness using a rotary evaporator. Product was assessed by LC-MS and brought to dryness in a clean vial using a rotary evaporator. Final product weighed 22.0 mg corresponding to a 32.5% yield.

*Synthesis of (4-ethyl-1H-indol-2-yl)(morpholino)methanone (CP4.46)*

In a 20 mL vial, 50 mg (0.264 mmol) of 4-ethyl-1H-indole-2-carboxylic acid (Enamine Ltd; Kyiv, Ukraine; EN300-321400), 116.8 mg (0.264 mmol) BOP, 92 µL DIPEA (d=0.742 g/mL, 0.528 mmol), and 3 mL of DMF were combined and allowed to stir for 15 minutes at room temperature. To the mixture, 22.8 µL of morpholine (d=1.007 g/mL, 0.264 mmol) were added. The reaction was allowed to run overnight at room temperature. The following morning, the reaction was filtered and allowed to continue for 1 additional day. Following the second day of the reaction, DMF was removed using a rotary evaporator. Crude product was dissolved in ethyl acetate and filtered. Organic layer was washed with saturated ammonium chloride. The organic layer was dried with sodium sulfate and filtered. Solvent was removed with a rotary evaporator before being dissolved back into DCM. Product was loaded onto a 40 g silica gel column equilibrated in DCM and eluted using a 0-10% MeOH gradient. Fractions containing product as determined by LC-MS were pooled and brought to dryness using a rotary evaporator. Product was assessed by LC-MS and brought to dryness in a clean vial using a rotary evaporator. Final product weighed 22.2 mg corresponding to a 32.5% yield.

*Synthesis of (3,5-dichloro-1H-indol-2-yl)(4-((3-ethyl-1,2,4-oxadiazol-5-yl)methyl)piperidin-1-yl)methanone (CP4.29.1)*

In a 20 mL vial, 52.7 mg (0.229 mmol) of 3,5-dichloro-1H-indole-2-carboxylic acid (Enamine Ltd., Kyiv, Ukraine; EN300-36960), 104.7 mg (0.275 mmol) HATU, 120 µL DIPEA (d=0.742 g/mL, 0.687 mmol) and 3 mL of dry dimethylformamide were added and allowed to mix with rapid stirring for 15 minutes. To the mixture, 61.6 mg of 4-[(3-ethyl-1,2,4-oxadiazol-5-yl)methyl]piperidine (0.315 mmol; Enamine Ltd., Kyiv, Ukraine; EN300-304609) were added and reaction was allowed to stir overnight at room temperature. Crude product was filtered, transferred to a 40 mL vial, and DMF removed using a rotary evaporator. Product was partitioned between water and dichloromethane (DMC). Organic layer was washed twice with water, dried with sodium sulfate, and transferred to a new vial. DCM was removed with a rotary evaporator and product was brought up in methanol (MeOH) for LC-MS analysis. MeOH was removed in similar fashion and product left dry overnight. Product was brought up in 1-3 mL of DCM and purified using normal phase chromatography on a 4 g silica column and a DCM/MeOH (0-10%) gradient. Fractions were allowed to dry via evaporation in the hood and pulled up in MeOH for LC-MS analysis. Product was dried using a rotary evaporator, dissolved in DMSO, and purified using reverse phase HPLC on a C18 column using a linear water/acetonitrile gradient (with 0.1% formic acid). Fractions containing product as assessed by LC-MS were pooled into a weighed vial and dried using a Genevac with maximum temperature set to 60°C. Product was dissolved in $CDCl_3$ and analyzed by 1H NMR on a Bruker 400 MHz NMR spectrometer. Sample was returned to vial and $CDCl_3$ was allowed to evaporate in the hood. Product was re-dissolved in 85% acetonitrile/15% water mixture and dried on Genevac overnight on the same settings. Product was dissolved in DCM and transferred to a new pre-weighed 4 mL vial and solvent allowed to evaporate overnight in the hood. Vial was covered with a Kim wipe and placed in a vacuum desiccator overnight. Final compound dry weight was 7.3 mg which corresponded to a 7.8% yield. 1H NMR ($CDCl_3$, 400 MHz): δ 9.27 (s, 1H), 7.38 (d, 1H, J=1.92Hz), 7.08 (dd, 1H, J=8.76 Hz, 0.32 Hz), 7.04 (residual $CHCl_3$), 7.03 (dd, 1H, J=8.72 Hz, 1.92 Hz), 4.20 (br, 2H), 2.80 (br, 2H), 2.64 (d, 2H, J=7.08 Hz), 2.54 (q, 2H, J=7.6 Hz), 2.00 (m, 1H), 1.66 (br, 2H), 1.22 (qd, 2H, J=12.52 Hz, 3.84 Hz), 1.11 (t, 3H, J= 7.56 Hz). Found m/z = 407.1 $(M+H)^+$.

*Synthesis of (4-((3-cyclobutyl-1,2,4-oxadiazol-5-yl)methyl)piperidin-1-yl)(3,5-dichloro-1H-indol-2-yl)methanone (CP4.29.3)*

In a 20 mL vial, 53.2 mg (0.231 mmol) of 3,5-dichloro-1H-indole-2-carboxylic acid (Enamine Ltd., Kyiv, Ukraine; EN300-36960), 107.4 mg (0.283 mmol) HATU, 121 µL DIPEA (d=0.742 g/mL, 0.694 mmol) and 3 mL of dry dimethylformamide were added and allowed to mix with rapid stirring for 15 minutes. To the resulting mixture, 80.6 mg of 4-[(3-cyclobutyl-1,2,4-oxadiazol-5-yl)methyl]piperidine hydrochloride (0.313 mmol; Enamine Ltd., Kyiv, Ukraine; EN300-262395) were added and the reaction was left to stir overnight at room temperature. Product was purified as described above for CP4.29.1. Final dry compound weight was 7.8 mg which corresponded to an 7.8% yield as well. 1H NMR ($CDCl_3$, 400 MHz): δ 9.75 (s, 1H), 7.50 (d, 1H, J=1.92 Hz), 7.20 (d, 1H, J= 8.72 Hz), 7.19 (residual $CHCl_3$), 7.14 (dd, 1H, J= 8.76 Hz, 1.96 Hz), 4.30 (br, 2H), 3.58 (qd, 1H, J= 17.04 Hz, 0.92 Hz), 3.00 (br, 2H), 2.79 (d, 2H), 2.30 (m, 4H), 2.14 (m, 1H), 2.03 (m, 1H), 1.90 (m, 1H), 1.80 (br, 2H), 1.37 (m, 2H). Found m/z = 433.1 $(M+H)^+$.

*Synthesis of (4-((3-cyclopentyl-1,2,4-oxadiazol-5-yl)methyl)piperidin-1-yl)(3,5-dichloro-1H-indol-2-yl)methanone (CP4.29.4)*

In a 20 mL vial, 54.1 mg (0.235 mmol) of 3,5-dichloro-1H-indole-2-carboxylic acid (Enamine Ltd., Kyiv, Ukraine; EN300-36960), 108.9 mg (0.286 mmol) HATU, 123 µL DIPEA (d=0.742 g/mL, 0.705 mmol) and 3 mL of dry dimethylformamide were added and allowed to mix with rapid stirring for 15 minutes. To the resulting mixture, 77.8 mg of 4-[(3-cyclopentyl-1,2,4-oxadiazol-5-yl)methyl]piperidine hydrochloride (0.286 mmol; Enamine Ltd., Kyiv, Ukraine; EN300-313578) were added and the reaction was left to stir overnight at room temperature. Product was purified as described above for CP4.29.1. Final dry compound weight was 4.6 mg which corresponded to a 4.38% yield. 1H NMR ($CDCl_3$, 400 MHz): δ 9.20 (s, 1H), 7.53 (d, 1H, J=1.88 Hz), 7.22 (d, 1H, J= 8.48 Hz), 7.19 (residual $CHCl_3$), 7.17 (dd, 1H, J = 8.72 Hz, 1.92 Hz), 4.25 (br, 2H), 2.90 (br, 2H), 3.12 (p, 2H, J=7.8 Hz), 2.78 (d, 2H, J=7.04 Hz), 2.14 (m, 1H), 1.98 (m, 2H), 1.75 (m, 7H), 1.62 (m, 5H), 1.35 (qd, 2H, J=12.6 Hz, 3.96 Hz). Found m/z = 447.1 $(M+H)^+$.

*Synthesis of (4-((3-cyclohexyl-1,2,4-oxadiazol-5-yl)methyl)piperidin-1-yl)(3,5-dichloro-1H-indol-2-yl)methanone (CP4.29.5)*

In a 20 mL vial, 53.8 mg (0.234 mmol) of 3,5-dichloro-1H-indole-2-carboxylic acid (Enamine Ltd., Kyiv, Ukraine; EN300-36960), 108.2 mg (0.285 mmol) HATU, 122 µL DIPEA (d=0.742 g/mL, 0.702 mmol) and 3 mL of dry dimethylformamide were added and allowed to mix with rapid stirring for 15 minutes. To the resulting mixture, 85.3 mg of 4-[(3-cyclohexyl-1,2,4-oxadiazol-5-yl)methyl]piperidine hydrochloride (0.298 mmol; Enamine Ltd., Kyiv, Ukraine; EN300-312344) were added and the reaction was left to stir overnight at room temperature. Product was purified as described above for CP4.29.1. Final dry weight of the product was 3.4 mg which corresponded to a 3.1% yield. 1H NMR ($CDCl_3$, 400 MHz): δ 9.28 (s, 1H), 7.52 (d, 1H, J=1.88 Hz), 7.21 (d, 1H, J=8.72 Hz), 7.19 (residual $CHCl_3$), 7.17 (dd, 1H, J=8.72 Hz, 1.92 Hz), 4.30 (br, 2H), 3.00 (br, 2H), 2.78 (d, 2H, J=7.08 Hz), 2.71 (tt, 1H, J=11.48 Hz, 3.56 Hz), 2.14 (m, 1H), 1.92 (m, 2H), 1.77 (m, 5H), 1.64 (m, 4H), 1.50 (qd, 3H, J=11.92 Hz), 1.40 (m, 7H). Found m/z = 461.1 $(M+H)^+$.

*Synthesis of (3,5-dichloro-1H-indol-2-yl)(4-((3-(tetrahydro-2H-pyran-4-yl)-1,2,4-oxadiazol-5-yl)methyl)piperidin-1-yl)methanone (CP4.29.6)*

In a 20 mL vial, 53.3 mg (0.232 mmol) of 3,5-dichloro-1H-indole-2-carboxylic acid (Enamine Ltd., Kyiv, Ukraine; EN300-36960), 113.7 mg (0.299 mmol) HATU, 121 µL DIPEA (d=0.742 g/mL, 0.696 mmol) and 3 mL of dry dimethylformamide were added and allowed to mix with rapid stirring for 15 minutes. To the resulting mixture, 82.7 mg of 4-[[3-(oxan-4-yl)-1,2,4-oxadiazol-5-yl]methyl]piperidine hydrochloride (0.287 mmol; Enamine Ltd., Kyiv, Ukraine; EN300-318107) were added and the reaction was left to stir overnight at room temperature. Product was purified as described above for CP4.29.1. Final compound weight was 6.7 mg which corresponded to a 6.2% yield. 1H NMR ($CDCl_3$, 400 MHz): δ 9.39 (s, 1H), 7.62 (d, 1H), 7.31 (d, 1H, J= 8.72 Hz), 7.28 (residual $CHCl_3$), 7.26 (dd, 1H, J = 8.72 Hz, 1.92 Hz), 4.40 (br, 2H), 4.05 (dt, 2H, J = Hz), 3.55 (td, 2H, J = Hz), 3.1 (br, 2H), 3.05 (septet, 1H, J = Hz), 2.88 (d, 2H, J = Hz), 2.23 (m, 1H), 1.94 (m, 7H), 1.68 (s, 2H), 1.46 (qd, 2H, J = Hz). Found m/z = 463.1 $(M+H)^+$.

*Synthesis of (3,5-dichloro-1H-indol-2-yl)(4-(piperidin-1-ylmethyl)piperidin-1-yl)methanone (CP4.29.9)*

In a 20 mL vial, 47.7 mg (0.207 mmol) of 3,5-dichloro-1H-indole-2-carboxylic acid (Enamine Ltd., Kyiv, Ukraine; EN300-36960), 102 mg (0.268 mmol) HATU, 108 µL DIPEA (d=0.742 g/mL, 0.621 mmol) and 3 mL of dry dimethylformamide were added and allowed to mix with rapid stirring for 15 minutes. To the resulting mixture, 65.2 mg of 4-[(piperidin-1-yl)methyl]piperidine dihydrochloride (0.255

mmol; Enamine Ltd., Kyiv, Ukraine; EN300-54899) were added and the reaction was left to stir overnight at room temperature. Product was purified as described above for CP4.29.1. Final compound dry weight was 2.7 mg corresponding to a yield of 3.3%. 1H NMR ($CDCl_3$, 400 MHz): δ 10.27 (s, 1H), 8.48 (s, 1H), 8.04 (s, 0.2 H), 7.60 (d, 1H, J = 1.84 Hz), 7.35 (d, 1H, J = 8.76 Hz), 7.28 (residual $CHCl_3$), 7.23 (dd, 1H, J = 8.72 Hz, 2.0 Hz), 4.50 (br, 6H), 2.98 (m, 5H), 2.91 (s, 1H), 2.79 (br, 2H), 2.64 (s, 0.3H), 2.18 (m, 1H), 2.11 (s, 0.17H), 2.02 (m, 2H), 1.89 (m, 5H), 1.61 (br, 2H), 1.41 (qd, 2H, J = 12.56 Hz, 3.52 Hz). Found m/z = 394 $(M+H)^+$.

*Synthesis of (3,5-dichloro-1H-indol-2-yl)(4-(morpholinomethyl)piperidin-1-yl)methanone (CP4.29.10)*

In a 20 mL vial, 50.2 mg (0.218 mmol) of 3,5-dichloro-1H-indole-2-carboxylic acid (Enamine Ltd., Kyiv, Ukraine; EN300-36960), 105.3 mg (0.277 mmol) HATU, 114 µL DIPEA (d=0.742 g/mL, 0.654 mmol) and 3 mL of dry dimethylformamide were added and allowed to mix with rapid stirring for 15 minutes. To the resulting mixture, 68.3 mg of 4-[(piperidin-4-yl)methyl]morpholine (0.371 mmol; Enamine Ltd., Kyiv, Ukraine; EN300-42119) were added and the reaction was left to stir overnight at room temperature. Product was purified as described above for CP4.29.1 with several slight modifications. During aqueous workup, the organic layer was washed once with water and once with saturated brine. Additionally, the aqueous layer was reverse extracted with DCM and the resulting organic layer pooled with the original organic layer. Final compound dry weight was 18.2 mg corresponding to a yield of 21.1%. 1H NMR ($CDCl_3$, 400 MHz): δ 10.25 (s, 1H), 8.27 (s, 3H), 7.60 (d, 1H, J =1.6 Hz), 7.32 (dd, 1H, J = 8.76 Hz, 0.36 Hz), 7.28 (residual $CHCl_3$), 7.24 (dd, 1H, J = 8.76 Hz, 2.0 Hz), 4.40 (br, 2H), 3.85 (t, 4H, J = Hz), 3.00 (br, 2H), 2.99 (s, 0.4H), 2.91 (s, 0.4H), 2.56 (d, 2H, J=6.72 Hz), 2.00 (m, 3H), 1.35 (qd, 2H, J=12.12 Hz, 2.96 Hz). Found m/z = 396.1 $(M+H)^+$.

*Synthesis of (3,5-dichloro-1H-indol-2-yl)(4-(pyridin-3-ylmethyl)piperidin-1-yl)methanone (CP4.29.11)*

In a 20 mL vial, 49.9 mg (0.217 mmol) of 3,5-dichloro-1H-indole-2-carboxylic acid (Enamine Ltd., Kyiv, Ukraine; EN300-36960), 107.8 mg (0.284 mmol) HATU, 113 µL DIPEA (d=0.742 g/mL,

0.651 mmol) and 3 mL of dry dimethylformamide were added and allowed to mix with rapid stirring for 15 minutes. To the resulting mixture, 61.1 mg of 3-[(piperidin-4-yl)methyl]pyridine (0.347 mmol; Enamine Ltd., Kyiv, Ukraine; EN300-122729) were added and the reaction was left to stir overnight at room temperature. Product was purified as described above for CP4.29.10. Final compound dry weight was 5.5 mg corresponding to a yield of 6.5%. 1H NMR ($CDCl_3$, 400 MHz): δ 9.49 (s, 1H), 8.53 (s, 2H), 7.60 (m, 2H), 7.36 (br, 1H), 7.32 (d, 1H, J=8.72 Hz), 7.28 (residual $CHCl_3$), 7.26 (dd, 1H, J = 8.72 Hz, 1.96 Hz), 4.40 (br, 2H), 3.00 (br, 2H), 2.65 (d, 2H, J = Hz), 2.50 (br, 2H), 1.90 (m, 2H), 1.38 (qd, 2H, J = 12.32 Hz, 3.64 Hz). Found m/z = 388 $(M+H)^+$.

*Synthesis of (4-(cyclopentylmethyl)piperidin-1-yl)(3,5-dichloro-1H-indol-2-yl)methanone (CP4.29.12)*

In a 20 mL vial, 49.5 mg (0.215 mmol) of 3,5-dichloro-1H-indole-2-carboxylic acid (Enamine Ltd., Kyiv, Ukraine; EN300-36960), 103.9 mg (0.273 mmol) HATU, 112 µL DIPEA (d=0.742 g/mL, 0.645 mmol) and 3 mL of dry dimethylformamide were added and allowed to mix with rapid stirring for 15 minutes. To the resulting mixture, 52.0 mg of 4-(cyclopentylmethyl)piperidine hydrochloride (0.255 mmol; Enamine Ltd., Kyiv, Ukraine; EN300-240902) were added and the reaction was left to stir overnight at room temperature. Product was purified as described above for CP4.29.10. Final compound dry weight was 3.5 mg corresponding to a yield of 4.3%. 1H NMR ($CDCl_3$, 400 MHz): δ 9.62 (s, 1H), 7.61 (d, 1H, J = 1.96 Hz), 7.31 (d, 1H, J = 8.76 Hz), 7.28 (residual $CHCl_3$), 7.24 (dd, 1H, J= 8.76 Hz, 2.0 Hz), 4.30 (br, 2H), 3.10 (br, 2H), 1.80 (m, 6H), 1.6 (m, 7H), 1.30 (m, 5H), 1.10 (m, 2H). Found m/z = 379.1 $(M+H)^+$.

*Synthesis of (4-((1H-pyrrol-1-yl)methyl)piperidin-1-yl)(3,5-dichloro-1H-indol-2-yl)methanone (CP4.29.13)*

In a 20 mL vial, 49.2 mg (0.214 mmol) of 3,5-dichloro-1H-indole-2-carboxylic acid (Enamine Ltd., Kyiv, Ukraine; EN300-36960), 101.7 mg (0.267 mmol) HATU, 112 µL DIPEA (d=0.742 g/mL, 0.642 mmol) and 3 mL of dry dimethylformamide were added and allowed to mix with rapid stirring for 15 minutes. To the resulting mixture, 114 mg of 4-[(1H-pyrrol-1-yl)methyl]piperidine (0.694 mmol;

Enamine Ltd., Kyiv, Ukraine; EN300-262763) were added and the reaction was left to stir overnight at room temperature. Product was purified as described above for CP4.29.10. Final compound dry weight was 3.3 mg corresponding to a yield of 4.1%. 1H NMR ($CDCl_3$, 400 MHz): δ 9.58 (s, 1H), 7.61 (d, 1H, J = 1.92 Hz), 7.29 (dd, 1H, J = 8.76 Hz, 0.4 Hz), 7.28 (residual $CHCl_3$), 7.25 (dd, 1H, J = 8.72 Hz, 1.96 Hz), 6.64 (t, 2H, J = 2.04 Hz), 6.18 (t, 2H, J = 2.12 Hz), 4.40 (br, 2H), 3.81 (d, 2H, J = 7.2 Hz), 3.10 (br, 2H), 2.04 (m, 1H), 1.73 (d, 4H, J = 10.44 Hz), 1.34 (qd, 2H, J = 12.52 Hz, 3.8 Hz). Found m/z = 376.1 $(M+H)^+$.

*Synthesis of (3,5-dimethyl-1H-indol-2-yl)(4-((3-isopropyl-1,2,4-oxadiazol-5-yl)methyl)piperidin-1-yl)methanone (CP4.29.14)*

In a 20 mL vial, 99.5 mg (0.526 mmol) of 3,5-dimethyl-1H-indole-2-carboxylic acid (Enamine Ltd., Kyiv, Ukraine; EN300-103388), 246.5 mg (0.648 mmol) HATU, 275 µL DIPEA (d=0.742 g/mL, 1.578 mmol) and 3 mL of dry dimethylformamide were added and allowed to mix with rapid stirring for 10 minutes. To the resulting mixture, 155.3 mg of 4-[[3-(propan-2-yl)-1,2,4-oxadiazol-5-yl]methyl]piperidine hydrochloride (0.632 mmol; Enamine Ltd., Kyiv, Ukraine; EN300-260977) were added and the reaction was left to stir overnight at room temperature. The following day, the DMF was removed using a rotary evaporator and the crude product was washed with 10 mL of water twice (washings were saved). 5-10 mL of ethyl acetate were added to the crude product but the product remained insoluble. Ethyl acetate was removed using a rotary evaporator and the product was dissolved in 10 mL of dichloromethane. The resulting solution was added to the saved initial aqueous washes and product was partitioned between the DCM and water layers.

After removal of the aqueous layer, the organic layer was washed once with 10 mL of 2N HCl, once with 10 mL of saturated sodium bicarbonate, and once with 10 mL of brine. Remaining product in the combined aqueous waste was reverse extracted using 10 mL of DCM and the washes repeated before combining the organic layers together. The organic layer was dried with sodium sulfate and transferred to new vials at which point the DCM was removed with a rotary evaporator. 5 mL of methanol (MeOH) was added to the crude product at which point a white solid crashed out of solution. Methanol solution was removed and placed into a second vial leaving behind the white solid. The remaining white solid was washed twice more with methanol and the methanol washes were pooled. A sample of the white solid was dissolved in 1 mL of DMSO and analyzed by LC-MS which confirmed the compound identity as product.

The remaining product had solubility tested in 10 mL of hot methanol after which it was dried on a rotary evaporator and dissolved in 3 mL of DCM. A small sample of the resulting solution was transferred to a new vial and the DCM was removed by evaporation. The sample was dissolved in 1 mL of DMSO and analyzed using LC-MS. Remaining product was dissolved in $CDCl_3$ and analyzed by 1H NMR spectroscopy on a Bruker 400 MHz NMR spectrometer. The product was recovered and the $CDCl_3$ was removed via evaporation in the hood. Product was dissolved in DMSO and purified further using reverse phase HPLC on a C18 column with a water/acetonitrile gradient (with 0.1% formic acid). Fractions containing product were pooled and dried down on the Genevac with a maximum temperature of 60°C. The resulting product was once again dissolved in $CDCl_3$ and analyzed by 1H NMR. After evaporation of $CDCl_3$, the product was dissolved in DCM and transferred to a clean 4 mL vial, covered with a Kim wipe, and left in a vacuum desiccator overnight. Final compound weight was 11.8 mg which corresponded to a 5.9% yield. 1H NMR ($CDCl_3$, 400 MHz): δ 8.55 (s, 1H), 7.38 (s, 1H), 7.27 (d, 1H, J = 9.48 Hz), 7.10 (dd, 1H, J = 8.4 Hz, 1.32 Hz), 4.38 (s, 2H), 3.10 (m, 3H), 2.85 (d, 2H, J = Hz), 2.48 (s, 3H), 2.35 (s, 3H), 2.20 (m, 1H), 1.84 (d, 2H, J = 12.8 Hz), 1.38 (m, 8H). Found m/z = 381.2 $(M+H)^+$.

*Synthesis of (3,5-dimethylbenzofuran-2-yl)(4-((3-isopropyl-1,2,4-oxadiazol-5-yl)methyl)piperidin-1-yl)methanone (CP4.29.16)*

In a 20 mL vial, 100.7 mg (0.529 mmol) of 3,5-dimethyl-1-benzofuran-2-carboxylic acid (Enamine Ltd., Kyiv, Ukraine; EN300-12940), 261.9 mg (0.689 mmol) HATU, 277 µL DIPEA (d=0.742 g/mL, 1.587 mmol) and 3 mL of dry dimethylformamide were added and allowed to mix with rapid stirring for 10 minutes. To the resulting mixture, 159.6 mg of 4-[[3-(propan-2-yl)-1,2,4-oxadiazol-5-yl]methyl]piperidine hydrochloride (0.649 mmol; Enamine Ltd., Kyiv, Ukraine; EN300-260977) were added and the reaction was left to stir overnight at room temperature. Product was purified in a similar manner as CP4.29.14 with a few differences. Following acid/base workup, the organic layer was dried down and dissolved in MeOH. A sample of the MeOH soluble solution was used for LC-MS analysis. Remaining product was dried down on a rotary evaporator, dissolved in 1-3 mL DCM, and purified using normal phase chromatography on a 12 g silica column. Fractions containing product were left in hood to evaporate. These fractions were later dissolved in hot methanol and a sample of each was run on LC-MS. Highest purity fractions were pooled in a 20 mL vial and dried on rotary evaporator. The product was subsequently dissolved in $CDCl_3$ and a 1H NMR was taken. The sample was recovered and $CDCl_3$ was evaporated in the hood. Remaining sample was dissolved in hot DMSO, allowed to cool to room temperature, and purified via reverse phase HPLC on a C18 column using the same gradient already described. Fractions containing product were pooled and dried on the Genevac. From this point, compound was dried as already described for CP4.29.14. Final compound weight was 53.0 mg corresponding to a yield of 26.2%. 1H NMR ($CDCl_3$, 400 MHz): δ 7.34 (m, 2H), 7.28 (residual $CHCl_3$), 7.18 (dd, 1H, J = 8.44 Hz, 1.32 Hz), 4.50 (br, 2H), 3.08 (septet, 1H, J = 7.0 Hz), 2.86 (d, 2H, J = 7.12 Hz), 2.20 (m, 1H), 1.85 (br, 2H), 1.42 (qd, 2H, J = 12.84 Hz, 4.12 Hz), 1.35 (d, 6H, J = 6.92 Hz). Found m/z = 382.2 $(M+H)^+$.

*Synthesis of 2-(4-((3-isopropyl-1,2,4-oxadiazol-5-yl)methyl)piperidine-1-carbonyl)-1H-indole-4-carbonitrile (CP4.29.17)*

In a 20 mL vial, 99.7 mg (0.535 mmol) of 4-cyano-1H-indole-2-carboxylic acid (Enamine Ltd., Kyiv, Ukraine; EN300-321079), 268.4 mg (0.706 mmol) HATU, 280 µL DIPEA (d=0.742 g/mL, 1.605 mmol) and 3 mL of dry dimethylformamide were added and allowed to mix with rapid stirring for 10 minutes. To the resulting mixture, 159.4 mg of 4-[[3-(propan-2-yl)-1,2,4-oxadiazol-5-yl]methyl]piperidine hydrochloride (0.649 mmol; Enamine Ltd., Kyiv, Ukraine; EN300-260977) were added and the reaction was left to stir overnight at room temperature. Product was purified in a similar way to CP4.29.16 with a few changes. Following purification via reverse phase HPLC, a contaminant from the instrument co-eluted with the product. The resulting product was re-run through the same reverse phase HPLC after cleaning the instrument, but contaminant was still present. Fractions were dried using the Genevac at a maximum temperature of 45°C. Remaining product was purified from the contaminant using normal phase chromatography with a 4 g silica column and a DCM/MeOH (up to 10%) linear gradient. Fractions were left to partially evaporate overnight in the hood and the following morning, fractions corresponding to single peak were pooled. Product was transferred to a clean pre-weighed vial and dried on a rotary evaporator. The resulting solid was dissolved in $CDCl_3$ and used for 1H NMR analysis. Sample was returned to the original vial and $CDCl_3$ removed by evaporation in the hood. Product was dissolved in DCM, transferred to 4 mL vial, and dried with a rotary evaporator. Finally, the vial was covered with a Kim wipe and left overnight in a vacuum desiccator. Final compound weight was 2.2 mg corresponding to a 1.1% yield. 1H NMR ($CDCl_3$, 400 MHz): δ 9.72 (s, 1H), 7.58 (d, 1H, J = 8.36 Hz), 7.45 (dd, 1H, J = 7.32 Hz, 0.68 Hz), 7.25 (t, 1H, J = 7.44 Hz), 7.19 (s, 2H), 6.87 (d, 1H, J = 1.28 Hz), 4.66 (d, 2H, J = 13.32 Hz), 3.42 (s, 0.6H), 3.01 (septet, 1H, J = 6.96 Hz), 2.81 (m, 3H), 2.19 (m, 1H), 1.87 (d, 2H, J = 11.36 Hz), 1.51 (br, 3H), 1.35 (qd, 3H, J = 8.36 Hz, 3.96 Hz), 1.25 (m, 11H). Found m/z = 378.2 $(M+H)^+$.

*Synthesis of 2-(4-((3-isopropyl-1,2,4-oxadiazol-5-yl)methyl)piperidine-1-carbonyl)-1H-indole-3-carbonitrile (CP4.29.18)*

In a 20 mL vial, 98.0 mg (0.526 mmol) of 3-cyano-1H-indole-2-carboxylic acid (Enamine Ltd., Kyiv, Ukraine; EN300-344668), 247.2 mg (0.650 mmol) HATU, 275 µL DIPEA (d=0.742 g/mL, 1.578 mmol) and 3 mL of dry dimethylformamide were added and allowed to mix with rapid stirring for 10 minutes. To the resulting mixture, 148.5 mg of 4-[[3-(propan-2-yl)-1,2,4-oxadiazol-5-yl]methyl]piperidine hydrochloride (0.604 mmol; Enamine Ltd., Kyiv, Ukraine; EN300-260977) were added and the reaction was left to stir overnight at room temperature. Product was purified in the same manner as described for CP4.29.17. Final compound weight was 3.6 mg corresponding to a 1.8% yield. 1H NMR ($CDCl_3$, 400 MHz): δ 10.26 (s, 1H), 7.68 (d, 1H, J = 8.12 Hz), 7.39 (d, 1H, 8.24 Hz), 7.30 (td,

1H, J = 7.04 Hz, 1.12 Hz), 7.23 (m, 1H), 7.19 (s, 1H), 4.43 (br, 2H), 3.17 (br, 2H), 3.01 (septet, 1H, J = 6.92 Hz), 2.80 (d, 2H, J = 7.12 Hz), 2.17 (m, 1H), 1.86 (br, 2H), 1.50 (m, 6H), 1.20 (m, 8H). Found m/z = 378.2 $(M+H)^+$.

*Synthesis of (4-cyclopropyl-1H-indol-2-yl)(4-((3-isopropyl-1,2,4-oxadiazol-5-yl)methyl)piperidin-1-yl)methanone (CP4.29.19)*

In a 20 mL vial, 94.6 mg (0.470 mmol) of 4-cyclopropyl-1H-indole-2-carboxylic acid (Enamine Ltd., Kyiv, Ukraine; EN300-323048), 219.7 mg (0.578 mmol) HATU, 246 µL DIPEA (d=0.742 g/mL, 1.41 mmol) and 3 mL of dry dimethylformamide were added and allowed to mix with rapid stirring for 10 minutes. To the resulting mixture, 142.9 mg of 4-[[3-(propan-2-yl)-1,2,4-oxadiazol-5-yl]methyl]piperidine hydrochloride (0.581 mmol; Enamine Ltd., Kyiv, Ukraine; EN300-260977) were added and the reaction was left to stir overnight at room temperature. Product was purified in the same manner as described for CP4.29.17. Final compound weight was 4.2 mg corresponding to a 2.3% yield. 1H NMR ($CDCl_3$, 400 MHz): δ 9.12 (s, 1H), 7.14 (m, 3H), 6.86 (d, 1H, J = 1.32 Hz), 6.64 (d, 1H, 7.0 Hz), 4.70 (d, 2H, J = 13.4 Hz), 3.01 (m, 3H), 2.80 (d, 2H, J = 7.12 Hz), 2.15 (m, 2H), 1.84 (br, 2H), 1.57 (br, 1H), 1.36 (qd, 2H, J = 12.6 Hz, 3.96 Hz), 1.20 (m, 8H), 0.93 (m, 2H), 0.75 (m, 2H). Found m/z = 393.2 $(M+H)^+$.

*Synthesis of (3-cyclopropyl-1H-indol-2-yl)(4-((3-isopropyl-1,2,4-oxadiazol-5-yl)methyl)piperidin-1-yl)methanone (CP4.29.20)*

In a 20 mL vial, 101.2 mg (0.503 mmol) of 3-cyclopropyl-1H-indole-2-carboxylic acid (Enamine Ltd., Kyiv, Ukraine; EN300-6764589), 237.3 mg (0.624 mmol) HATU, 263 µL DIPEA (d=0.742 g/mL, 1.509 mmol) and 3 mL of dry dimethylformamide were added and allowed to mix with rapid stirring for 10 minutes. To the resulting mixture, 152.5 mg of 4-[[3-(propan-2-yl)-1,2,4-oxadiazol-5-yl]methyl]piperidine hydrochloride (0.620 mmol; Enamine Ltd., Kyiv, Ukraine; EN300-260977) were added and the reaction was left to stir overnight at room temperature. Product was purified in the same manner as described for CP4.29.17. Final compound weight was 30.3 mg corresponding to a 15.3% yield.

1H NMR ($CDCl_3$, 400 MHz): δ 9.08 (s, 1H), 7.60 (d, 1H, J = 7.96 Hz), 7.26 (d, 1H, J = 8.2 Hz), 7.13 (td, 1H, J = 7.04 Hz, 1.0 Hz), 7.02 (t, 1H, J = 7.92 Hz), 4.42 (br, 2H), 3.39 (s, 1H), 2.99 (m, 3H), 2.74 (d, 2H, J = 7.08 Hz), 2.10 (m, 1H), 1.80 (m, 4H), 1.30 (m, 9H), 0.83 (m, 2H), 0.65 (m, 2H). Found m/z = 393.2 $(M+H)^+$.

*Synthesis of (4-((3-isopropyl-1,2,4-oxadiazol-5-yl)methyl)piperidin-1-yl)(3-phenyl-1H-indol-2-yl)methanone (CP4.29.21)*

In a 20 mL vial, 100.8 mg (0.425 mmol) of 3-phenyl-1H-indole-2-carboxylic acid (Enamine Ltd., Kyiv, Ukraine; EN300-7536218), 195.9 mg (0.515 mmol) HATU, 223 µL DIPEA (d=0.742 g/mL, 1.275 mmol) and 3 mL of dry dimethylformamide were added and allowed to mix with rapid stirring for 10 minutes. To the resulting mixture, 126.9 mg of 4-[[3-(propan-2-yl)-1,2,4-oxadiazol-5-yl]methyl]piperidine hydrochloride (0.516 mmol; Enamine Ltd., Kyiv, Ukraine; EN300-260977) were added and the reaction was left to stir overnight at room temperature. Product was purified in the same manner as described for CP4.29.17. Final compound weight was 10.3 mg corresponding to a 5.7% yield. 1H NMR ($CDCl_3$, 400 MHz): δ 9.14 (s, 1H), 7.56 (d, 1H, J = 2.2 Hz), 7.28 (m, 5H), 7.17 (tt, 1H, J = 6.52 Hz, 1.4 Hz), 7.10 (m, 1H), 7.07 (s, 1H), 6.97 (m, 1H), 4.00 (br, 2H), 2.85 (septet, 1H, J = 6.96 Hz), 2.39 (m, 4H), 1.69 (m, 1H), 1.56 (br, 2H), 1.11 (m, 8H), 0.00 (br, 1H). Found m/z = 429.2 $(M+H)^+$.

*Synthesis of (4-((3-isopropyl-1,2,4-oxadiazol-5-yl)methyl)piperidin-1-yl)(1H-pyrrolo[2,3-b]pyridin-2-yl)methanone (CP4.29.22)*

In a 20 mL vial, 97.0 mg (0.598 mmol) of 1H-pyrrolo[2,3-b]pyridine-2-carboxylic acid (Enamine Ltd., Kyiv, Ukraine; EN300-33541), 299.8 mg (0.788 mmol) HATU, 312 µL DIPEA (d=0.742 g/mL, 1.794 mmol) and 3 mL of dry dimethylformamide were added and allowed to mix with rapid stirring for 10 minutes. To the resulting mixture, 180.6 mg of 4-[[3-(propan-2-yl)-1,2,4-oxadiazol-5-yl]methyl]piperidine hydrochloride (0.735 mmol; Enamine Ltd., Kyiv, Ukraine; EN300-260977) were added and the reaction was left to stir overnight at room temperature. Product was purified in the same manner as described for CP4.29.17. Final compound weight was 7.7 mg corresponding to a 3.6% yield.

1H NMR ($CDCl_3$, 400 MHz): δ 8.43 (d, 1H, J = 4.0 Hz), 7.93 (dd, 1H, J = 7.96 Hz, 1.44 Hz), 7.19 (s, 1H), 7.08 (dd, 1H, J = 4.76 Hz, 3.2 Hz), 6.65 (s, 1H), 4.63 (d, 2H, J = 12.68 Hz), 3.42 (s, 2H), 3.01 (m, 3H), 2.80 (d, 2H, 7.08), 2.18 (m, 1H), 1.83 (d, 2H, J = 12.04 Hz), 1.25 (m, 9H). Found m/z = 354.2 $(M+H)^+$.

S9. Malhotra S, Andrianov GV, Karanicolas J. A Quantitative View of Protein Surface Druggability: Predicting Maximum Achievable Potency. (manuscript in preparation).

S10. Malhotra S, Karanicolas J. https://github.com/ShipraMalhotra/PocketDruggability. 2020.

S11. Malhotra S. Predicting the Most Tractable Protein Surfaces in the Human Proteome for Developing New Therapeutics. 2019;PhD thesis, University of Kansas(16583).

S12. Omega2. 2.5.1.4 ed. Santa Fe, NM: OpenEye Scientific Software.

S13. Hawkins PC, Nicholls A. Conformer generation with OMEGA: learning from the data set and the analysis of failures. *J Chem Inf Model*. 2012;52(11):2919-36.

S14. Hawkins PC, Skillman AG, Warren GL, Ellingson BA, Stahl MT. Conformer generation with OMEGA: algorithm and validation using high quality structures from the Protein Databank and Cambridge Structural Database. *J Chem Inf Model*. 2010;50(4):572-84.

S15. Hawkins PC, Skillman AG, Nicholls A. Comparison of shape-matching and docking as virtual screening tools. *J Med Chem*. 2007;50(1):74-82.

S16. ROCS. 3.2.1.4 ed. Santa Fe, NM: OpenEye Scientific Software.

S17. OEChem Toolkit. 2.1.3 ed. Santa Fe, NM: OpenEye Scientific Software; 2017.

S18. QUACPAC. 1.7.0.2 ed. Santa Fe, NM: OpenEye Scientific Software.

S19. Adeshina YO, Deeds EJ, Karanicolas J. Machine learning classification can reduce false positives in structure-based virtual screening. *Proc Natl Acad Sci U S A*. 2020;117(31):18477-88.

S20. REAL Space Navigator. Sankt Augustin, Germany: BioSolveIT GmbH; 2019.

S21. infiniSee. 3.2 Andromeda ed. Sankt Augustin, Germany: BioSolveIT GmbH; 2021.

S22. Tan S. A modular polycistronic expression system for overexpressing protein complexes in Escherichia coli. *Protein Expr Purif*. 2001;21(1):224-34.

S23. Davis B. Screening protein-small molecule interactions by NMR. *Methods Mol Biol*. 2013;1008:389-413.

S24. Duvaud S, Gabella C, Lisacek F, Stockinger H, Ioannidis V, Durinx C. Expasy, the Swiss Bioinformatics Resource Portal, as designed by its users. *Nucleic Acids Res*. 2021;49(W1):W216-W27.

S25. Calabrese DR, Connelly CM, Schneekloth JS, Jr. Ligand-observed NMR techniques to probe RNA-small molecule interactions. *Methods Enzymol*. 2019;623:131-49.

S26. Walpole S, Monaco S, Nepravishta R, Angulo J. STD NMR as a Technique for Ligand Screening and Structural Studies. *Methods Enzymol*. 2019;615:423-51.

S27. Huang R, Leung IKH. Protein-Small Molecule Interactions by WaterLOGSY. *Methods Enzymol*. 2019;615:477-500.

S28. Kao SH, Wang WL, Chen CY, Chang YL, Wu YY, Wang YT, Wang SP, Nesvizhskii AI, Chen YJ, Hong TM, Yang PC. Analysis of Protein Stability by the Cycloheximide Chase Assay. *Bio Protoc*. 2015;5(1).

### Supporting Tables

**Table S1:** Structures of compounds included in initial screen (CP1 to CP18).

| ID | Structure | SMILES |
|---|---|---|
| CP1 | 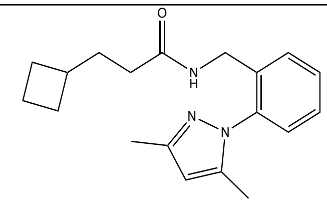 | CC1=NN(C2=C(CNC(CCC3CCC3)=O)C=CC=C2)C(C)=C1 |
| CP2 | 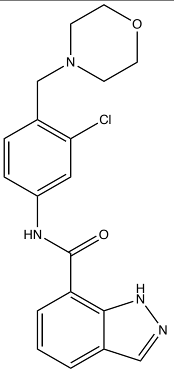 | ClC1=C(CN2CCOCC2)C=CC(NC(C3=C4NN=CC4=CC=C3)=O)=C1 |
| CP3 | 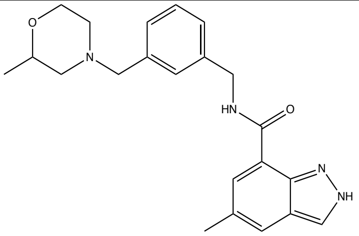 | CC1CN(CCO1)CC2=CC(CNC(C3=CC(C)=CC4=CNN=C34)=O)=CC=C2 |
| CP4 | 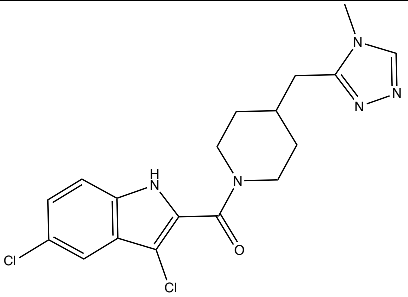 | CN1C=NN=C1CC2CCN(C(C3=C(Cl)C4=C(C=CC(Cl)=C4)N3)=O)CC2 |
| CP5 | 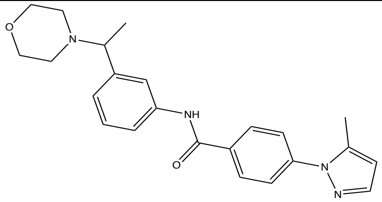 | CC(C1=CC(NC(C2=CC=C(N3N=CC=C3C)C=C2)=O)=CC=C1)N4CCOCC4 |

| | | |
|---|---|---|
| CP6 | 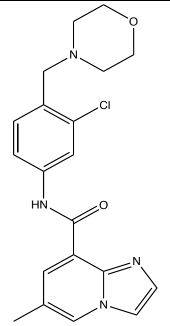 | CC1=CN2C=CN=C2C(C(NC3=CC(Cl)=C(CN4CCOCC4)C=C3)=O)=C1 |
| CP7 | 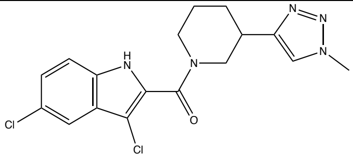 | CN1C=C(C2CCCN(C(C3=C(Cl)C4=C(C=CC(Cl)=C4)N3)=O)C2)N=N1 |
| CP8 | 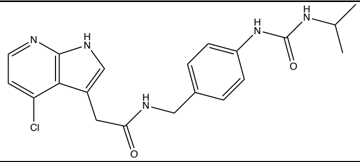 | CC(NC(NC1=CC=C(CNC(CC2=CNC3=C2C(Cl)=CC=N3)=O)C=C1)=O)C |
| CP9 | 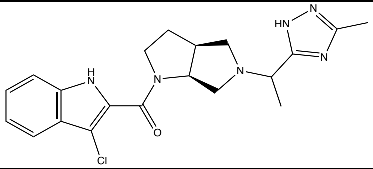 | CC(C1=NC(C)=NN1)N2C[C@@H]3CCN(C(C4=C(Cl)C5=C(C=CC=C5)N4)=O)[C@@H]3C2 |
| CP10 | 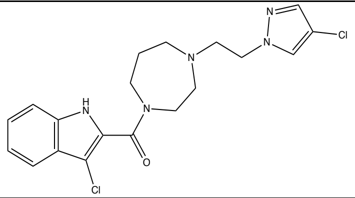 | ClC1=CN(CCN2CCCN(C(C3=C(Cl)C4=C(C=CC=C4)N3)=O)CC2)N=C1 |
| CP11 | 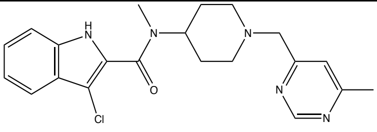 | CN(C1CCN(CC=2C=C(C)N=CN2)CC1)C(=O)C=3NC=4C=CC=CC4C3Cl |
| CP12 | 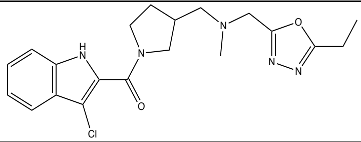 | CCC1=NN=C(O1)CN(CC2CCN(C(C3=C(Cl)C4=C(C=CC=C4)N3)=O)C2)C |
| CP13 | 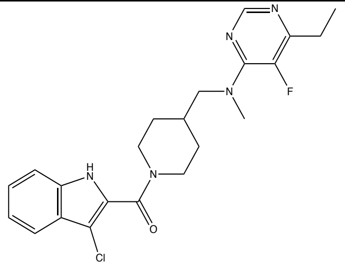 | CCC1=C(F)C(N(CC2CCN(C(C3=C(Cl)C4=C(C=CC=C4)N3)=O)CC2)C)=NC=N1 |

| | | |
|---|---|---|
| CP14 | 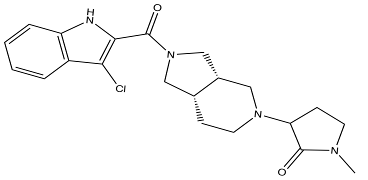 | CN1CCC(C1=O)N2CC[C@@H]3CN(C(C4=C(Cl)C5=C(C=CC=C5)N4)=O)C[C@@H]3C2 |
| CP15 | 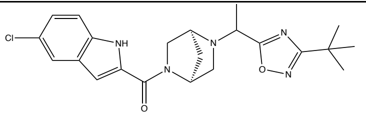 | CC(C1=NC(C(C)(C)C)=NO1)N2C[C@@H]3C[C@H]2CN3C(C4=CC5=C(N4)C=CC(Cl)=C5)=O |
| CP16 | 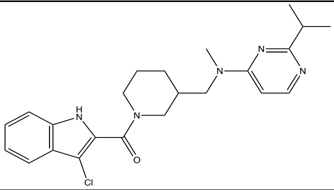 | CC(C1=NC(N(CC2CCCN(C(C3=C(Cl)C4=C(C=CC=C4)N3)=O)C2)C)=CC=N1)C |
| CP17 | 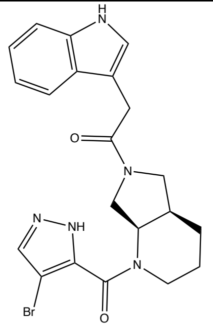 | BrC1=C(C(N2CCC[C@@H]3CN(C(CC4=CNC5=C4C=CC=C5)=O)C[C@@H]32)=O)NN=C1 |
| CP18 | 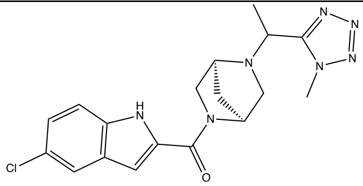 | CC(C1=NN=NN1C)N2C[C@@H]3C[C@H]2CN3C(C4=CC5=C(N4)C=CC(Cl)=C5)=O |

**Table S2:** Structures of CP4 derivatives included in second screen (CP4.1 to CP4.46).

| ID | Structure | SMILES |
|---|---|---|
| CP4.1 | 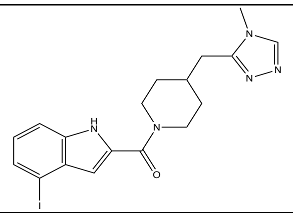 | CN1C=NN=C1CC2CCN(C(C3=CC4=C(N3)C=CC=C4I)=O)CC2 |
| CP4.2 | 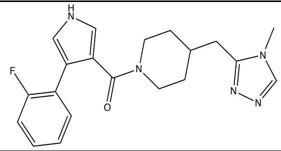 | O=C(C1=CNC=C1C2=C(F)C=CC=C2)N(CC3)CCC3CC4=NN=CN4C |
| CP4.3 | 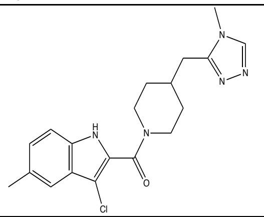 | CC1=CC2=C(NC(C(N3CCC(CC3)CC4=NN=CN4C)=O)=C2Cl)C=C1 |
| CP4.4 | 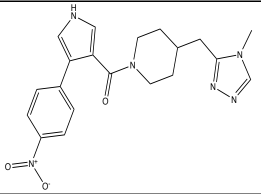 | O=C(C1=CNC=C1C2=CC=C([N+]([O-])=O)C=C2)N(CC3)CCC3CC4=NN=CN4C |
| CP4.5 | 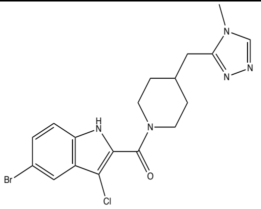 | CN1C=NN=C1CC2CCN(C(C3=C(Cl)C4=C(C=CC(Br)=C4)N3)=O)CC2 |
| CP4.6 | 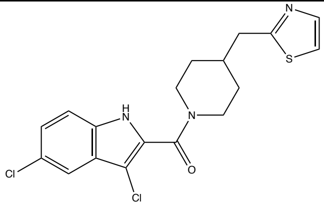 | ClC1=C(C(N2CCC(CC2)CC3=NC=CS3)=O)NC4=C1C=C(Cl)C=C4 |

| | | |
|---|---|---|
| CP4.7 | 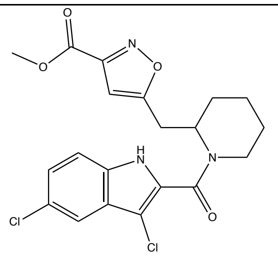 | COC(C1=NOC(CC2CCCCN2C(C3=C(Cl)C4=C(C=CC(Cl)=C4)N3)=O)=C1)=O |
| CP4.8 | 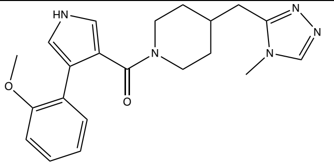 | COC1=C(C2=CNC=C2C(N3CCC(CC3)CC4=NN=CN4C)=O)C=CC=C1 |
| CP4.9 | 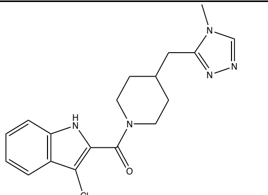 | CN1C=NN=C1CC2CCN(C(C3=C(Cl)C4=C(C=CC=C4)N3)=O)CC2 |
| CP4.10 | 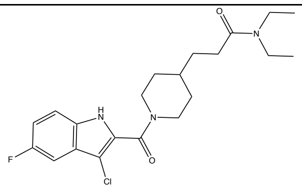 | CCN(C(CCC1CCN(C(C2=C(Cl)C3=C(C=CC(F)=C3)N2)=O)CC1)=O)CC |
| CP4.11 | 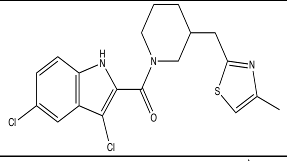 | CC1=CSC(CC2CCCN(C(C3=C(Cl)C4=C(C=CC(Cl)=C4)N3)=O)C2)=N1 |
| CP4.12 | 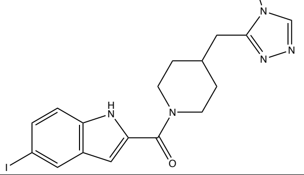 | CN1C=NN=C1CC2CCN(C(C3=CC4=C(N3)C=CC(I)=C4)=O)CC2 |
| CP4.13 | 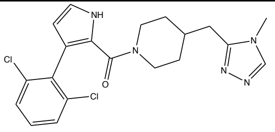 | CN1C=NN=C1CC2CCN(CC2)C(C3=C(C=CN3)C4=C(C=CC=C4Cl)Cl)=O |

| | | |
|---|---|---|
| CP4.14 | 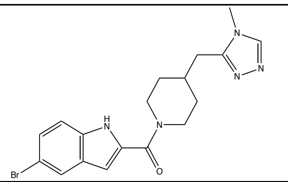 | CN1C=NN=C1CC2CCN(C(C3=CC4=C(N3)C=CC(Br)=C4)=O)CC2 |
| CP4.15 | 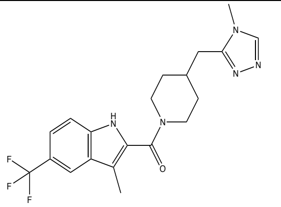 | CC1=C(C(N2CCC(CC2)CC3=NN=CN3C)=O)NC4=C1C=C(C(F)(F)F)C=C4 |
| CP4.16 | 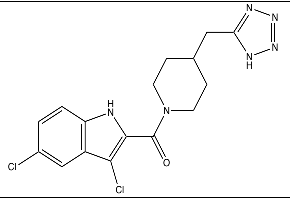 | ClC1=C(C(N2CCC(CC2)CC3=NN=NN3)=O)NC4=C1C=C(Cl)C=C4 |
| CP4.17 | 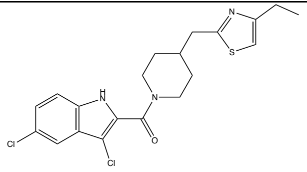 | CCC1=CSC(CC2CCN(C(C3=C(Cl)C4=C(C=CC(Cl)=C4)N3)=O)CC2)=N1 |
| CP4.18 | 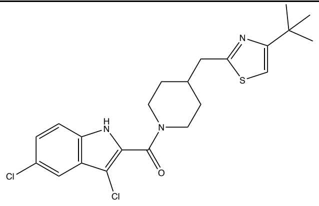 | CC(C)(C1=CSC(CC2CCN(C(C3=C(Cl)C4=C(C=CC(Cl)=C4)N3)=O)CC2)=N1)C |
| CP4.19 | 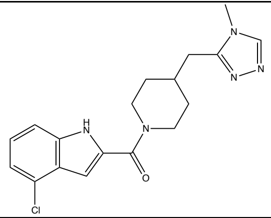 | CN1C=NN=C1CC2CCN(C(C3=CC4=C(N3)C=CC=C4Cl)=O)CC2 |
| CP4.20 | 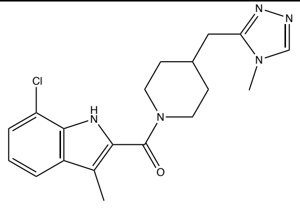 | CC1=C(C(N2CCC(CC2)CC3=NN=CN3C)=O)NC4=C1C=CC=C4Cl |
| CP4.21 | 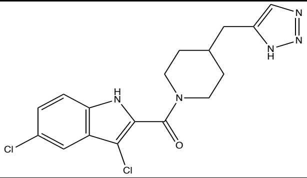 | ClC1=C(C(N2CCC(CC2)CC3=CN=NN3)=O)NC4=C1C=C(Cl)C=C4 |

| | | |
|---|---|---|
| CP4.22 | 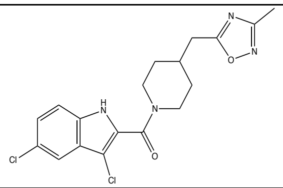 | CC1=NOC(CC2CCN(C(C3=C(Cl)C4=C(C=CC(Cl)=C4)N3)=O)CC2)=N1 |
| CP4.23 | 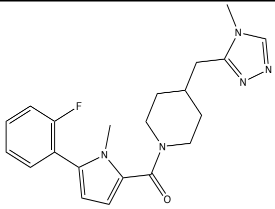 | CN1C=NN=C1CC2CCN(C(C3=CC=C(N3C)C4=C(F)C=CC=C4)=O)CC2 |
| CP4.24 | 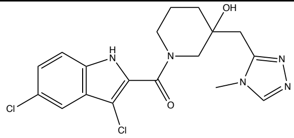 | CN1C=NN=C1CC2(CCCN(C(C3=C(Cl)C4=C(C=CC(Cl)=C4)N3)=O)C2)O |
| CP4.25 | 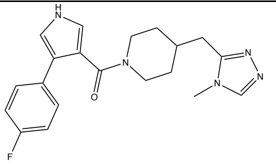 | CN1C=NN=C1CC2CCN(C(C3=CNC=C3C4=CC=C(F)C=C4)=O)CC2 |
| CP4.26 | 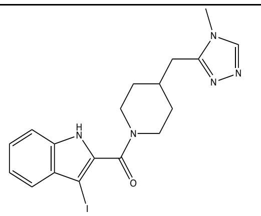 | CN1C=NN=C1CC2CCN(C(C3=C(I)C4=C(C=CC=C4)N3)=O)CC2 |
| CP4.27 | 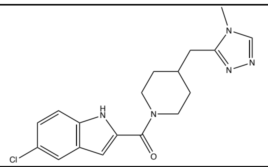 | CN1C=NN=C1CC2CCN(C(C3=CC4=C(N3)C=CC(Cl)=C4)=O)CC2 |
| CP4.28 | 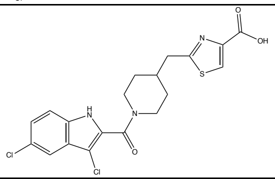 | OC(C1=CSC(CC2CCN(C(C3=C(Cl)C4=C(C=CC(Cl)=C4)N3)=O)CC2)=N1)=O |
| CP4.29 | 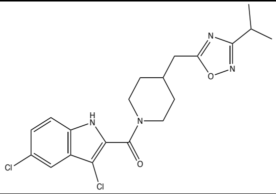 | CC(C1=NOC(CC2CCN(C(C3=C(Cl)C4=C(C=CC(Cl)=C4)N3)=O)CC2)=N1)C |
| CP4.30 | 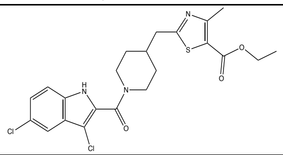 | CCOC(C1=C(C)N=C(CC2CCN(C(C3=C(Cl)C4=C(C=CC(Cl)=C4)N3)=O)CC2)S1)=O |

| | | |
|---|---|---|
| CP4.31 | 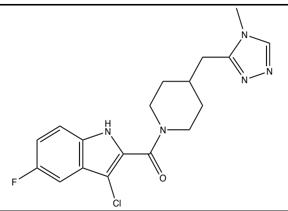 | CN1C=NN=C1CC2CCN(C(C3=C(Cl)C4=C(C=CC(F)=C4)N3)=O)CC2 |
| CP4.32 | 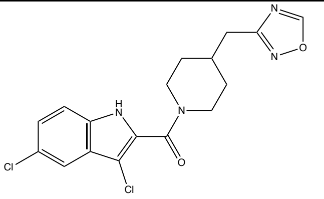 | ClC1=C(C(N2CCC(CC2)CC3=NOC=N3)=O)NC4=C1C=C(Cl)C=C4 |
| CP4.35 | 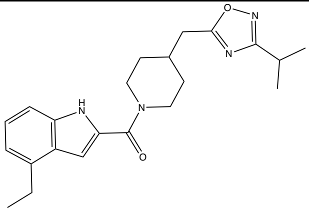 | O=C(C(N1)=CC2=C1C=CC=C2CC)N3CCC(CC4=NC(C(C)C)=NO4)CC3 |
| CP4.36 | 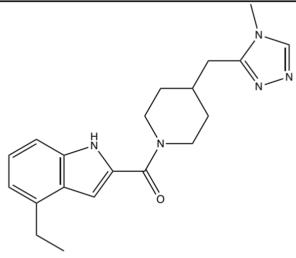 | O=C(C(N1)=CC2=C1C=CC=C2CC)N3CCC(CC4=NN=CN4C)CC3 |
| CP4.40 | 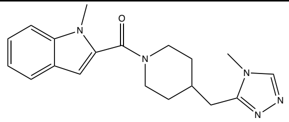 | O=C(N1CCC(CC2=NN=CN2C)CC1)C3=CC4=CC=CC=C4N3C |
| CP4.41 | 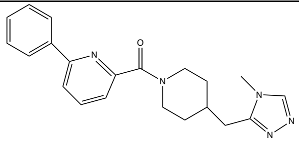 | O=C(C1=NC(C2=CC=CC=C2)=CC=C1)N3CCC(CC4=NN=CN4C)CC3 |
| CP4.42 | 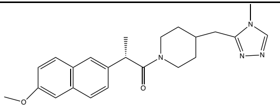 | O=C(N1CCC(CC2=NN=CN2C)CC1)[C@@H](C)C3=CC4=C(C=C(OC)C=C4)C=C3 |

| | | |
|---|---|---|
| CP4.43 | 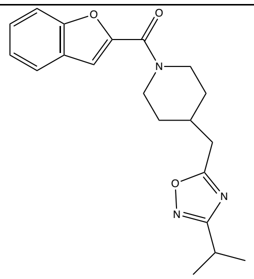 | O=C(N1CCC(CC2=NC(C(C)C)=NO2)CC1)C(O3)=C C4=C3C=CC=C4 |
| CP4.44 | 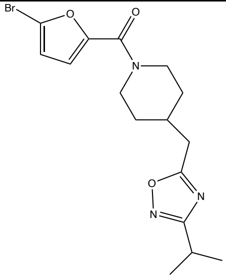 | BrC1=CC=C(C(N2CCC(CC3=NC(C(C)C)=NO3)CC2) =O)O1 |
| CP4.45 | 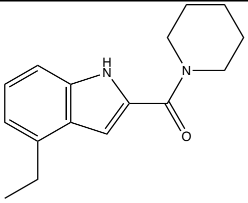 | O=C(C(N1)=CC2=C1C=CC=C2CC)N3CCCCC3 |
| CP4.46 | 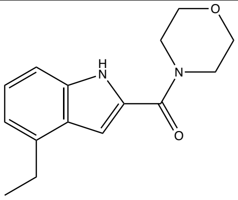 | O=C(C(N1)=CC2=C1C=CC=C2CC)N3CCOCC3 |

**Table S3:** Structures of CP4.29 derivatives included in third screen (CP4.29.1 to CP4.29.22).

| ID | Structure | SMILES |
|---|---|---|
| CP4.29.1 | 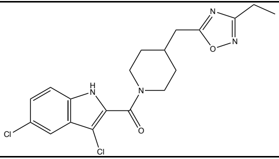 | O=C(C1=C(Cl)C2=C(C=CC(Cl)=C2)N1)N(CC3)CCC3CC4=NC(CC)=NO4 |
| CP4.29.3 | 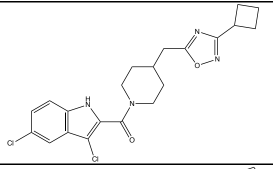 | O=C(C1=C(Cl)C2=C(C=CC(Cl)=C2)N1)N(CC3)CCC3CC4=NC(C5CCC5)=NO4 |
| CP4.29.4 | 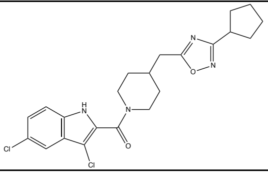 | O=C(C1=C(Cl)C2=C(C=CC(Cl)=C2)N1)N(CC3)CCC3CC4=NC(C5CCCC5)=NO4 |
| CP4.29.5 | 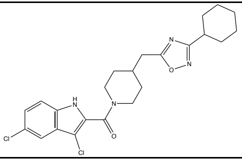 | O=C(C1=C(Cl)C2=C(C=CC(Cl)=C2)N1)N(CC3)CCC3CC4=NC(C5CCCCC5)=NO4 |
| CP4.29.6 | 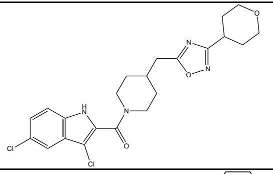 | O=C(C1=C(Cl)C2=C(C=CC(Cl)=C2)N1)N(CC3)CCC3CC4=NC(C5CCOCC5)=NO4 |
| CP4.29.9 | 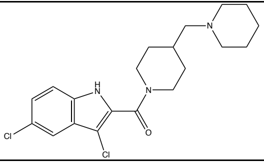 | O=C(N1CCC(CN2CCCCC2)CC1)C3=C(Cl)C4=C(C=CC(Cl)=C4)N3 |
| CP4.29.10 | 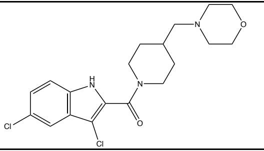 | O=C(N1CCC(CC1)CN2CCOCC2)C3=C(Cl)C4=C(C=CC(Cl)=C4)N3 |
| CP4.29.11 | 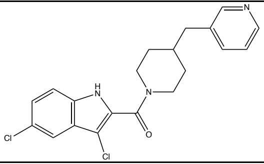 | O=C(N1CCC(CC2=CC=CN=C2)CC1)C3=C(Cl)C4=C(C=CC(Cl)=C4)N3 |
| CP4.29.12 | 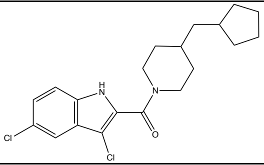 | O=C(N1CCC(CC2CCCC2)CC1)C3=C(Cl)C4=C(C=CC(Cl)=C4)N3 |

| | | |
|---|---|---|
| CP4.29.13 | 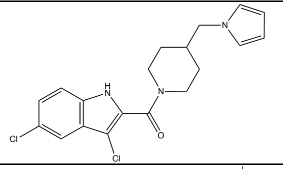 | O=C(N(CC1)CCC1CN2C=CC=C2)C3=C(Cl)C4=C(C=CC(Cl)=C4)N3 |
| CP4.29.14 | 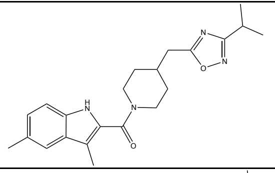 | O=C(N1CCC(CC2=NC(C(C)C)=NO2)CC1)C3=C(C)C4=CC(C)=CC=C4N3 |
| CP4.29.16 | 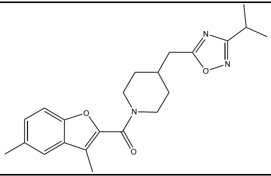 | O=C(N1CCC(CC2=NC(C(C)C)=NO2)CC1)C3=C(C)C4=CC(C)=CC=C4O3 |
| CP4.29.17 | 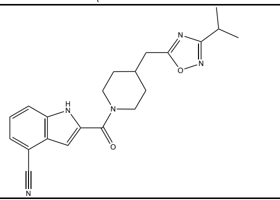 | O=C(N1CCC(CC2=NC(C(C)C)=NO2)CC1)C3=CC4=C(C=CC=C4C#N)N3 |
| CP4.29.18 | 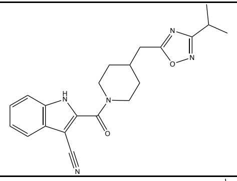 | N#CC1=C(C(N2CCC(CC3=NC(C(C)C)=NO3)CC2)=O)NC4=C1C=CC=C4 |
| CP4.29.19 | 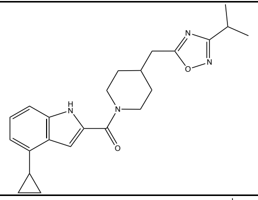 | O=C(N1CCC(CC2=NC(C(C)C)=NO2)CC1)C3=CC4=C(C=CC=C4C5CC5)N3 |
| CP4.29.20 | 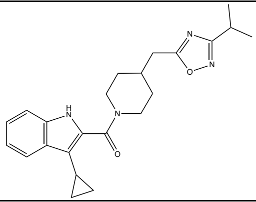 | O=C(N1CCC(CC2=NC(C(C)C)=NO2)CC1)C(NC3=C4C=CC=C3)=C4C5CC5 |
| CP4.29.21 | 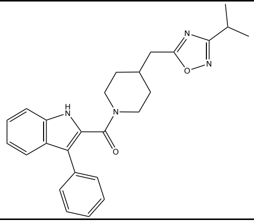 | O=C(N1CCC(CC2=NC(C(C)C)=NO2)CC1)C3=C(C4=CC=CC=C4)C(C=CC=C5)=C5N3 |
| CP4.29.22 | 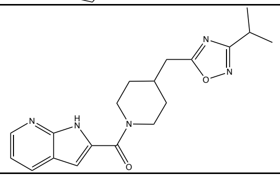 | O=C(N1CCC(CC2=NC(C(C)C)=NO2)CC1)C3=CC4=CC=CN=C4N3 |

### Supporting Figures

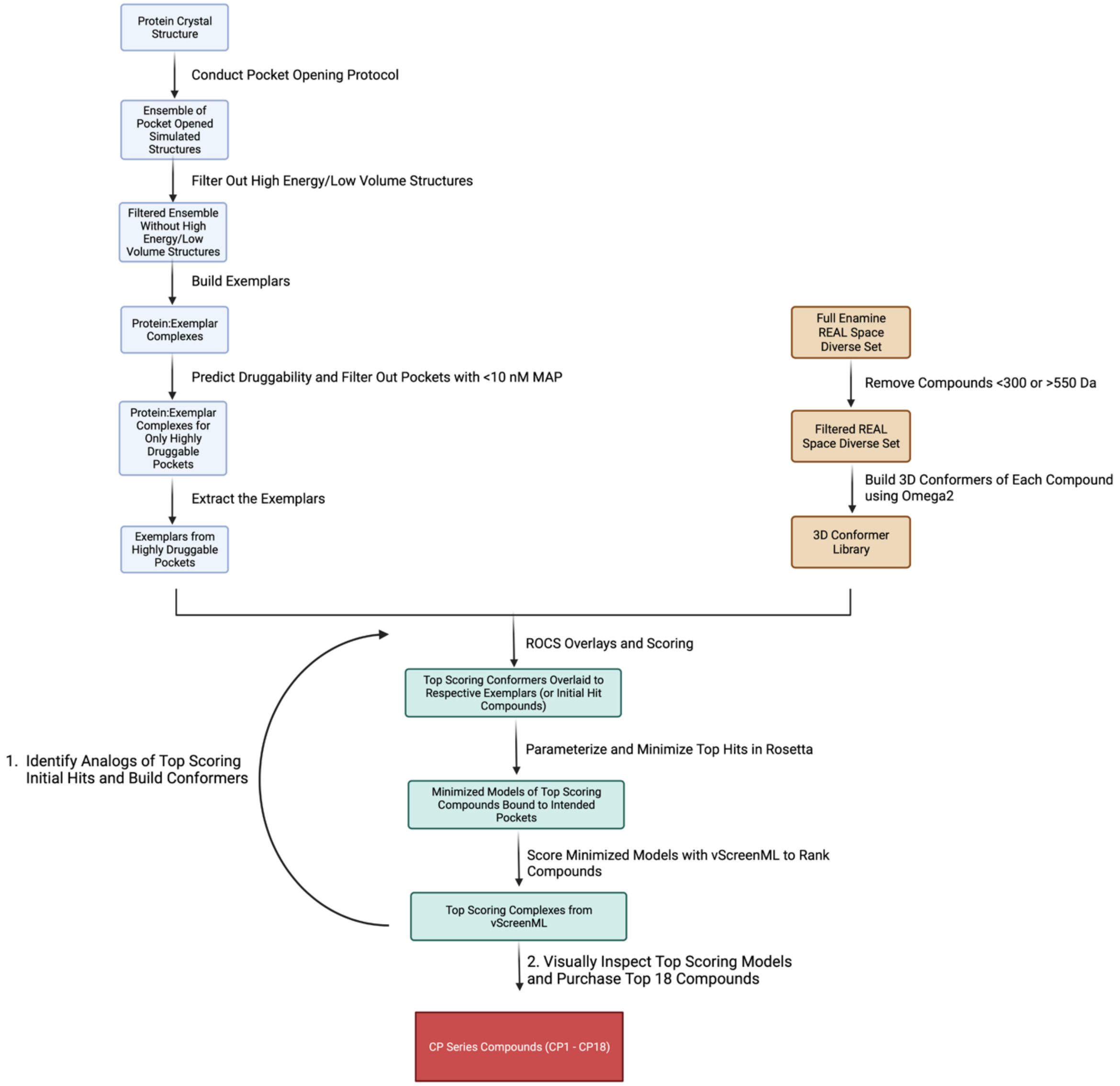


**Figure S1:** Summary of computational workflow used to identify initial hit compounds CP1 to CP18.

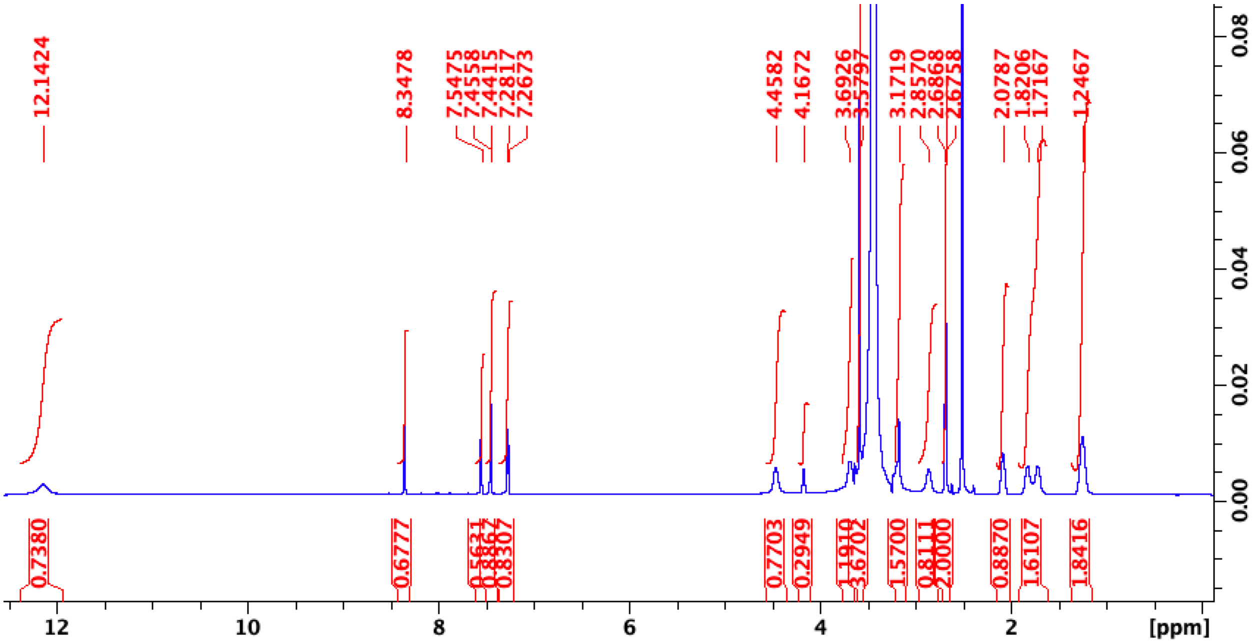


**Figure S2:** $^{1}$H NMR of CP4 in DMSO-d6 at 295K obtained on Bruker 600 MHz NMR.

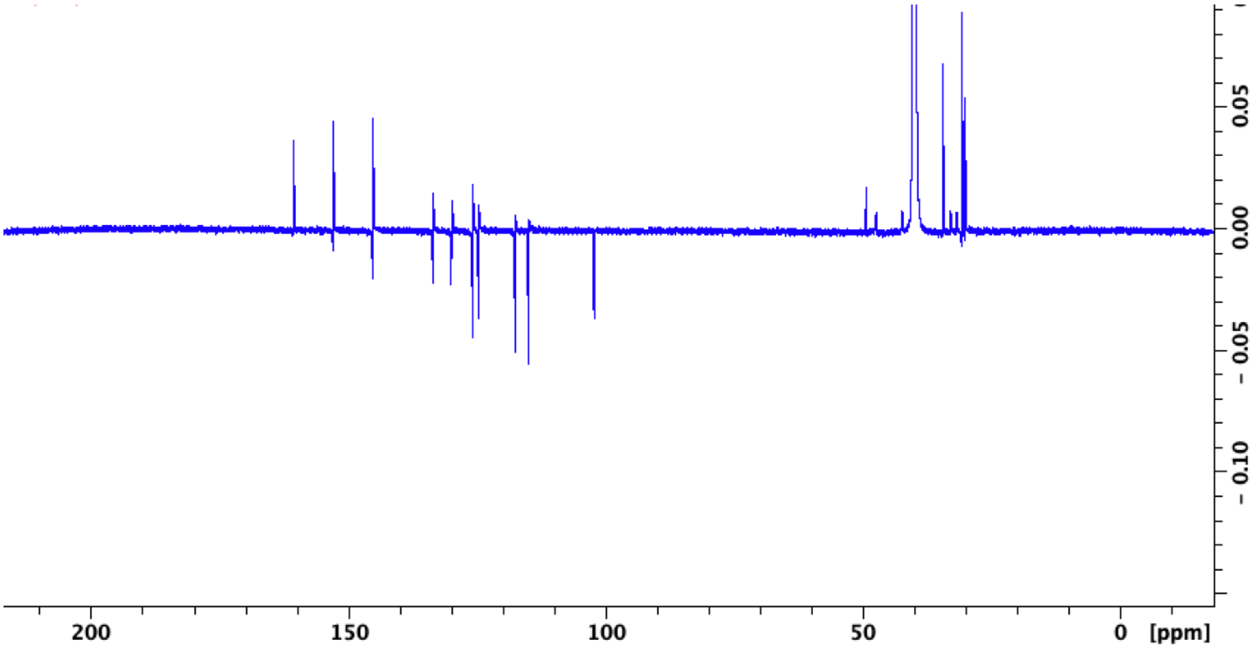


**Figure S3:** $^{13}$C NMR of CP4 in DMSO-d6 at 295K obtained on Bruker 600 MHz NMR.

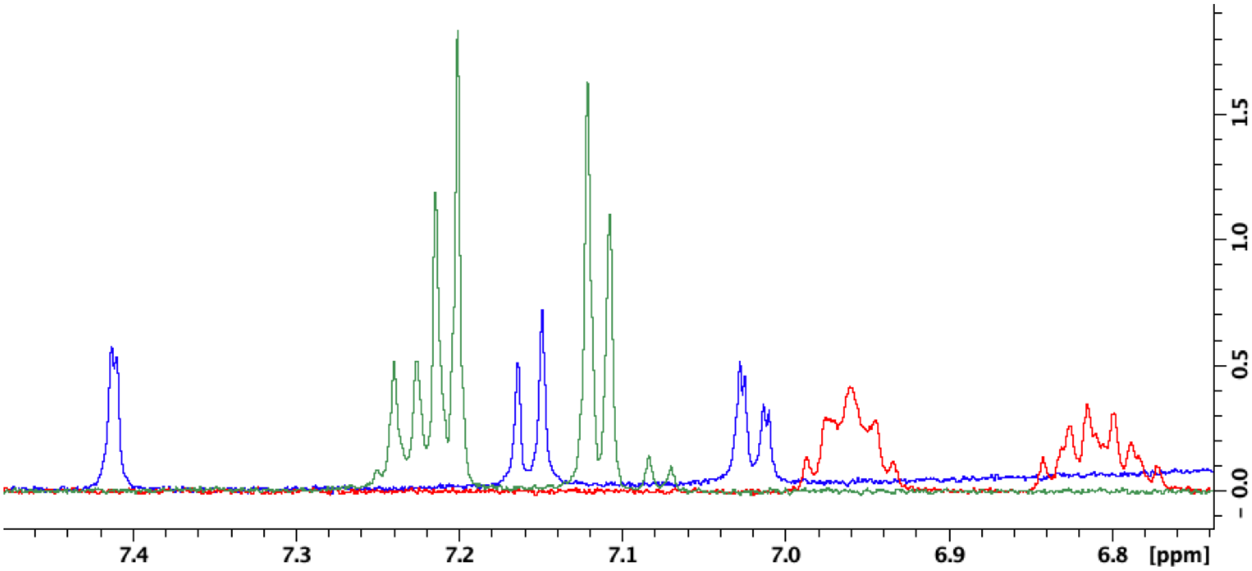


**Figure S4:** $^{1}$H NMR with water suppression (ZGESGP) of 100 μM CP4 (*blue*), 100 μM sitagliptin (*red*), or 100 μM VH298 (*green*) in 50 mM sodium phosphate buffer, 10% $D_2O$, pH=6.95, 1% DMSO-d6.

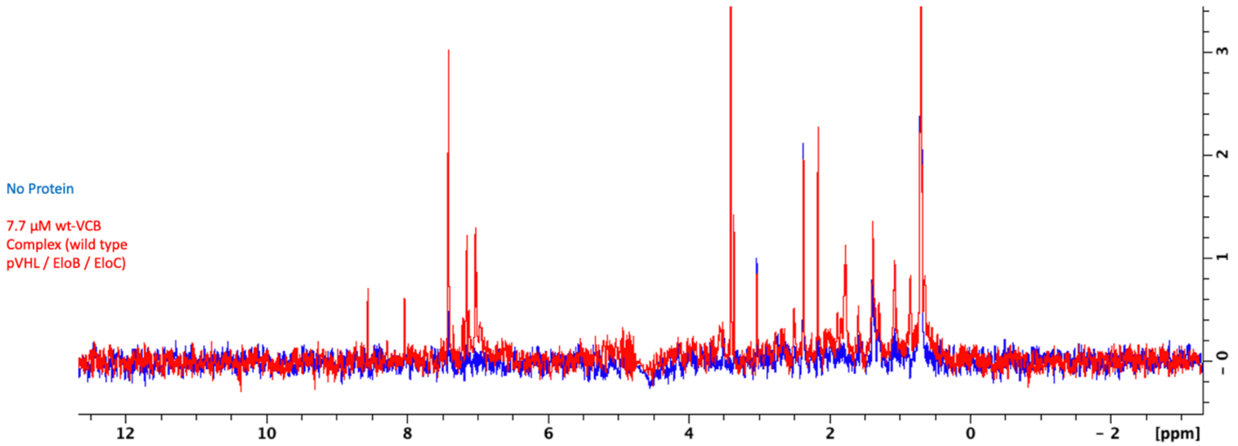


**Figure S5:** Full STD NMR spectrum for 100 μM CP4 / 100 μM Sitagliptin / 100 μM VH298 / 1% DMSO-d6 in the absence (*blue*) and presence (*red*) of 7.7 μM VCB protein complex.

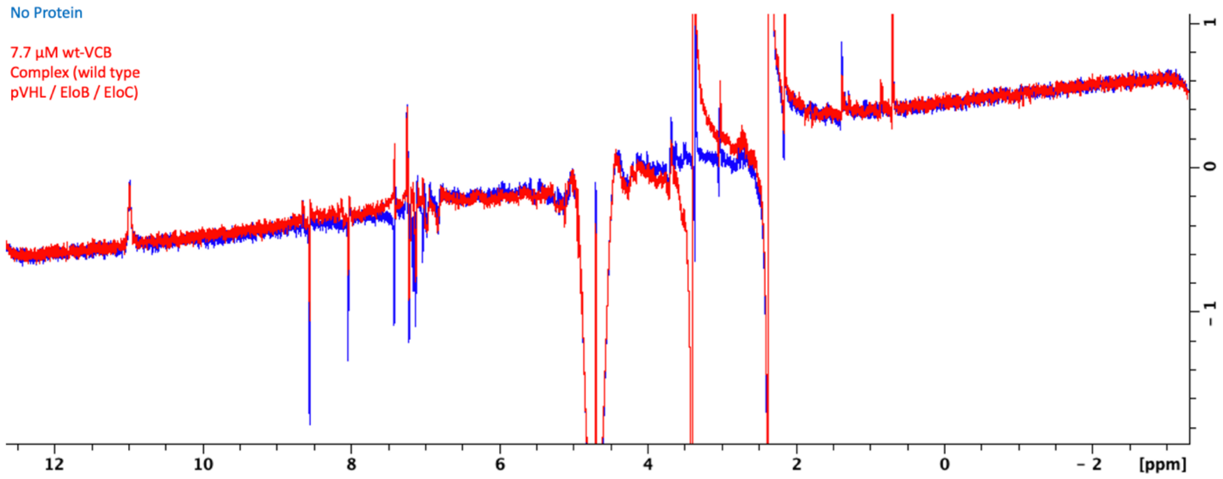


**Figure S6:** Full WaterLOGSY NMR spectrum for 100 µM CP4 / 100 µM Sitagliptin / 100 µM VH298 / 1% DMSO-d6 in the absence (*blue*) and presence (*red*) of 7.7 µM VCB protein complex.

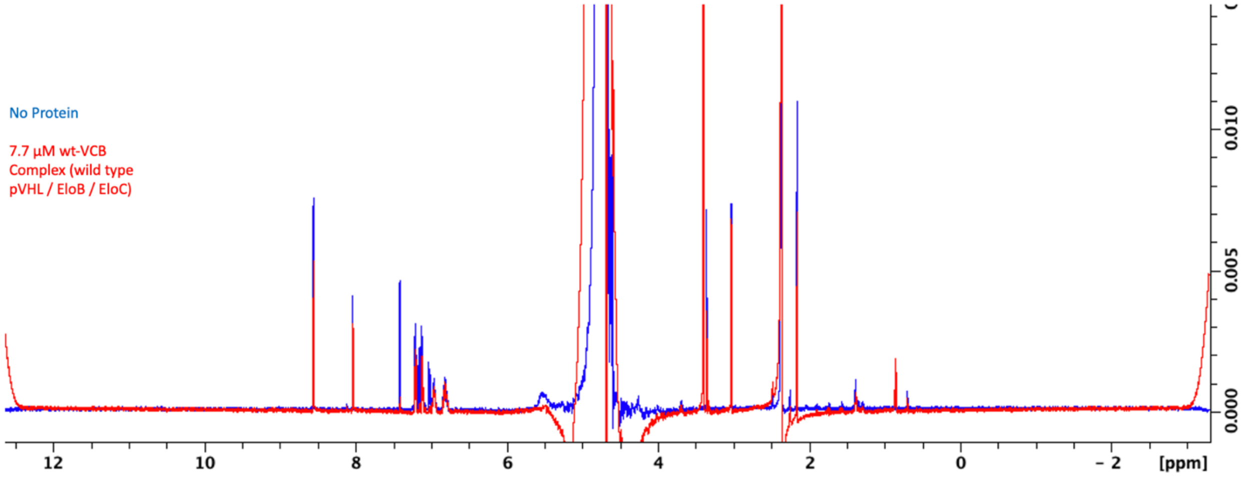


**Figure S7:** Full CPMG NMR spectrum for 100 µM CP4 / 100 µM Sitagliptin / 100 µM VH298 / 1% DMSO-d6 in the absence (*blue*) and presence (*red*) of 7.7 µM VCB protein complex.

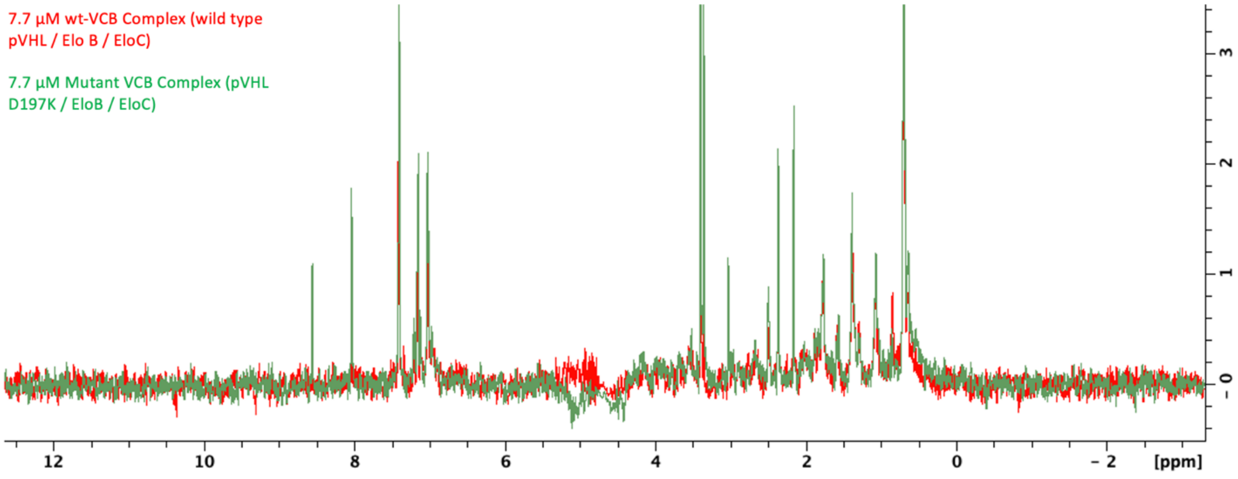


**Figure S8:** Full STD NMR spectrum for 100 µM CP4 / 100 µM Sitagliptin / 100 µM VH298 / 1% DMSO-d6 in the presence of 7.7 µM WT VCB protein complex (*red*) or 7.7 µM $V_{D197K}CB$ (*green*).

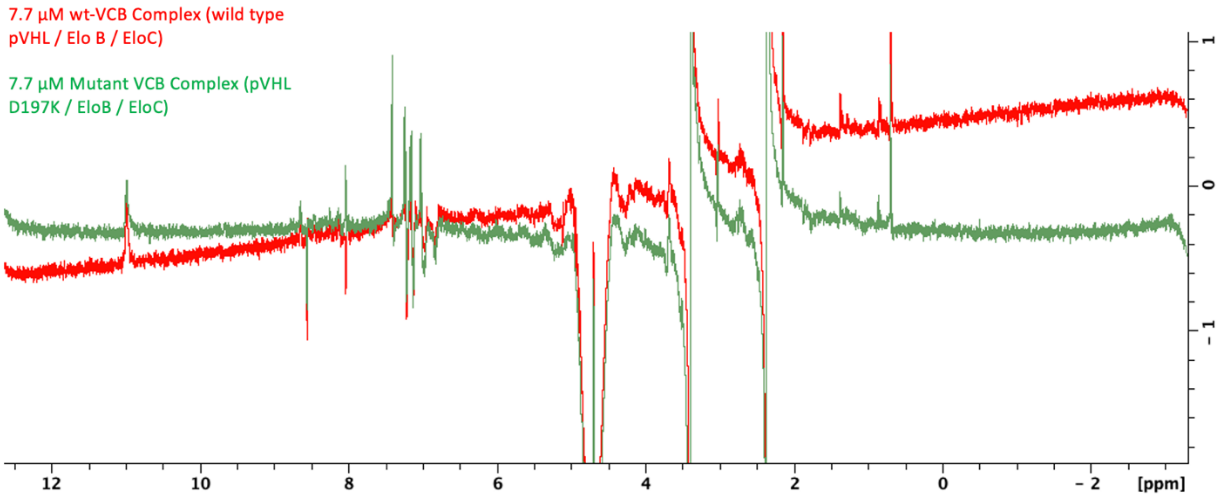


**Figure S9:** Full WaterLOGSY NMR spectrum for 100 µM CP4 / 100 µM Sitagliptin / 100 µM VH298 / 1% DMSO-d6 in the presence of 7.7 µM WT VCB protein complex (*red*) or 7.7 µM $V_{D197K}CB$ (*green*).

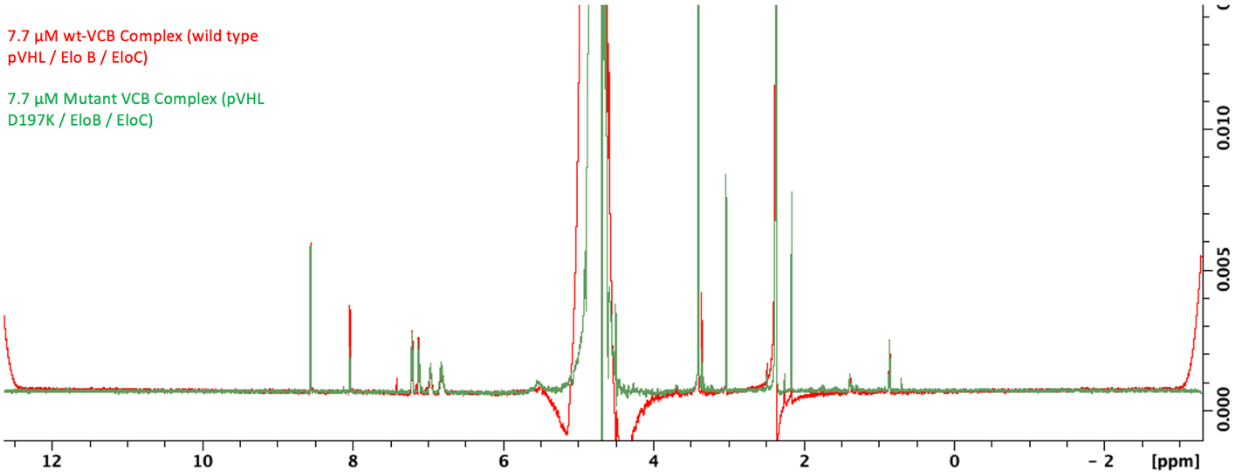


**Figure S10:** Full CPMG NMR spectrum for 100 µM CP4 / 100 µM Sitagliptin / 100 µM VH298 / 1% DMSO-d6 in the presence of 7.7 µM WT VCB protein complex (*red*) or 7.7 µM $V_{D197K}CB$ (*green*).

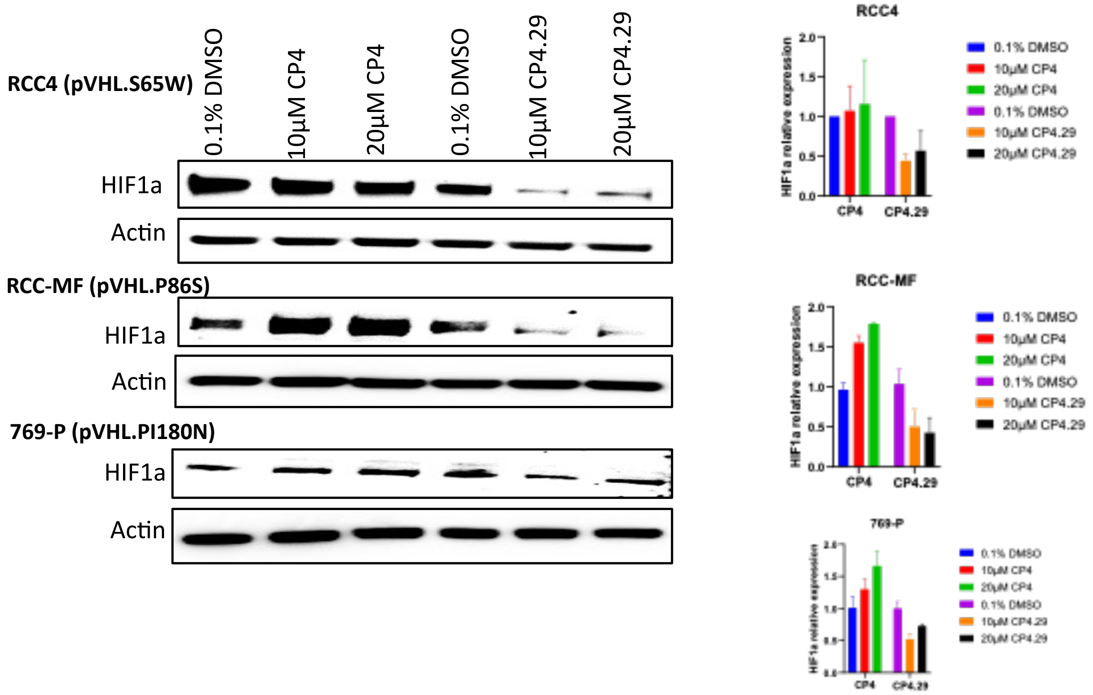


**Figure S11:** Treatment with CP4.29 (but not CP4) led to depletion of HIF-1α in RCC4, RCC-MF, and 769-P cells.

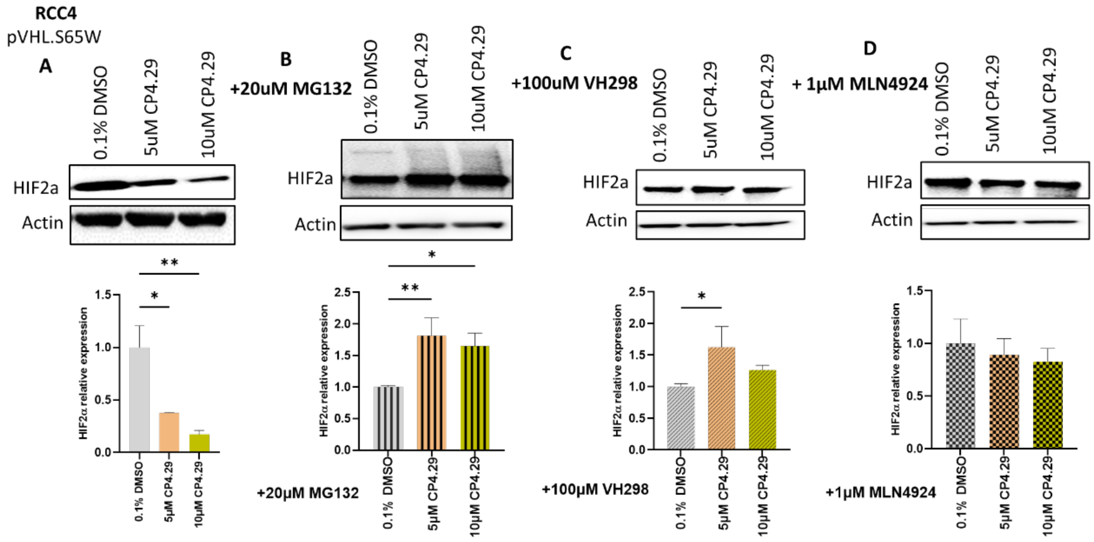


**Figure S12:** In RCC4 cells, **(A)** treatment with CP4.29 reduced the cellular abundance of HIF-2α. However, the activity of CP4.29 on HIF-2α could be reversed by co-treatment with either **(B)** proteasome inhibitor MG132, **(C)** VHL inhibitor VH298, or **(D)** neddylation inhibitor MLN4924.

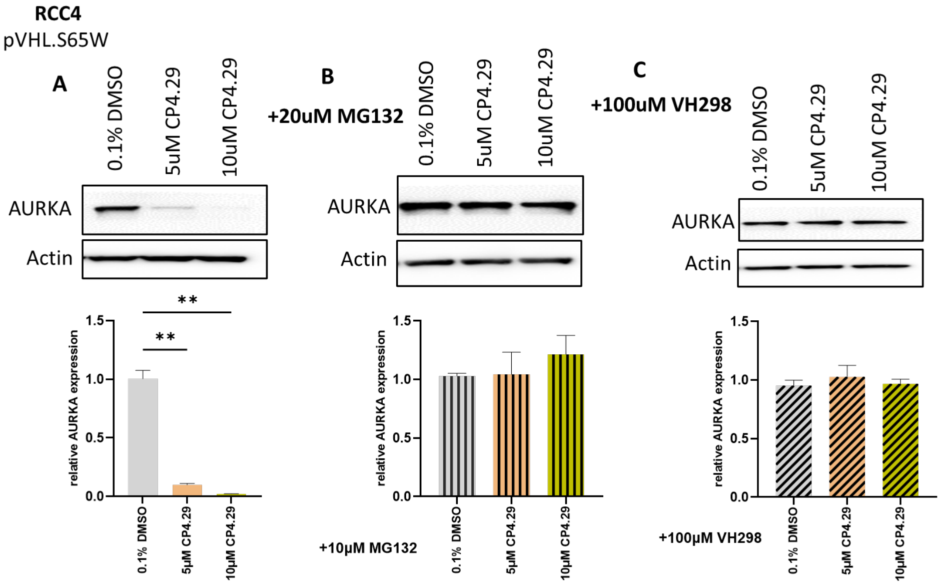


**Figure S13:** In RCC4 cells, **(A)** treatment with CP4.29 reduced the cellular abundance of Aurora A. However, the activity of CP4.29 on Aurora A could be reversed by co-treatment with **(B)** proteasome inhibitor MG132, or **(C)** VHL inhibitor VH298.

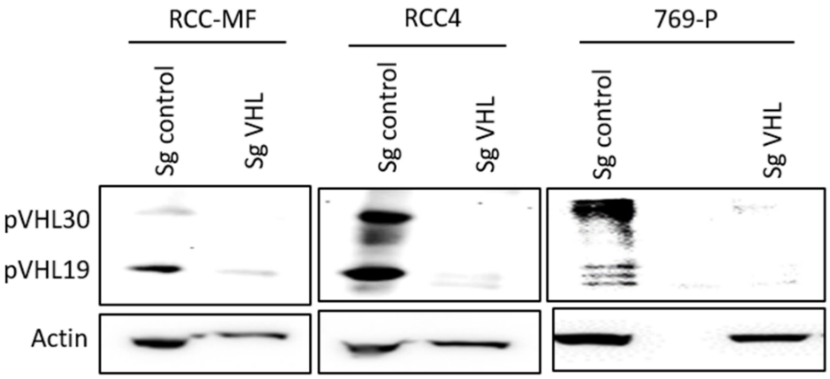


**Figure S14.** Parental cell lines RCC-MF, RCC4, and 769-P were each transfected with plasmids of pLenti-Cas9-GFP and were either provided with control sg-RNA or with sg-RNA for *VHL*. The loss of pVHL expression each *VHL* KO cell line was confirmed by Western blot.

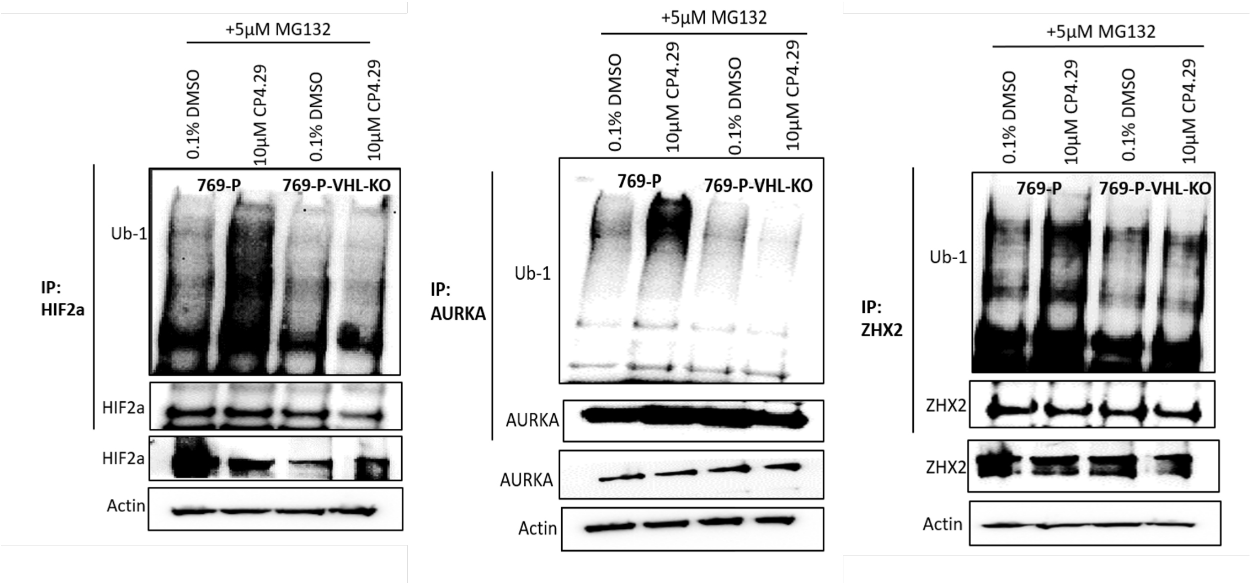


**Figure S15:** Immunoprecipitation (under non-denaturing conditions) of HIF-2α (*left*), AURKA (*middle*), or ZHX2 (*right*) followed by Western blot analysis for Ub-1. 769-P cells (or isogenic 769-P *VHL* KO cells) were treated with 20 μM proteasome inhibitor MG132, and with 10 μM CP4.29 for 2 hours in the HIF-2α experiment or for 24 hours in the AURKA and ZHX2 experiments.

| Alias | Solubility | Mouse Microsome Stability | | | Human Microsome Stability | | | Mouse plasma protein binding | | Mouse liver microsome partitioning | | hCYP 3A4 | hCYP2D6 | hCYP2C9 |
|---|---|---|---|---|---|---|---|---|---|---|---|---|---|---|
| | PBS (7.4) | t1/2 | Clint | w/o NADPH | t1/2 | Clint | w/o NADPH | Bound | Stable 20 hrs 37 C | Bound | Stable 20 hrs 37 C | IC50 | IC50 | IC50 |
| | uM | min | uL/min/ mg | % Stable | min | uL/min/ mg | % Stable | % | % | % | % | nM | nM | nM |
| CP4.29 | 2.5 | 15.6 | 88.6 | 78 | 26.7 | 51.9 | 89 | 99.1 | 107 | 97.9 | 131 | 2880 | > 10000 | > 10000 |

**Figure S16:** First-tier ADME characterization of CP4.29 indicates that further optimization is warranted prior to advancement into *in vivo* studies.

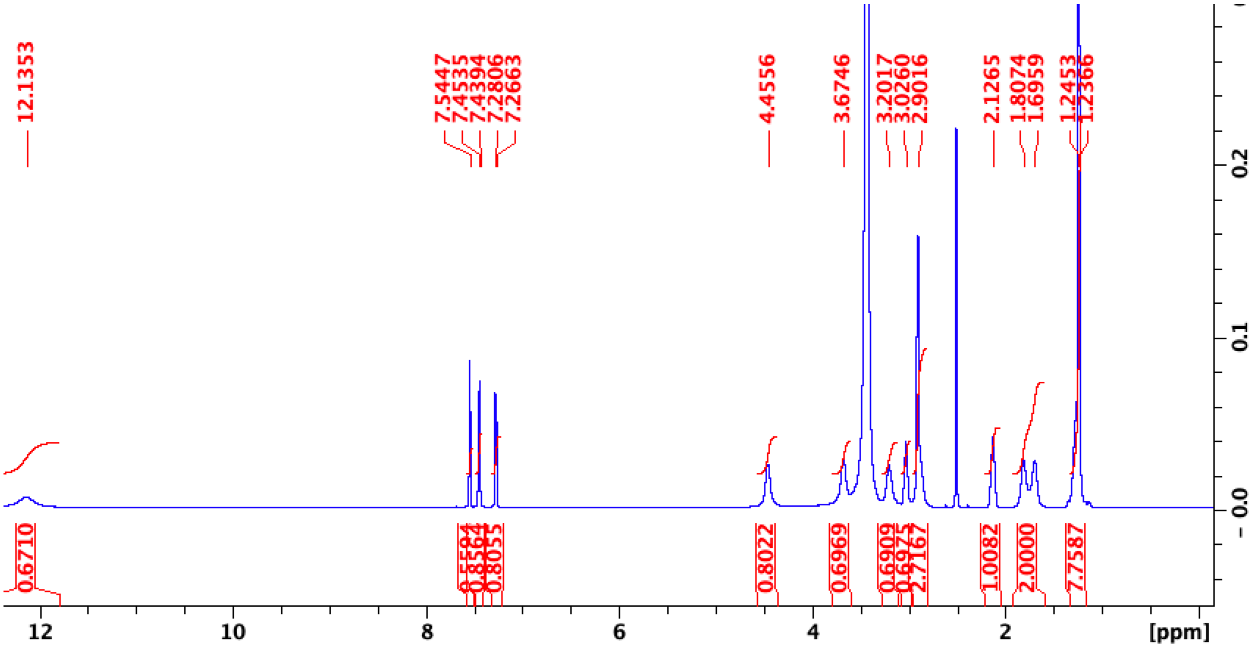


**Figure S17:** $^{1}$H NMR of CP4.29 in DMSO-d6 at 295K obtained on Bruker 600 MHz NMR.

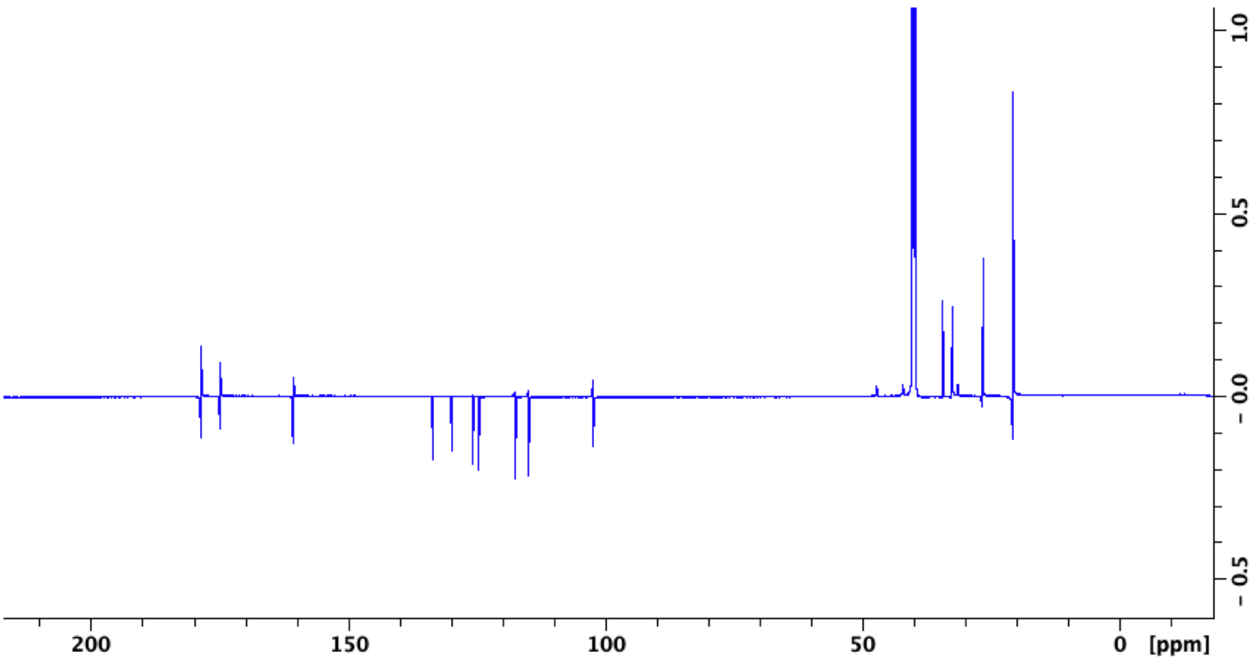


**Figure S18:** $^{13}$C NMR of CP4.29 in DMSO-d6 at 295K obtained on Bruker 600 MHz NMR.
